# Probing the role of sequential sampling and integration in decisions about protracted, noiseless stimuli

**DOI:** 10.1101/2025.10.22.684029

**Authors:** Hadiseh Hajimohammadi, Kieran S Mohr, Redmond G O’Connell, Simon P Kelly

## Abstract

Perceptual decision behaviour is known to be well-captured by models based on sequential sampling and temporal integration, but doubts have been raised about the generality and identifiability of these operations. Here we used neurally-constrained modelling to probe their role and temporal extent in the uncertain case of perceptual judgments about long-duration stimuli with no physical noise but weak evidence. We found that accuracy steadily improved across four covertly-manipulated evidence durations, indicating protracted sampling, but these delayed behavioural reports alone were insufficient to establish the operation of integration or of a decision-terminating bound. We then elaborated the models to prescribe how they would generate decision variable signals as well as choices, and fit them additionally to the average dynamics of a centroparietal electroencephalographic signal (‘CPP’) that traces decision formation. This established the setting of a bound and ruled out some non-integration mechanisms. However, one extrema detection model, which evokes a stereotyped signal ‘flagging’ the first bound-exceeding sample, rivalled the integration model in reproducing the evidence-dependent buildup dynamics of the CPP, alongside behavioural data. Moreover, the two models captured different features of neural motor preparation signals but neither captured all of them. We discuss the implications for the generality of integration and the technical challenges of neurally-informed modelling given limited behavioural data.

## Introduction

In the most prominent models of perceptual decision-making, evidence is sequentially sampled and integrated over time into a decision variable that can be subjected to a criterion to generate a choice (Gold & Shadlen, 2007). These models provide an intuitive means to achieve optimal performance despite perceptual noise and provide excellent fits to behaviour on a wide range of tasks (Ratcliff et al., 2016). Perfect integration over long timescales (tens of seconds) has been demonstrated using expanded judgment tasks where evidence is provided in just a few discrete, highly stochastic tokens (Hyafil et al., 2023; Waskom & Kiani, 2018), and extensive behavioural and neurophysiological work points to a key role of temporal integration in speeded judgments of continuously-presented, noisy stimuli like dot-motion (Gold & Shadlen 2007; Steinemann et al 2024). However, outside of these experimentally-tractable, stochastic and/or noisy task cases, an extended window of temporal integration is less clearly motivated theoretically (Uchida et al., 2006; Thura et al 2012) and can be more difficult to identify due to a lack of constraints (O’Connell et al 2018). Here we examine the case of judging a subtle feature of a long-duration, continuously-presented stimulus with no physical noise, to guide a later action choice. This case is of interest theoretically because there is no external noise - only internal neural noise - to motivate integration, and methodologically because decision completion times are not available to guide model comparison.

Even in the case of noisy stimuli, the temporal extent of integration is uncertain. In some cases sensory information is found to be strongly choice-predictive only over a restricted time range (e.g. Ludwig et al., 2005), and across experimenter-controlled, variable stimulus durations, improvements in accuracy are distinctly less than would be predicted if decision makers utilised all evidence through continued sampling, suggesting premature termination (Kiani et al., 2008). Such early decision termination is also expressed in neurophysiological decision signals reaching their peaks long before the end of noisy stimulus durations of >1 sec (Twomey et al., 2016; McCone et al 2026). It is not clear whether this reflects a general tendency to time-limit integration due to capacity limits or effort-saving, or the strategic setting of a bound that produces early terminations because the prevalence of strong (Twomey et al 2016) or short (Kiani et al 2008) stimuli meant that there was relatively little to be gained by extended sampling.

Here, we sought to assess the temporal extent of sampling, and the operation of integration in particular, in a task in which participants viewed a pair of noise-free gratings for a fixed 1.6 seconds before indicating which of them had higher contrast. While the lack of noise and long duration of the stimuli might obviate a strategy of integrating throughout, we made the vast majority of trials very difficult to confer a potential benefit of protracted integration. To probe the temporal extent of sampling behaviourally, while avoiding the above possibility of strategic bound-setting due to an expectation of short stimulus durations, we covertly manipulated evidence duration within the 1.6-sec stimulus window by seamlessly stepping the small contrast difference back to zero at a variable time, without telling the participants.

A key technical challenge that limits the ability to identify the operation of integration through behavioural modelling is the problem of model mimicry, where data generated from one mechanism can be quantitatively explained equally well by another mechanism (Evans & Wagenmakers, 2020; Khodadadi & Townsend, 2015; Trueblood et al., 2021). For example, even for dot-motion discrimination, non-integration strategies such as extrema detection have been shown to provide competitive fits to key behavioural data patterns classically associated with integration, including improvements in accuracy with duration and reduction of reaction time (RT) and error rate with increasing evidence strength (Stine et al., 2020). An emerging approach aimed at overcoming such mimicry issues is to further constrain the models with neurophysiological signals that reflect evidence-dependent decision formation dynamics (Hanks et al 2014; Purcell & Palmeri, 2017; O’Connell et al., 2018). Such decision signals have been reported in multiple brain areas of multiple species, most commonly in neural signals associated with preparing the decision-reporting actions (Gold & Shadlen, 2001; Hanks & Summerfield, 2017; O’Connell & Kelly 2021). In human electrophysiology, evidence-dependent buildup dynamics are observed not only in signatures of motor preparation such as decreases in Mu/Beta (8-30 Hz) amplitude contralateral to the upcoming movement (Pfurtscheller & Da Silva, 1999; Donner et al 2009; de Lange et al 2013; Murphy et al 2016), but also in a distinct, motor-independent Centro-Parietal Positivity (CPP) in the event-related potential (ERP). The CPP onsets at a time locked to evidence onset and builds at a rate that scales with evidence strength (Kelly & O’Connell 2013), dynamically tracking perturbations in the evidence, even in the absence of motor requirements (O’Connell et al., 2012). When decisions require immediate responses, the CPP peaks and falls at a time closely aligned to single-trial RT, with an amplitude that co-varies with decision accuracy (Kelly et al., 2021; Steinemann et al., 2018), and when responses are deferred until after a protracted stimulus, the CPP’s peak and fall during stimulus viewing marks early decision termination (Twomey et al. 2016; Pares-Pujolras et al., 2025; McCone et al., 2026). We thus used CPP dynamics alongside behavioural choice accuracy across evidence durations to examine the temporal extent of sampling in the present task. While much of the research to date has focused on the CPP’s consistency with integration dynamics, there has been little formal testing of non-integration mechanisms that when averaged across trials could mimic buildup patterns characteristic of integration (Latimer et al 2015). Thus, we devised non-integration models based on extrema detection that could viably generate the observed buildup dynamics of decision formation as well as choice accuracy trends, to examine the extent to which joint CPP-behaviour fitting can uniquely distinguish integration from non-integration mechanisms under such conditions.

We found that behavioural accuracy steadily improved with evidence duration, including from 0.8 to 1.6 sec, and that the CPP remained elevated up to the end of the stimulus period, together indicating protracted sampling. Behavioural accuracy on its own could be quantitatively well-captured not only by integration (drift-diffusion) but also by several non-integration models, and neither model class identified the setting of a decision bound. CPP dynamics in occasionally-presented, trivially easy trials indicated the setting of a bound, and models jointly fit to CPP waveforms and choice accuracy confirmed this, and ruled out some non-integration models. However, contrary to expectations, an extrema detection model that evoked a stereotyped signal ‘flagging’ decision termination at the first bound-crossing produced a fit that rivalled that of integration and reproduced the slow ramping build-up of the trial-averaged CPP and its modulation by evidence duration. All model classes benefitted from the inclusion of a collapsing decision bound and/or starting point variability to achieve a good fit, evidence for which was also observed in empirical motor preparation signals. While the degree of these features estimated by the Integration model was closer to that observed in the empirical signals, the Extremum-flagging model predicted drift rate modulations by difficulty that were closer to estimates from the empirical motor preparation signals. We go on to discuss the relative plausibility of these predictions and potential implications of these findings for other task conditions such as those where immediate responses provide information on decision termination timing, and what might be required of such an extremum-flagging model in order to further explain such data.

## Results

We recorded EEG from 16 human participants during a task where they viewed two oppositely oriented, overlaid gratings flickering in alternation with no noise or stochasticity. After a 1.2-sec baseline period of equal, 50% contrast, the gratings underwent an equal and opposite contrast change, and after an additional 1.6-sec stimulus-assessment period, participants reported by mouse button press which grating had higher contrast. In 80% of trials, the contrast differential was small (±10%) and lasted for 0.2, 0.4, 0.8, or 1.6 sec before stepping back to equal contrast and ultimately disappearing at 1.6 sec. For the remaining 20% of trials, a trivially easy (±40%) contrast difference was presented for 1.6 sec. The participants were not informed of the evidence duration manipulation, and this long, easy condition both reinforced the belief that all trials contained 1.6 sec of evidence (Figure 1A) and provided a means to probe the setting of a decision bound causing early terminations.

**Figure 1.**
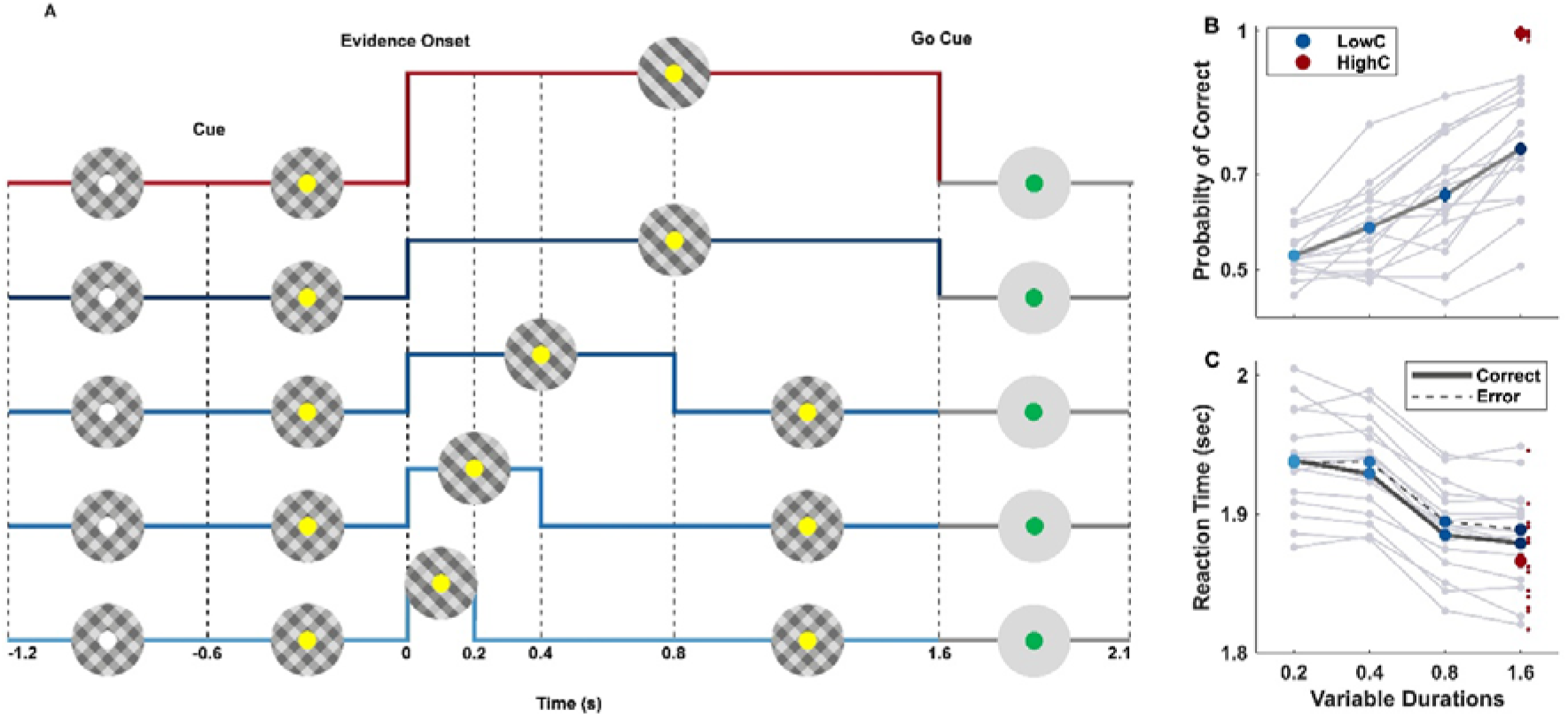
Contrast comparison task and behavioural results. (A) Stimulus and evidence time courses: The stimulus consisted of two interleaved gratings flickering on and off in alternation, whose relative contrast had to be judged. 0.6 sec after achieving fixation and triggering stimulus onset, the fixation point turned yellow to cue participants to prepare for the onset of contrast-difference evidence another 0.6 sec later. In all trials, the stimulus remained on for a full 1.6 sec following evidence onset, after which it disappeared and the fixation turned green to prompt the subject to make a left/right-hand button press to indicate that the left- or right-tilted grating had higher-contrast, within a 0.5-sec deadline. In 80% of trials, a weak (±10%) contrast increment/decrement was oppositely applied to the two gratings for 0.2, 0.4, 0.8, or 1.6 sec, returning to equal baseline contrast thereafter until stimulus offset. In the remaining 20% of trials, an easy contrast increment/decrement of ±40% was applied for the entire 1.6-sec period. (B) Accuracy significantly increased with the duration of the stimulus for low-contrast trials (blue dots) and was at the ceiling for the high-contrast condition (red dot). Individual participants’ accuracies are shown in grey lines. (C) The mean reaction time (RT; relative to evidence onset) slightly decreased in trials with longer durations and higher contrast levels. Correct responses (solid, with individuals also shown in grey) were also slightly faster than error responses (dashed) in this delayed response task. For b and c, data are mean ± s.e.m. after between-participant variance was factored out (subtracting each participant’s mean across conditions and adding the grand mean), in keeping with the repeated-measures experimental design.

Accuracy varied significantly across the five conditions (rmANOVA, F (4,60) = 150.47, p **=** 4.4×10^-16^, p^2^= 0.79, Figure 1B) and increased with duration within the low-contrast conditions (Linear Regression: β = 0.153, t(62) = 6.85, p < .001, R² = .431; see Table S1 for pairwise comparisons). Despite responses being deferred to stimulus offset, response time (RT) also significantly varied across conditions (rmANOVA F(4,60)=12.35, p_corr_=0.002, p^2^= 0.3), showing a mild downward trend with increasing duration in the low-contrast conditions (all three consecutive duration pairs p<2.48×10^2^), suggesting that subjects had not always reached commitment before the end of the stimulus period (Figure 1C; Table S2).

### Models fit to behaviour only

We began by comparing an integration model to two classes of non-integration model in terms of their ability to capture the improvements in accuracy averaged over all participants with increasing evidence duration. The Integration model entailed a simple drift-diffusion process (Ratcliff & McKoon, 2008; Figure 2A, top row, see Methods). The first non-integration model was an Extrema detection model, which entails sequential sampling but with the decision determined by an individual evidence sample reaching an extreme value (Hyafil et al., 2023; Stine et al., 2020; Waskom & Kiani, 2018; Figure 2A middle row). The second non-integration model was a Snapshot model, which entails neither integration or sequential sampling, instead basing the decision on a single randomly-chosen sample within the 1.6-sec stimulus-evaluation period (Stine et al, 2020). For all three models, we implemented both bounded and unbounded versions to test for early termination. In the Integration model, choices were determined by the sign of the first decision variable value to exceed a bound, or if the bound was absent/not hit, the value of the decision variable at the end of the stimulus. In three variants of the bounded Extrema detection model, choices were similarly determined by the sign of the first bound-crossing sample, and if not hit, choices were determined by i) a random 50/50 guess, ii) the sign of the last sample, or iii) the sign of the most extreme recorded sample (‘Extremum-tracking’). The latter extremum-tracking mechanism applied on all trials in the unbounded Extrema detection model. Finally, in the Snapshot model, the bound, when present, determined whether or not the randomly chosen sample would determine the choice, requiring a random 50/50 guess when the bound was not exceeded by the sample.

**Figure 2.**
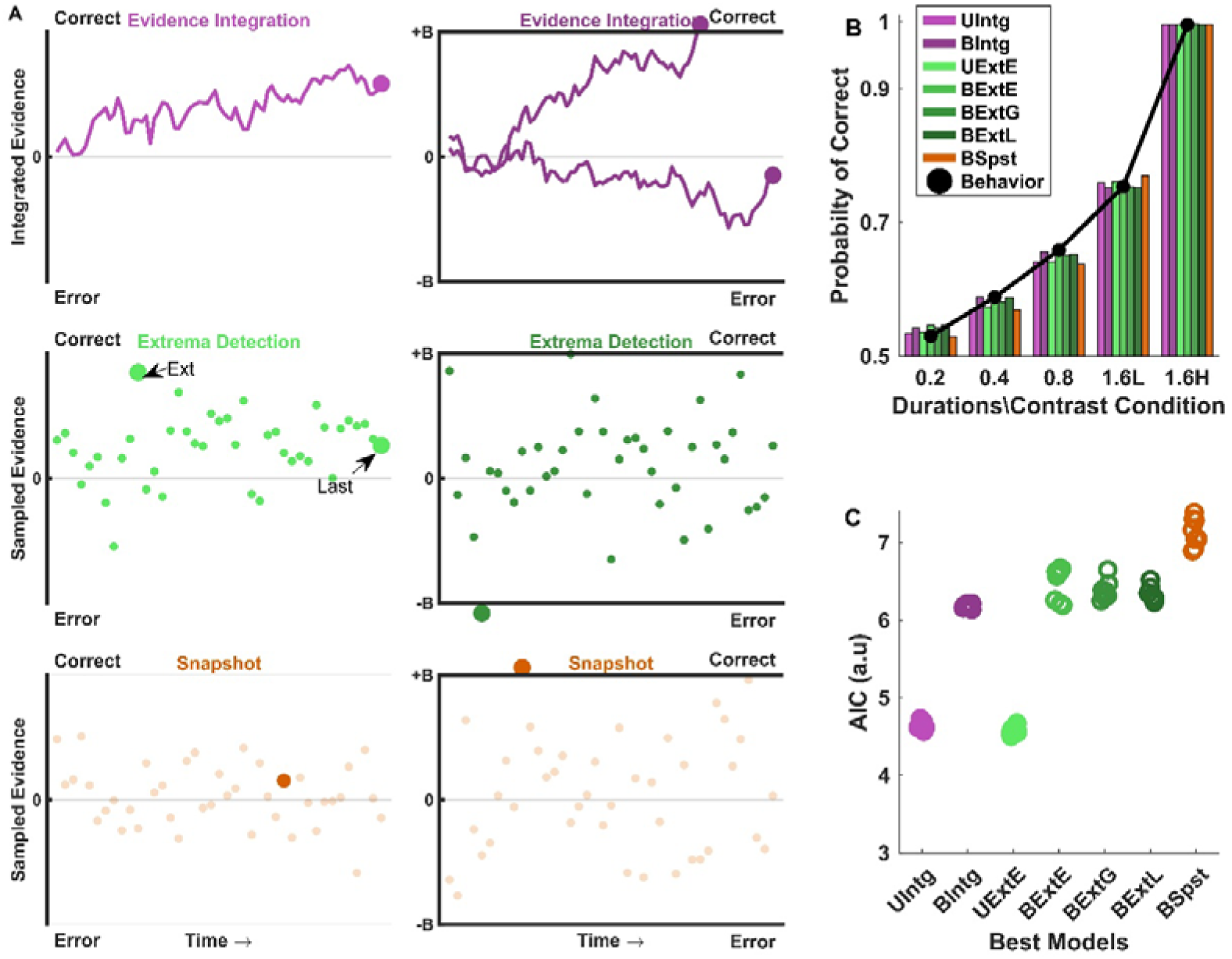
Alternative models and their fits to behaviour only. (A) Model schematics. In the unbounded integration model (‘UIntg;’ top left), the sign of the accumulated evidence at stimulus offset determines the choice, while in bounded integration (‘BIntg;’ top right), the decision is either terminated early by reaching the correct or error bound or otherwise determined in the same way as in the unbounded model. In the unbounded Extremum-tracking model (‘UExtE;’ middle left), the decision is based on the sign of the evidence sample that attains the maximum absolute value, which by making use of the full stimulus-assessment period can produce accuracy improvements with increasing evidence duration. In the bounded Extrema detection model (middle right), a choice is determined as soon as a single sample exceeds a bound or, if neither bound is reached, the choice is determined by: i) a random guess (‘BExtG’); ii) the sign of the last sample (‘BExtL’); or iii) the sign of the sample that attains the most extreme value (‘BExtE’). Finally, the Snapshot model, where the choice is based on a single sample chosen at random at any time (highlighted with the larger dot), is shown in the bottom row. In the unbounded version (‘USpst’), the decision is made based on the sign of the sample. In the bounded version (‘BSpst’), the decision is based on whether the sample surpasses the bound; otherwise, the decision is guessed. (B) Observed (black dots) and model-predicted (bars) accuracy, showing that the five best-fitting models can capture the observed accuracy improvement with duration well. (C) AIC values across the seven most competitive behaviour-only models show that models without bounds are favored due to parsimony, and selection among integration and most of the extrema detection mechanisms is largely inconclusive. Given that we computed G^2^ values through Monte-Carlo simulation, we conducted the fits with 10 different instantiations of noise, each shown here as a separate point (see Methods).

The model fit estimates and metrics are shown in Table 1. The drift rate for the high-contrast condition was in all cases higher than for low-contrast, as expected, and non-integration models generally needed much higher drift rates than integration models, simply because the same accuracies must be achieved based on one rather than many samples (Hyafil et al., 2023; Stine et al., 2020). All Extrema detection models with two drift rates were able to fit the data just as well as the equivalent Integration models (Figure 2B,C; see Figure 2 - Figure Supplement 1 for bound crossing histograms), with the snapshot model providing a fit that was only marginally worse. Further, the benefit of including a bound to fit the accuracy data was too small to warrant the additional complexity for any of the models (Figure 2C).

**Table 1:** Goodness of fit metrics and parameter estimates of models fit to behaviour only. Models beginning with U were unbounded, and beginning with B were bounded, with the bound parameter denoted by B. Drift rates for low and high contrast difference were denoted by D1 and D2, respectively; single-drift rate variants (denoted ‘_1d’) of the unbounded models provided a worst-case benchmark for each mechanism and s refers to common scaling parameter. G^2^ quantifies how closely the model quantitatively fits the data, while Akaike’s Information Criteria (AIC) adds a penalty for complexity. For both metrics, lower values indicate better fits. Colour shading highlights the competitive models depicted in Figure 2C.

| Models | | AIC | $G^2$ | D1 | D2 | B | s |
| --- | --- | --- | --- | --- | --- | --- | --- |
| Snap Shot | USpst_1d | 96.02 | 96.02 | 3.97 | - | - | 0.1 |
|  | USpst | 36.31 | 32.31 | 2.0 | 8.1 | - | 0.1 |
|  | BSpst | 39.4 | 33.4 | 2.0 | 9.06 | 0.00007 | 0.1 |
| Integration | UIntg_1d | 64.4 | 64.4 |  | - | - | 0.1 |
|  | UIntg | 4.53 | 0.53 | 0.057 | 0.20 | - | 0.1 |
|  | BIntg | 6.14 | 0.14 | 0.056 | 0.226 | 0.128 | 0.1 |
| Extrema Detection | BExtG | 6.17 | 0.17 | 10.3 | 11.65 | 0.034 | 0.1 |
|  | BExtL | 21.25 | 15.25 | 1.41 | 6.96 | 0.017 | 0.1 |
|  | UExtE_1d | 65.01 | 65.01 | 0.569 | - | - | 0.1 |
|  | UExtE | 4.54 | 0.54 | 0.382 | 1.59 | - | 0.1 |
|  | BExtE | 6.49 | 0.49 | 0.385 | 1.59 | 0.045 | 0.1 |

### Decision-related Neural Signals

Since behaviour alone was insufficient to arbitrate between integration and multiple extrema detection mechanisms and was inconclusive regarding the setting of a bound, we turned to neural signatures of decision formation to further inform and constrain the models. We began by appraising qualitative features of the CPP (Figure 3B) alongside signals reflecting sensory evidence encoding (Figure 3A) and motor preparation (Figure 3C), which inform model structure. The differential steady-state visual evoked potential (SSVEP), driven by the contrast difference between the correct (higher-contrast) and incorrect gratings, stepped from a zero baseline up to a level that scales with the contrast difference, remaining stable for the duration of the evidence before falling back down to zero after evidence offset (Figure 3A). This stable contrast-difference encoding verifies that the sensory evidence did not adapt over time, and although we cannot see whether time-variant effects yet occur downstream, we took this to justify the simplifying model assumption of stationary drift rate.

**Figure 3.**
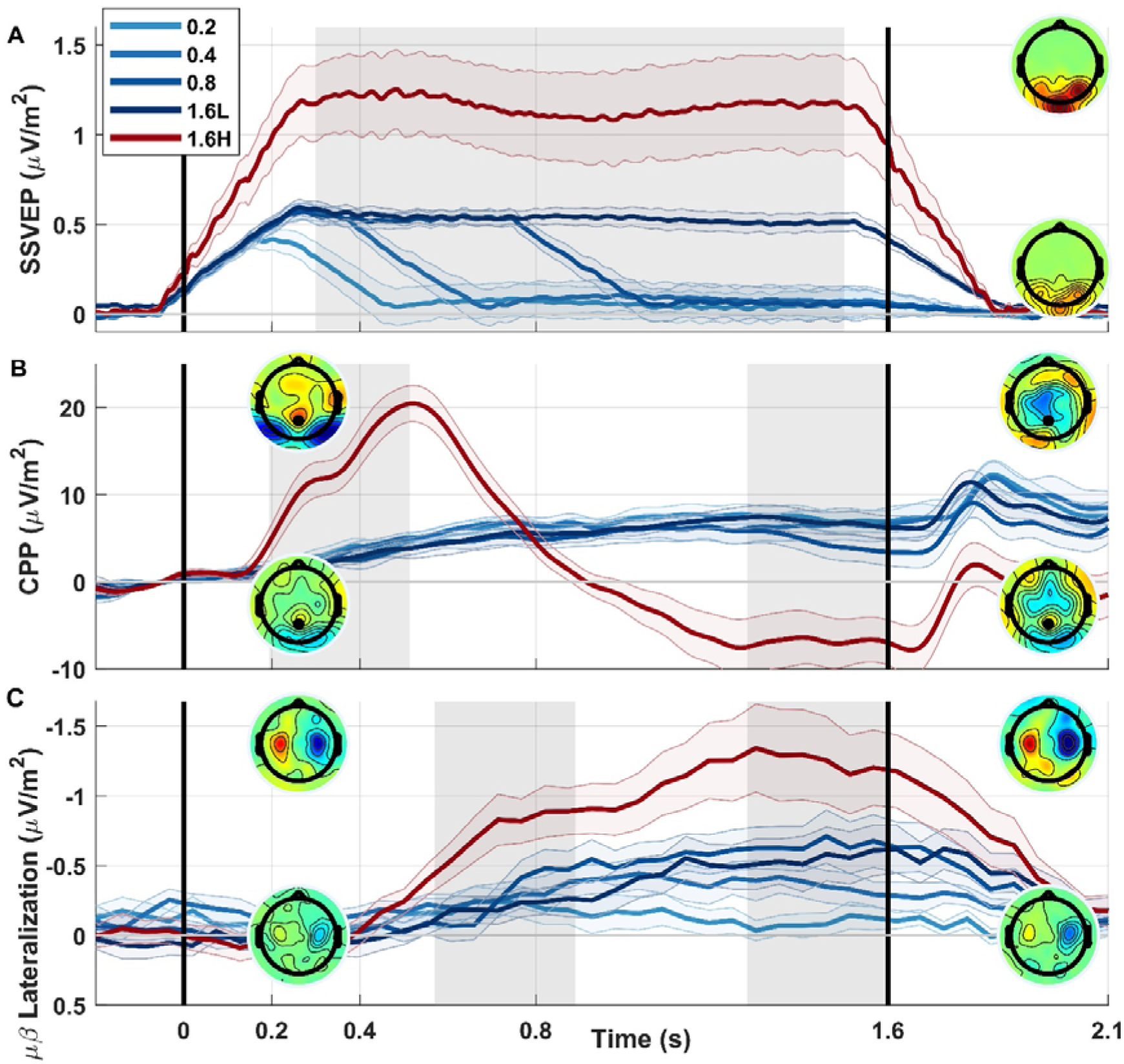
Neural Signals. (A) SSVEP showed a robust representation of sensory evidence encoding. Note that the signal ramps rather than suddenly steps simply due to the width of the window for spectral estimation (230 ms), and lags the physical contrast changes simply due to transmission delays to the visual cortex. Topographies show the average SSVEP amplitude from 300 ms to 1500 ms (shaded period) for the high contrast conditions (top) and the longest low contrast condition (bottom), with a primary focus at standard site Oz. (B) Event-related potential (ERP) at centroparietal sites averaged over trials (regardless of outcome) within each condition, showing that the centro-parietal positivity (CPP) undergoes a gradual buildup and sustained elevation for lower contrast conditions, and a peak at around 530 ms for the high contrast trials, at more than twice the amplitude. Topographies show ERP amplitude for the high contrast (top) and the average over low contrast conditions (bottom), for the shaded time ranges 210-530 ms (left) and 1280-1600 ms (right). (C) Motor preparation was measured by subtracting ipsilateral from contralateral mu/beta-band activity (MB; 8-30 Hz) with respect to the correct side. The topographies show the difference between trials favouring a left minus right response, again shown for both high contrast (top) and averaged low contrast (bottom) conditions in the shaded windows 570-890 ms (left) and 1280-1600 ms (right).

Turning to relative motor preparation reflected in Mu/Beta lateralisation (Figure 3C), we observed evidence-dependent buildup toward the correct alternative, remaining elevated until the delayed response as previously observed (Twomey et al 2016; McCone et al 2026), at a level that scaled with low-contrast evidence duration (rmANOVA, F (3,45) = 6.4, p=0.003, p^2^= 0.17). At the level of decision formation, the CPP similarly exhibited an evidence-dependent buildup initially. However, as in previous studies (Twomey et al 2016; McCone et al 2026), the high-contrast condition did not sustain but rather peaked and fell to baseline (Figure 3B), indicating a bound setting that leads to early terminations in those trials, and enabling us to rule out unbounded models (Figure 2 - Figure Supplement 2). This dynamic profile also rules out the Snapshot model, which assumes a uniform distribution of sample selection and hence predicts a uniform profile across the stimulus duration in any signal that might be evoked by that selection. In the low-contrast conditions, the CPP exhibited a sustained elevation above baseline, indicating protracted sampling in line with the observed accuracy improvements. In contrast to Mu/Beta lateralisation, the final CPP amplitude, measured in the 320 ms window ending at evidence offset, did not vary significantly across the four low-contrast evidence durations (rmANOVA, F (3,45) =1.23, p=0.3, p^2^=0.03). This lack of duration-dependence can be explained by the fact that whereas Mu/Beta lateralisation is a signed quantity whose polarity reflects the favoured alternative, the CPP always builds positively regardless of the favoured alternative, consistent with the sum of two underlying neural populations coding for the two alternatives (Afacan-Seref et al 2018). This means that noise variations do not cancel and the average CPP will exhibit a buildup even in conditions with weak or no evidence, blunting the distinction between noise-driven build-up and truly evidence-driven build-up for very weak evidence strengths (Steinemann et al., 2018). While all of these signal properties are consistent with previous observations, the data do not provide qualitative grounds to judge whether they arise from integration or non-integration mechanisms, which we next investigate through quantitative modelling.

### Neurally-constrained models

We proceeded to devise a decision-signal generating mechanism for the competing bounded models that could not be resolved based on behavioural modelling alone, in order to constrain them further by comparing simulated trial-averaged decision variable (DV) waveforms with observed CPP waveforms. We chose the CPP to provide the neural constraints because one of our core interests was the temporal extent of sampling, and the CPP’s peak and fall dynamics render it more diagnostic of early decision termination than Mu/beta, which sustains until response.

In the Integration model, the DV was computed as the perfect (lossless) integration of the momentary evidence samples over time with a fall-down back to zero following a bound crossing (Figure 4B). For extrema detection, we devised two mechanisms by which buildup dynamics could arise in average decision signals. In the Extremum-tracking model (corresponding to ‘BExtE’ above), a running record of the most extreme evidence sample witnessed so far was encoded in the decision variable, again with a fall-down to zero following a bound crossing. In principle, this enables an early peak-and-fall in the high-contrast condition and a slower more sustained buildup in the low-contrast conditions (Figure 4C). As a second Extrema detection mechanism, we simulated a DV involving no evidence-driven buildup dynamics on any single trial but rather a stereotyped signal marking each single-trial bound crossing, which when averaged across trials could approximate a gradual buildup. In this “Extremum-flagging” model, we assumed that bound-crossings triggered a Half-Sine-shaped (upper half; Weindel et al 2025) temporal function with an amplitude and width that was fixed across trials (Figure 4D), which would be expected to cluster at early latencies in high-contrast trials and spread over longer latencies in low-contrast trials, thus plausibly approximating the observed CPP dynamics. Though we use a neutral ‘flagging’ function here, the idea is consistent with classic theories of the centroparietal P300 or P3b component, which posit that it reflects a process triggered by completion of a perceptual judgment, such as “context updating” (Donchin & Coles 1988) or response facilitation (Nieuwenhuis et al., 2005; Verleger, 2020, see Discussion). If no bound was reached by stimulus offset, there was no flag signal elicited, and the subject made a random guess (see also Figure 4 - Figure Supplement 8 for demonstration that neurally-constrained modelling conclusions do not change with a last-sample instead of guess default). Since the half-sine flagging signal inherently incorporates a quarter-sine-shaped fall-down, we also used a quarter-sine-shaped fall-down in the Integration and Extremum-tracking models for consistency.

**Figure 4.**
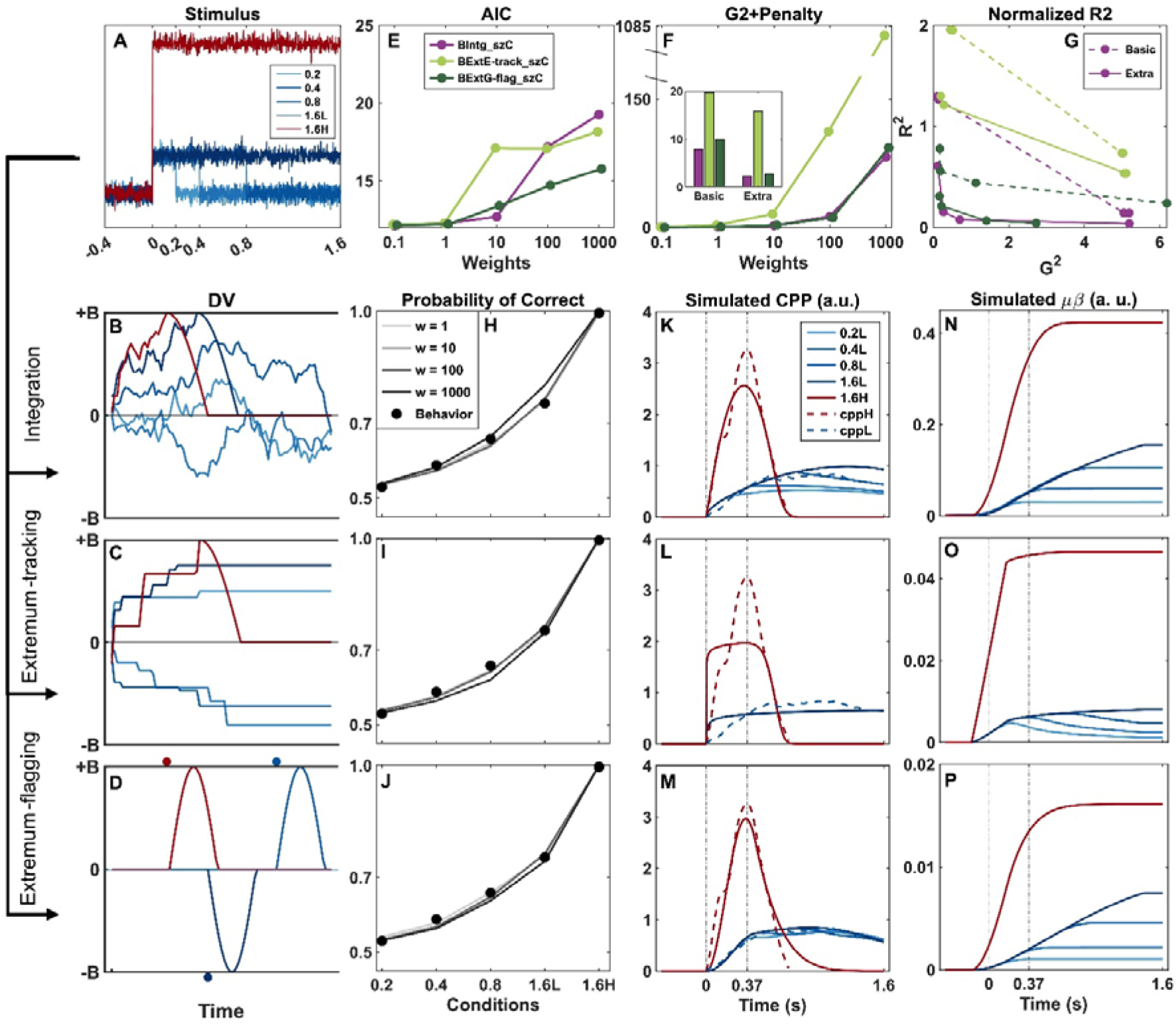
Neurally constrained models and their predictions. Among the models with various additional parameters, starting-point variability and collapsing bounds stood out as helping to achieve good CPP waveform fits without substantially sacrificing behavioural fits (Figure 4 - Figure Supplement 1), and given that both of these features were exhibited empirically in motor preparation signal effects (Figure 5), we show these models here (but see Figure 4 - Figure Supplements 4-6 for basic versions of the models without the extra parameters). (**A**) Single-trial sensory evidence traces were simulated for each of the five conditions using boxcar functions (stepping up during positive evidence) with added Gaussian noise as in behaviour-only models (one example for each condition shown). (B-D) single-trial examples of the decision variable in each of the three competing models, again simulating one trial per condition. (B) In the **Integration model,** the evidence was integrated over time until it reached a bound, after which it fell back to zero. Where the decision variable did not hit the bound the decision was made based on the sign of accumulated evidence at 1.6 sec. (C) In the **Extremum-tracking** model, the DV keeps track of the most extreme value so far, until it reaches the bound, after which it falls to zero. The sign of the extremum determined the choice, and the most extreme sub-bound value was used in the event that it did not cross the bound. (**D**) In the **Extremum-flagging** model, an all-or-nothing Half-Sine-profile signal marking choice termination onsetting at the bound crossing time (see main text). As in the other models, one example DV trace per condition is shown; for three of these (red and two dark blue traces) the bound was reached, while in the others it was not, so no signal was generated (flat light blue lines), and the choice was made as a random guess. In all 3 models in this figure, starting point variability and collapsing bound parameters are included, and they have an equal number of free parameters. (E) Akaike’s Information Criterion (AIC) derived from only the behavioural component of the neurally-constrained fit, plotted as a function of the weighting of CPP-waveform constraint. With increasing emphasis on neural data in the fit, the fit to behaviour was compromised to varying extents, and the Extremum-flagging model was best at retaining good behavioural fits at higher neural weights. (F) Overall neurally-constrained objective function (behavioural G^2^ value plus weighted neural penalty quantifying divergence in observed vs simulated CPP) as a function of neural constraint weighting. The inset compares the objective function values with starting point variability and collapsing bound parameters (labelled ‘Extra’ in G) to the ‘basic’ models without either feature, at w = 10. (G) Neural fit (R^2^) plotted against behavioural fit (G^2^), illustrating a trade-off between how well the models can capture behaviour and the CPP up to w = 100. The closer the joint values come to the origin the better they are at simultaneously fitting both. (H-J) Predicted behavioural accuracy for a range of neural constraint weightings (‘w’). (K-M) Simulated average CPP (where the DV is rectified on each single trial in keeping with CPP’s always-positive nature; see Methods) for the best fits of each model at a neural weighting of w=10 (see Figure 4 - Figure Supplement 2 for simulations from all weightings, and Figure 4 - Figure Supplement 3 for predicted bound crossing frequencies). The simulated, normalised average CPP (solid lines) is shown for all five conditions. The normalised empirical data is also shown (dashed lines; see Methods for details) with the waveforms for the four low-contrast conditions averaged together, as they were for the neurally-constrained fitting. (N-P) Simulated differential motor preparation. This is based on the same simulated decision variable as in (K-M) but now retaining its differential form by not taking the absolute value and having the signal sustain at the bound level after it is crossed. This is not normalised since it is not directly fit to empirical data. Smoothing is applied equivalent to the Fourier windowing in the Mu/Beta measurements.

To constrain the models to fit CPP waveform dynamics in addition to accuracy, we added a penalty term quantifying the divergence between normalized real and simulated CPP waveforms (200-900 ms for high-contrast, 200-1600 ms for averaged low contrast; see Methods) to the goodness-of-fit metric to be minimised. We comprehensively examined the relative ability of the three model classes to jointly fit accuracy and CPP dynamics by fitting the models with not only the basic bound, drift rate and sine-width parameters, but also plausible additional parameters, tested one at a time. These included drift rate variability, starting point variability, sequential sampling duration (allowing premature quitting), sampling onset time (allowing sampling to begin before or after evidence is available), collapsing bound, and in the case of integration, leakage (see methods for full explanation of each). We fit all models with five neural penalty term weightings (w = 0.1, 1, 10, 100, 1000) to examine how the behavioural and neural fits change as a function of emphasis placed on reproducing the CPP dynamics.

In Figure 4 we compare simulations of the three model classes including both starting point variability and collapsing bounds, which helped achieve a good joint fit in all cases (Figure 4 - Figure Supplement 1). AIC derived from the behavioural fit element of the objective function generally increased with neural weighting (Figure 4E), demonstrating an expected trade-off where the higher the emphasis on a close fit to CPP waveforms, the more the behavioural fit is compromised. Looking at the overall objective function (G^2^ plus the neural penalty term; Figure 4F), the Extremum-tracking model clearly could not achieve a competitive fit, but the Integration and Extremum-flagging models achieve similarly close fits. Although the Integration model appeared to gain an advantage at extremely high neural constraint weightings (Figure 4F), this came with a greater sacrifice to behavioural fit (Figure 4E). We traced the neural-behavioural trade-off by plotting dissimilarity between simulated and observed data for the neural fit against the behavioural fit (Figure 4G). When endowed with starting point variability and collapsing bounds, both the Integration and Extremum-flagging models were able to achieve a better neural fit without sacrificing behavioural fit, compared to ‘basic’ versions of the models without those features (Figure 4G). However, which of the two models gives the overall better fit depends on the relative importance one places on fitting behaviour versus neural dynamics. This is reflected in the fact that the trade-off curve for neither model lies fully inside (closer to the origin) that of the other model, which would have indicated a winner independent of neural constraint weighting. Accuracy fits for a range of weightings (‘w’) produced a close fit to behaviour (Figure 4H-J). We accordingly evaluate the models in terms of their ability to capture key features of the neural data. The average DV dynamics simulated from each model illustrate that these two models could both qualitatively produce average CPPs with a shallow incline and continued elevation in the low-contrast conditions, while also producing a steep initial rise and early peak in the high-contrast condition (Figure 4K,M; Figure 4 - Figure Supplement 2). Although the Integration model appeared to produce some duration dependence in low contrast amplitudes (Figure 4K), this feature was not consistently observed across neural weightings (Figure 4 - Figure Supplement 2). By contrast, the Extremum-tracking model could produce a peak-and-fall morphology in high-contrast, and sustained elevation in low-contrast conditions, but could not do so while retaining a gradual evidence-dependent buildup (Figure 4L, Figure 4 - Figure Supplement 2). The DV waveform simulations in Figure 4, as well as the estimated parameters and fit metrics in Table 2, are for a neural constraint weight of w = 10, for which both competitive models achieved a good fit to CPP dynamics without substantively sacrificing behavioural fit.

**Table 2.** Fit metrics and parameter estimates for neurally-constrained models with a neural-constraint weighting of w = 10. B refers to the bound; D1 and D2 refer to the drift rates in the low- and high-contrast conditions, respectively. FlagWidth refers to the width of the half-sine, sz refers to starting-point variability, K refers to the temporal slope of the collapsing bound and s refers to within-trial noise, the common scaling parameter (see Tables S3 for fits using a nonlinear collapse function, or Extremum-flagging with last-sample default, with similar findings). Models including starting point variability and collapsing bounds are distinguished from basic models without those features, by the model name ending ‘_szC.’

| w = 10 | G<sup>2</sup>+Penalty | B | D1 | D2 | FlagWidth | sz | K | s |
| --- | --- | --- | --- | --- | --- | --- | --- | --- |
| BIntg | 7.9864 | 0.3591 | 0.0566 | 1.0991 | 0.6688 | - | - | 0.1 |
| BIntg_szC | 2.2533 | 0.4683 | 0.1607 | 1.4049 | 0.2502 | 0.9419 | 0.312 | 0.1 |
| BExtE | 19.7663 | 0.035 | 0.3815 | 10.865 | 0.9 | - | - | 0.1 |
| BExtE_szC | 15.83 | 0.059 | 0.385 | 16.79 | 0.1 | 0.0018 | 0.46 | 0.1 |
| BExtG | 9.9581 | 0.0145 | 0.5079 | 2.0663 | 0.52 | - | - | 0.1 |
| BExtG_szC | 2.7589 | 0.015 | 0.4387 | 2.1614 | 0.4462 | 0.000135 | 0.1062 | 0.1 |

Since model fits based on the CPP were equivocal, we evaluated the models further by simulating relative motor preparation from the best fitting parameters found in the CPP-constrained fits of all models (assumed to sustain at bound level once crossed). These exhibited dynamics that were qualitatively similar to the empirical MB lateralisation signal, including a steeper rise and higher sustained level for the higher contrast difference and greater amplitude variations across low-contrast durations than seen in the CPP. Temporal profiles varied across models, however: While both the Integration and Extremum-flagging models exhibited sustaining across the low contrast conditions (Figure 4N,P) similar to the empirical MB signal, the Extremum-tracking model showed a drop beginning at evidence offset in each condition, since many trials would find a negative extremum arise by chance during the zero-evidence lead-out period (Figure 4O). Between the Integration and Extremum-flagging models, a difference is apparent in the degree of average buildup in lateralised motor preparation for the low-contrast conditions relative to the maximal level reached by the high-contrast condition, with the Extremum-flagging model reaching higher levels closer to those observed in the empirical MB signals (Figure 3C). Indeed, with increasing neural constraint weighting, the Integration model tends to estimate a proportional difference in drift rates for high-vs low-contrast conditions that increasingly exceeds that of the physical contrast differences (40% vs 10%, i.e. a factor of 4); at w=10, this proportional difference is a factor of 8.7 for the Integration model, compared to 4.9 for the Extremum-flagging model, and a factor of 3.5 measured from the empirical, relative motor preparation slopes in the window 500-700 ms (Figure 3C). Indeed, when the drift rates were constrained to scale in direct proportion with physical contrast to force a 4-fold difference, the Integration model was far more hampered than the Extremum-Flagging model (Figure 4 - Supplement 7; see Discussion).

To further explore empirical Mu/Beta dynamics against model predictions, we plotted the contralateral and ipsilateral Mu/Beta signals separately, broken out by decision outcome (correct/error; Figure 5). In immediate-response tasks, this analysis approach has previously revealed that the two competing motor alternatives race each other up to a stereotyped level at the time of response that is invariant across stimulus conditions or reaction time (RT), indicating a fixed threshold associated with triggering the contralateral action (Corbett et al., 2023; Devine et al., 2019; Feuerriegel et al., 2021; Kelly et al., 2021; O’Connell et al., 2012). When evidence onset is anticipated, motor preparation reflected in Mu/Beta amplitude reduction contralateral to both hands has been found to begin building towards threshold even before evidence appears (Kelly et al., 2021), consistent with a dynamic ‘Urgency’ signal, defined as an evidence-independent buildup component that adds to cumulative evidence, gradually reducing the amount of cumulative evidence required to reach threshold and thus effectively implementing a collapsing decision bound (O’Connell et al., 2018). Moreover, fluctuations in pre-evidence Mu/Beta activity have been shown to predict trial-to-trial variability in choice and RT, in the way accumulator starting point variability does in models (Steinemann et al., 2018).

**Figure 5.**
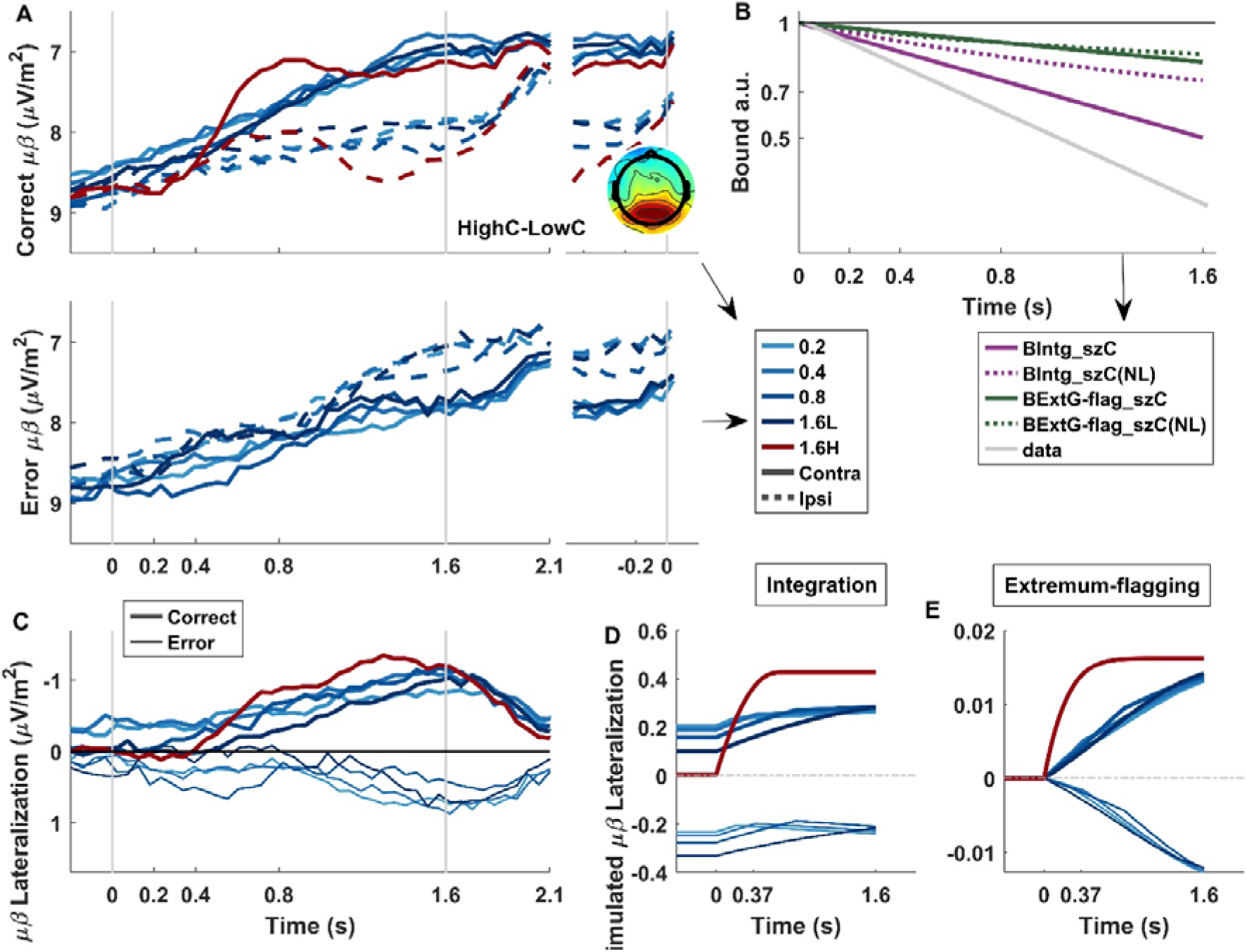
Observed and simulated motor preparation and bound dynamics. (A) Mu/beta (MB) decreases reflecting building motor preparation, contralateral and ipsilateral to the correct response, in correct (top) and error trials (bottom), shown aligned to evidence onset (left) and response (right). Contralateral traces, on correct trials, and ipsilateral traces, on error trials, reach a fixed threshold prior to response (marked by horizontal grey dashed line). Interestingly, on easy, high-contrast trials, this threshold (marked by the grey dashed horizontal line to aid comparison) is reached at approximately 800 ms and sustains at just below that level until the response. Note that on high-contrast trials, contralateral MB appears to reach a less decreased level prior to response compared to low-contrast trials. The topography inset shows the difference in MB between high-contrast and low-contrast conditions, revealing that this difference is unlikely motor-related, and more likely reflects greater attentional engagement in the harder low-contrast trials, known to be reflected in posterior alpha activity (Foxe & Snyder, 2011; Kelly & O’Connell, 2013). (B) Collapsing bound estimated from the empirical data and the model fits. Linear (solid) and non-linear (‘NL,’ dashed) collapsing bounds were calculated for Integration and Extremum-flagging models. All traces were normalised to the initial bound height. (C) Observed Mu/beta lateralisation for correct and error trials, demonstrating baseline choice-predictive effects that arise from starting-point variability. This can be compared to simulated average relative motor preparation at a neural constraint weighting of w = 10 for the Integration (D) and Extremum-flagging (E) models. Note again that models do not include non-decision delays, and so evidence-dependent dynamics begin at time zero in the simulated traces, whereas they are clearly delayed as usual in the empirically observed data. All simulations are from models containing both starting point variability and collapsing bounds.

Here, despite the fact that the response had to be withheld until the post-stimulus response cue, MB-indexed motor preparation exhibited both of these effects (Figure 5). It rose significantly from -0.2 sec to 0 sec relative to evidence onset (main effect of time, F(1,14) = 6.43, p = 0.02, p^2^= 0.001), reflecting anticipatory urgency, before exhibiting evidence-dependent buildup after evidence onset, with contralateral preparation steepening in high-contrast trials, peaking at around 700 ms, and maintaining almost this level until the response period (Figure 5A). By averaging motor preparation traces across all trials regardless of condition or outcome, and extrapolating from a linear fit in the same baseline period, we could estimate the temporal profile of the collapsing bound, normalised relative to the initial bound height (Figure 5B, grey line). This could then be compared against the profile of collapsing bounds estimated from the models. Although both models showed a collapse that was less extensive than the empirical estimate, the Integration model provided a closer match than the Extremum-flagging model. Further examining Mu/beta lateralisation broken out by correct/incorrect outcome (Figure 5C), it is clear that in low-contrast trials, baseline levels of motor preparation already favoured the response that was eventually chosen before any evidence was presented (correctness x side interaction F(1,14) = 8.85, p = 0.01), reflecting significant starting-point variability that impacts eventual choice. As expected, the more difficult the trial (shortest duration), the greater the degree to which choice is linked to the baseline level randomly favouring one or the other alternative, and this is recapitulated in the Integration model (Figure 5D) but not the Extremum-flagging model (Figure 5E), the latter of which did not estimate appreciable starting-point variability at any neural constraint weighting (Tables S3, S5-8).

## Discussion

Sequential sampling models have long been known to provide excellent quantitative fits to decision behaviour on diverse tasks (Ratcliff et al., 2016), but the same ubiquity cannot be assumed of the role of protracted integration in practice, across all perceptual tasks (Cisek et al., 2009; Ludwig et al., 2005; Stine et al., 2020; Uchida et al., 2006). The present study examined a contrast-comparison task whose long, noiseless stimuli and delayed decision reports might seem to obviate protracted sampling, but whose subtle feature differences may be vulnerable to internal neural noise and hence benefit from protracted integration. This emulates real-life situations where subtle features need to be judged in good visibility conditions in order to guide later actions, but is a situation that, due to the absence of information on decision timing and the dominance of one fixed evidence strength, offers very limited behavioural data for adjudicating between alternative mechanisms. Moreover, given the increasingly apparent task-dependence of neural decision mechanisms (Okazawa & Kiani, 2023; Parés-Pujolràs et al., 2025; Shushruth et al., 2022), one cannot simply extrapolate them from distinct task types such as immediate-response or expanded judgement tasks. Hence, to gather more information for model comparison while retaining the core task demands of interest, we manipulated the duration of the weak evidence without telling the participants, and traced the average neural dynamics of decision formation in the centro-parietal positivity (CPP) and spectral EEG measures of motor preparation. We found convincing evidence for protracted sampling: accuracy steadily improved with evidence duration, including a substantial increase from 800 to 1600-ms duration, and in difficult trials the CPP remained elevated throughout the stimulus period. Backing up this apparent utilisation of the full stimulus duration, models allowing for flexible sampling onsets generally estimated this close to evidence onset, and models allowing for leak estimated this as very low (0.0002 at w=10).

CPP dynamics also established the operation of a bound, which was reached early and often for the rare strong-evidence trials, but less frequently and later for the weak trials. At the motor preparation level, pre-evidence, baseline biases toward the ultimately chosen alternative indicated the operation of starting point variability, and dynamic anticipatory preparation suggested a collapsing bound effect. However, quantitative modelling revealed that although the addition of neural constraints did offer many more insights than was possible from behaviour alone, it could not identify one uniquely viable account of the data; specifically, when allowing for starting point variability and collapsing bounds, similarly close fits to both behaviour and CPP dynamics could be attained by an integration model and a non-integration model based on extrema detection.

Extrema detection models were proposed as accounts of decision behaviour decades ago (Watson, 1979), and although integration-based models such as the DDM have dominated behavioural modelling in the intervening years due to their mechanistic appeal and versatility, it was recently shown that these two accounts are surprisingly competitive in their ability to explain a range of important behavioural effects (Stine et al., 2020). Here, we went a step further by considering plausible ways in which such an extrema detection model could produce a gradual buildup in its trial-averaged decision variable dynamics, and thereby predict neural decision signal dynamics. One mechanism, which continually tracked the most extreme sample encoded so far (‘Extremum-tracking’), could fit behavioural data but could not simultaneously capture CPP dynamics because it predicts an initially steep and decelerating temporal profile that is not observed in the CPP (Figure 4L). A second mechanism, in which a stereotyped signal flagged single-trial bound crossings that could produce gradual buildup when averaged, could capture both behaviour and CPP dynamics to a degree comparable with Integration. Although this may be the first time that centroparietal signal buildup has been captured quantitatively through a non-integration mechanism, the general idea of centroparietal activity being triggered by decision completion was a part of many decades-old hypotheses regarding the function of the classic ‘P3b’ ERP component, centred on concepts of stimulus-response activation, context updating and memory storage (Polich 2007; Verleger et al 2020). In one physiologically-elaborated account, Nieuwenhuis et al (2005) proposed that the P300/P3b reflects a phasic response of the neuromodulatory locus coeruleus-norepinephrine (LC-NE) system arising as the outcome of internal decision-making processes. Triggered by each discrete decision bound crossing, such a physiological response could conceivably unfold over a duration similar to what the model estimates here. However, while successful in this particular delayed-response paradigm, the degree to which such a process could account for decision signal dynamics previously observed on other tasks such as those with immediate-response requirements, remains unclear. In discrete-trial, forced-choice tasks, the response-locked CPP builds with a shallower slope and reaches a lower amplitude for weaker relative to stronger evidence, which in bounded accumulation frameworks is explained by evidence-dependent drift rates and a collapsing bound (Steinemann et al., 2018). The version of the Extremum-flagging model in the current study assumes a stereotyped signal triggered by each bound crossing and thus it should not show any such variation by evidence strength when aligned to the response; however, in principle, if one allowed for the possibility that the flag signal was smaller in amplitude for weaker stimuli (e.g. if it scaled with confidence) and more temporally jittered with respect to response execution, it could be possible to produce response-locked CPP waveforms that cross-over each other for high vs low evidence strength (because low-evidence builds more slowly to a lower final amplitude) in the way that is observed. Whether this can be achieved with reasonable estimates of non-decision delays will have to be explicitly tested in future work.

While the Integration and Extremum-flagging models agreed on the operation of protracted sampling throughout the stimulus period unless a bound was reached, and the relative insensitivity of the simulated CPP to duration due to its always-positive nature, there were notable differences in some key quantitative predictions. The Integration model predicted a substantially higher ratio of drift rate for high versus low contrast (8.7 at w = 10) than the Extremum-flagging model (4.9), which was closer to the ratio of the contrast difference itself (a factor of 4). In principle, a drift rate ratio exceeding the contrast difference ratio could plausibly arise from the well-known nonlinear contrast response function of early visual cortical neural activity (saturating typically before 50%, especially in extrastriate areas) and the fact that the high-contrast condition had a highly salient evidence onset to trigger attentive sampling, compared to the low-contrast condition. However, the ratio of buildup rates of relative motor preparation signals (3.5) suggests a drift rate ratio much closer to the physical contrast ratio, even allowing for some degree of temporal blurring associated with the time-frequency analysis. The excessive drift rate ratio of the Integration model was not observed in the fit to behaviour alone, and generally grew with increasing model-weighting of the CPP waveforms, which indeed show a more extreme relative buildup in high-contrast versus low contrast conditions compared to Mu/Beta lateralisation. A possible way to reconcile this with the Integration model lies in the strong bilateral occipital negativity, labelled ‘N2,’ which, consistent with previous work (Loughnane et al 2016), is large and transient for the salient step-transition in the high-evidence condition. Follow-up examination showed that the peak of the N2 coincides with an initial intermediate peak during the buildup of the CPP (see Figure 3 - Figure Supplement 1), suggesting that the opposite end of the dipolar neural generators of the occipital N2 may contribute to the positive amplitude measured at the CPP scalp site, thus artificially inflating the steepness of initial buildup in the high-contrast condition which the Integration model had to capture through an extreme drift rate ratio. While our aim in the present study was to leverage the peak-and-fall property of the high-evidence CPP to decipher whether and where a bound is set, this analysis demonstrates a limitation whereby overlapping activity can compromise the directness with which the CPP reflects decision variable dynamics, and this warrants further research into related ways in which the linking function may need adjustment. Meanwhile, the quantitative degree of starting point variability and bound collapse was substantially greater in the Integration model, toward the levels exhibited empirically in motor preparation signals. Because the Extremum-flagging model relies on a precise bound setting on an evidence representation with a high drift rate in order to explain the accuracy improvements across durations, it is highly sensitive to even a small bound collapse, and starting point variability effectively adds noise that cannot be absorbed by the precise bound setting, hence leading the Extremum-flagging model to favour a near-zero level of starting point variability. The inferred operation of collapsing bounds is intriguing from the point of view that responses were deferred in this task until after stimulus offset; however, given the prevalently weak evidence strength and tight 500-ms deadline after stimulus onset, it may be that subjects experienced urgency to commit to a choice in a timely fashion. Indeed, it has recently been shown that the dynamics of the Contingent Negative Variation in delayed response tasks is consistent with a cognitive-level urgency to terminate a decision even without the requirement of immediate response execution (McCone et al., 2026).

Our joint modelling approach aims to mitigate the model comparison limitations inherent in behaviour-only models fit exclusively to accuracy data from delayed-response tasks, by leveraging neural measurements for additional constraints. Nevertheless, there are several limitations worth noting. First, while we fitted models to the grand-average neural and behavioural data across participants, due to the noisiness of the CPP signal at the individual level, this precludes assessment of potential individual differences. To the extent that individual data patterns such as the accuracy trends over duration diverge systematically from the grand average, this limitation also means that we would miss models that may be decisively distinguishable for certain individuals, despite being equivocal at the grand average level. Second, in the neurally-constrained modelling approach we adopted here, we conducted a sensitivity analysis where changes in the model fits were evaluated as a function of increasing emphasis on fitting the neural data. A potential shortcoming of this is that it is hard to announce a quantitative ‘winner’ among the models in cases such as this, as this depends on how much one favours behaviour versus neural data; while Bayesian modelling in theory provides a principled means to determine weighting based on reliability as defined by data variance, it does not address the less quantifiable degree to which neural decision indices can be assumed to faithfully map onto latent decision processes. We believe that this range-of-weightings approach was the right one in this case, because in principle, with such narrow margins it could be possible to make further tweaks to aspects of the models to produce slightly better quantitative fits and it would be hard to know when to stop in an unbiased way. One element, for example, that in principle could impact the relative quantitative fit of the models pertains to the fall-down dynamics of the CPP once a bound is reached. While we have access to this in immediate response paradigms, we do not in delayed-response paradigms and it is not clear that one can be readily extrapolated from the other. Aside from the uncertain shape of the CPP fall-down, which in the case of integration could be linear rather than the sine-shaped function used here for consistency with the Extremum-flagging model, fall-down dynamics would also be affected by how soon the accumulation quits or the flagging signal onsets following commitment (bound-crossing), and whether there is some amount of post-commitment sampling as has been observed in immediate-response tasks (Murphy et al 2015; Grogan et al 2023), all of which may have some impact on the morphology of the average waveforms. Also, we assume a linear mapping between decision variable units in our models and the empirical decision signals (CPP and Mu/Beta), when the mapping may not be precisely linear in practice. Finally, a larger number of subjects may have smoothened and refined the empirical average waveforms used to constrain the models, although the potential changes in quantitative fits are unlikely to have substantively altered the model comparison outcome, relative to the impact of tweaks related to such nuances as fall-down or post-commitment dynamics.

In summary, this study presents a non-integrative sampling mechanism that can rival Integration in capturing not only behavioural accuracy of delayed perceptual decision reports about a protracted stimulus, but also decision variable dynamics reflected in neural decision signatures during that stimulus, which lends itself to further quantitative testing in a broader range of conditions.

## Acknowledgments

H.H., K.S.M. and S.P.K. were supported by a Wellcome Trust Investigator Award (219572/Z/19/Z). R.G.O. was supported by Horizon 2020 European Research Council Consolidator Grant In Decision 865474. We acknowledge the Research IT HPC Service at University College Dublin for providing computational facilities and support that contributed to the research results reported in this paper.

## Methods

### Data Collection

#### Participants

Sixteen healthy human participants (9 males, 4 females (aged 18-40); the gender of three participants was not recorded) were invited to take part in our experiment, which was approved by the Human Research Ethics Committee for the Sciences, University College Dublin in accordance with the Declaration of Helsinki. Sample size was not based on formal power analysis, but was close to similar EEG studies. All participants had normal or corrected to normal vision with no history of psychiatric diagnosis, sensitivity to flickering light, or head injury. Participants gave us written and informed consent and were compensated at €10 / hour. All together the participants completed 188 blocks of 80 trials (15040 trials in total).

#### Contrast Discrimination Task

Participants were instructed to perform a delayed-response, two-alternative forced choice contrast discrimination task, programmed using PsychToolbox-3 (Brainard & Vision, 1997; Kleiner et al., 2007; Pelli, 1997) in MATLAB and presented on a gamma-corrected 51 cm Cathode Ray Tube monitor (Dell E771p) with a refresh rate of 75 Hz and resolution of 1024 768. Participants were seated at a viewing distance of 57 cm inside a dark, sound-attenuated booth with their heads stabilised in a chin rest with forehead support and their left and right thumbs resting on the left and right buttons of a symmetrically shaped computer mouse, which formed the response alternatives. Eye position was monitored continuously throughout the experiment using a remote eye tracker (EyeLink 1000, SR Research, 1,000Hz).

### Stimulus and procedure

The stimulus contained two interleaved grating patterns with a spatial frequency of 1 cycle per degree tilted at −45 and +45 degrees relative to the vertical meridian (Figure 1A), which flickered on and off in alternation, with each presented for one screen refresh on every 4th refresh, thus generating two phase-opposed steady-state visual evoked potentials at 18.75 Hz that cancel each other when contrasts are equal (see below). A central fixation dot (0.2° diameter, grey) was presented, on which the participants fixated, followed 1200 ms later by the overlaid grating stimuli at 50% contrast (baseline period). After 600 ms of the baseline period, the fixation dot changed colour to yellow to warn the participants that evidence would be presented after another 600 ms. Evidence consisted of an equal-sized increment/decrement in the contrast of the two gratings such that one appeared brighter, which lasted for up to 1600 ms with the stimulus returning to baseline for shorter durations until the stimulus was finally removed at the end of the 1600 ms period. Participants were not informed that the duration of evidence would vary, and the task structure encouraged a belief that it would always last 1600 ms. ّThere were five equiprobable, randomly interleaved evidence conditions. In one easy condition, a high contrast difference (±40%, i.e., one grating 10% and the other 90%) was shown for the full 1600 ms, which served to reinforce the belief that evidence lasted for 1600 ms. The remaining four conditions consisted of a low contrast difference (±10%), which was presented for 200, 400, 800, or 1600 ms. This contrast difference was difficult, with performance reaching 75% only on the longest duration trials (Figure 1B), so that participants were unlikely to detect with any confidence that the stimulus returned to baseline on the shorter duration trials (Figure 1A). The subjects were instructed that difficulty would vary, and indeed experienced this strongly in the interleaved high-contrast trials. Thus, any variations in difficulty they experienced due to duration, were much more likely attributed to the instructed strength variations than to undisclosed duration variations. The inability to confidently detect evidence onsets would have been aided by the on-off flickering nature of the stimuli, and is confirmed in our results showing longer - not shorter - RTs for shorter evidence durations and sustained elevation of the CPP for even those short durations. This lack of explicit knowledge of variability in evidence duration is what differentiates our task from other variable-duration tasks (Kiani et al., 2008). The fixed and explicitly instructed lead-in period as well as the interleaved easy trials, would have continually reinforced the participants’ ability to time their sampling onset close to evidence onset. This, along with the difficulty of most trials, would have meant that participants were deterred from ignoring any initial evidence, and, indeed, accuracy in even the shortest (0.2 s) condition was reliably above chance (t(15) = 2.60, p = 0.0201). Supporting this, a model that allowed for delayed sampling onset (as well as premature onset) estimated this onset consistently within a few tens of msec of evidence onset, often slightly preceding it (Tables S3, S5-8).

Once the stimulus was removed, a go-cue instructed participants to indicate the higher contrast grating by clicking a mouse button with the thumb of the corresponding hand within 500 ms. The mouse was positioned flat and centrally in front of the participants, with fingers resting flat on the desk. Feedback was presented at the end of each trial in text and audio format. The text feedback informed the subjects whether they responded correctly (‘Correct’), incorrectly (‘Error’), prematurely (responded before the go-cue; ‘Clicked too Early’), or missed the deadline of 500 ms to confirm their choice (‘Too Late’). Although the short deadline was set on the response-cue to encourage the participants to indicate their decision immediately after the stimulus offset, since accuracy was the critical behavioural measure we analysed all responses up to 1 second after the go-cue. 0.4% of trials were missed (no button pressed up to 1 sec) and 2.6 % were premature responses (pressing before go-cue at stimulus offset), and these were excluded from calculating the accuracy. Participants received 10 points for correct, timely trials and 0 points for the other responses. At the end of each block, feedback was indicated on the mean accuracy and cumulative points gained during the block and participants were encouraged to attain as high a score as possible.

### Training

Before performing the main experiment, participants conducted a practice version of the task to become familiar with the experiment and to be able to achieve the desired correct responses within the required deadlines. The training contained a series of short blocks of 20 trials with a fixed contrast difference shown for the full 1600 ms without any duration manipulation. The contrast difference started off with a high contrast difference of ±50%, dropping to ±20% with a 10% reduction in each block. They were required to achieve >90% in the easiest (±50%) and >70% in the hardest conditions (±20%), before progressing to the next difficulty level.

#### Statistical Analysis

Accuracies and response times were measured for each condition based on all trials with a valid response, which excluded 0.4% of trials that were missed (no response before 1s following the go-cue) and 2.6% of premature responses prior to the go-cue. Response times (RT) were recorded from the onset of evidence in milliseconds (ms).

One-way repeated measures analyses of variance (ANOVA) were carried out on the five task conditions, followed by pairwise comparisons (paired-sample t-tests), for RTs, accuracies, and neural amplitudes. For all statistical analyses, prior investigation was carried out to determine whether the data were suitable for parametric analysis by screening for any major violations of normality. Mauchley’s test of sphericity was also conducted to assess the assumption of equal variances of the differences across all condition pairings in the rmANOVAs, and Greenhouse-Geisser corrections were applied when the sphericity assumption was violated.

#### EEG Acquisition and Preprocessing

Continuous EEG was acquired using a 128-channel ActiveTwo system (BioSemi) at a sample rate of 512 Hz. Vertical and horizontal eye movements were monitored using two vertical electrooculogram (EOG) electrodes placed above and below the left eye and two horizontal EOG electrodes placed at the outer canthus of each eye. Customised scripts in MATLAB (Mathworks, Natick, MA) and the EEGLAB toolbox (Delorme & Makeig, 2004) were used to analyse EEG data. EEG data were low-pass filtered by convolution with a 77-tap Hanning-windowed sinc function to give a 3-dB corner frequency of 38 Hz with strong attenuation at the mains frequency (50 Hz) (Widmann et al., 2015). Each block was detrended quadratically. Noisy and flat-lined channels were identified by eye by comparing standard deviations to neighbouring channels and were interpolated based on surrounding channels using spherical splines. No high-pass filter was applied. Initially, EEG data were segmented into stimulus and response-aligned epochs. Stimulus-aligned epochs were extracted from stimulus onset to 960 ms after stimulus offset. All epochs were baseline-corrected relative to the interval of 106.6 ms pre-evidence onset (2 SSVEP cycles). We again applied spherical spline interpolation to any channel with excessively high variance with respect to neighbouring channels in each trial. Trials were rejected if an artifact occurred between evidence onset and stimulus offset (any channel with absolute magnitude >70 µV). One participant was excluded from EEG analyses due to excessive perspiration artifacts (>50% in each condition). Finally, to minimise the effects of volume conduction and avoid the possibility of overlapping signals confounding our interpretation of key signals such as the CPP (Kelly & O’Connell, 2013; Philiastides et al., 2014), a current source density (CSD) transformation was applied to each trial (Kayser & Tenke, 2006). After preprocessing, 12017 trials remained across all conditions, with an average rejection rate of 15% (± 12.9%) across participants.

### Electrophysiological data analysis

We extracted neural signatures of three processing levels of decision making: sensory evidence encoding, accumulation of evidence, and motor execution.

### Neural signature of Sensory Evidence (Steady State Evoked Potential, SSVEP)

Flickering a stimulus at a certain frequency generates an oscillatory signal with the same frequency, known as the steady-state visual evoked potential (SSVEP), which provides a robust representation of physical contrast due to its narrow-band content (Regan, 1989). To obtain a direct measure of the sensory evidence upon which observers based their decisions (i.e. the relative contrast of the overlaid gratings) we generated a difference-SSVEP (d-SSVEP) by flickering the left- and right-tilted gratings in alternation at 18.75 Hz. This produced a signal that was zero, on average, for the equal-contrast baseline stimulus because the alternating gratings cancelled each other. When contrast differed, an aggregate d-SSVEP was produced with a phase dictated by the higher-contrast grating. The d-SSVEP signal was derived for each single trial by computing a Fourier Transform on untapered (boxcar) 320-ms time-windows (exactly 6 SSVEP cycles) in time-steps of 53.33 ms (1 cycle) and extracting the complex value representing oscillatory phase and magnitude at the 18.75-Hz frequency bin. We used the average complex oscillatory 18.75-Hz signal produced by the low-contrast 1600-ms condition to provide a common reference. This reference phase was calculated by trial-averaging the complex signal for the intermediate contrast level (S_ref_). This was done independently for each electrode and time point. The angle of this resulting complex average was _ref_ = arg(S_ref_) then used as the reference phase to which all signal-trial data were aligned using phase rotation. Subsequently, these reference phases were subtracted from all trials, effectively rotating the phase data to correct for arbitrary offsets. One implication of the current approach is that we are unable to assess potential anticipatory modulations preceding the onset of the evidence. This means no reliable phase information is available for the pre-evidence period. However, this has minor concern in the current study, as our primary interest lies in modulations occurring during the evidence presentation. Oz electrode where the d-SSVEP was maximal on grand average was used to calculate this signal. The grand-average topography of the d-SSVEP was calculated by averaging from 300 ms to 1500 ms after stimulus onset (grey shading in Figure 3A) for the low-contrast, 1600-ms condition.

### Neural signature of Evidence Accumulation (Centro Parietal Positivity)

Decision formation dynamics were traced via the centro-parietal positivity (CPP), which has previously been shown to exhibit many properties predicted of evidence accumulation (O’Connell et al., 2012; Twomey et al., 2016). The evidence-locked CPP, baseline-corrected to a 106.6-ms window ending at evidence onset, was measured from a single electrode near standard site Pz (see marker in Fig 3B topographies), where the grand-average, high-contrast CPP was maximal at its peak latency (513 ms). For modelling and visualisation (Fig 3) we smoothed the centroparietal waveform using an additional low-pass filter (convolution with a 110-tap Hanning-windowed sinc function to give a 3-dB corner frequency of 5.05 Hz). This was to reduce the impact of an apparent initial peak and lull during the centroparietal buildup, since all of the models we were considering would assume a unitary build-peak-fall profile. We discuss the potential impact of this initial peak (shown in Figure 3 - Figure Supplement 1) in the Discussion.

### Motor Level decision threshold (Mu-Beta-Band spectral amplitude)

As in previous work (de Lange et al., 2013; Donner et al., 2009; Kelly & O’Connell, 2013; Steinemann et al., 2018), effector-selective motor preparation (Hanks & Summerfield, 2017; Kelly & O’Connell, 2015) was measured in the reduction of amplitude across the Mu/Beta bands (8-30 Hz) over the motor cortex (electrodes C3 and C4). Here, we calculated Mu/Beta amplitude using a short-time Fourier transform with a boxcar window size of 320 ms and time-steps of 53.33 ms, averaging across frequencies in the 8-30 Hz range but omitting 18.75 Hz to exclude the SSVEP. To produce a relative motor preparation signal that would reflect differential evidence for the decision, lateralized Mu/Beta was calculated by subtracting ipsilateral from contralateral amplitude, with respect to the hand corresponding to a correct answer on each trial. In the analysis of Figure 5, we computed average waveforms for correct and error trials separately, for the separate contralateral and ipsilateral signals without subtraction. The mean amplitude of contralateral signals across conditions at the time of response was taken to represent the ‘motor threshold’ (grey line in Figure 5A).

### Model Fits to behaviour only

We explored five decision-making models, of which one was integration, three were extrema detection, and one was a snapshot strategy (Figure 2A). In all models, evidence samples were drawn from a normal distribution with a positive mean (“drift rate”) during the evidence period and zero mean during the equal-contrast periods, with positive/negative values representing samples favouring the correct/incorrect alternative. What differs among these models is how the decision variable was formed from this evidence and used to determine the choice, as explained below. Evidence in all models had a standard deviation of 0.1 arbitrary units, which acted as the common scaling parameter (s = 0.1), i.e. the parameter that sets the scale of the decision variable units, which are otherwise arbitrary. Core free parameters were common across models to enable direct comparisons of their fitting performance. These consisted of a separate “drift rate” for high and low contrast trials, dictating the strength of sensory evidence, and a “bound” parameter to set positive and negative criterion levels that the DV needed to reach to trigger a correct and incorrect decision, respectively.

All models were fit to the accuracies averaged over all participants across the five experiment conditions (one 1600-msec high-contrast condition and four low-contrast conditions with durations of 200, 400, 800 and 1600 ms) by minimising the ^2^ based statistic, G^2^ (Ratcliff & Smith, 2004), described in detail below. The input sensory evidence signal for all models consisted of 1600 ms of Gaussian noise, plus a mean that stepped up and down according to a boxcar function during the periods of evidence to a height determined by the drift rate. To simulate accuracies across the five conditions, Monte-Carlo simulations of such trials were generated for a given set of parameter values and these parameters were optimised to match the empirical accuracies by minimising the objective function (G^2^) using a SIMPLEX algorithm (MATLAB’s *fminsearch* function), generating a new Monte-Carlo simulation at each step of the optimisation process. The G^2^ statistic captured the difference in trial proportions between the real and simulated data for each of the five conditions, and was calculated using the equation:

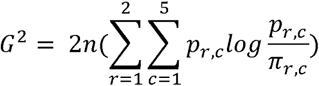

Where n is the number of trials per condition, *p_r,c_* and *π_r,c_*, are the observed and predicted proportions of responses r (correct/error) for each of the five experimental conditions, c. Models with lower G^2^ values fit the data more accurately.

The Akaike Information Criterion (AIC) was used to evaluate the goodness of fit (Kelly et al., 2021) while penalising for model complexity, calculated from G^2^ using the equation:

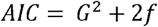

where f is the number of free parameters.

To find the best-fitting parameters for a given model, we took three steps: 1) The G^2^ was evaluated for a large grid of parameter vectors using 1000 simulated trials per parameter set, within the parameter limits listed in Table S4, with 10 points along each parameter dimension; 2) Next, we identified the grid point with the best G^2^ value for each model and ran the SIMPLEX algorithm with 20,000 trials per iteration starting from that parameter vector to refine the parameter estimates. To facilitate the convergence of the SIMPLEX algorithm, the random number generator was set to a fixed seed across all model simulations in a given fit. Then in order to illustrate the degree of uncertainty associated with this monte-carlo simulation approach (Figure 2C), we repeated the fits (step 1 and 2) with 10 different random seeds (i.e. 10 different instantiations of noise). 3) Starting from the parameter vectors attaining the lowest G^2^ in the previous step, the parameter estimates were refined by repeating each instantiation of the SIMPLEX algorithm with 100,000 trials per iteration, with the final parameter estimate and associated G^2^ computed from a new random seed taken as the optimum. This approach starting with a grid search has the advantage of ensuring a comprehensive exploration of the parameter space. We expanded the parameter value limits (Table S4) whenever the fits landed at the outer limit of the existing range. We also verified that the results did not change when we ran the step-2 fits from the 10 best grid points rather than the one best gridpoint with 10 random seeds.

*Integration Models*

We used a simple version of the well-known drift-diffusion model to implement temporal integration (Ratcliff & McKoon, 2008). The samples of momentary evidence were integrated perfectly over time into a decision variable, DV(t), given by:

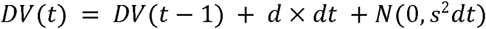

where d(t) is the drift rate at time t that was non-zero only for the duration of evidence, dt is the discrete time increment (set to 2 ms), and the last term refers to Gaussian noise with mean 0 and standard deviation of s×√dt, where s is set to 0.1 as the scaling parameter. This is similar to the EZ-diffusion model (Wagenmakers et al., 2007). Decisions were terminated by reaching the upper/lower bound, or if no bound was reached, determined by the sign of the DV at stimulus offset (Kiani et al, 2008).

Three variants of this model were fitted, the first with only one drift rate applied equally to both high and low contrast-differences to provide a worst-case benchmark; the second with two separate drift rates (UIntg) and no decision bound so that choices were always determined at stimulus offset by the sign of the DV; and the third with a bound parameter included alongside the two drift rates (BIntg).

*Extrema Detection models*

Rather than accumulate evidence samples over time, extrema detection models compared each independent sample of momentary evidence to a criterion or Bound level, terminating upon the first sample to exceed it. The decision variable was thus given by

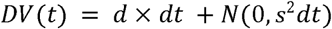

Three variants of this Extrema Detection Model were tested, differing only in how choice was determined when no bound was exceeded within the evidence period: i) by a random (50-50) Guess (labelled ‘ExtG’ for extrema detection with guess default; see Stine et al., 2020), ii) based on the sign of the last evidence sample (‘ExtL’), or iii) based on the sign of the most extreme (sub-bound) sample encountered throughout the evidence period (‘ExtE’). This third variant, labelled ‘Extremum-tracking,’ entailed a memory component to update the extrema as and when they were encountered throughout the trial. Although memory is sometimes associated with integration in terms of potentially shared mechanisms (Wong & Wang, 2006), we class this variant as a non-integration model because the one remembered item (the most recent extremum) is updated sparsely and replaces rather than builds on the previous value, unlike integration which entails continuous, iterative updates. Like the Integration model, we tested three versions of this model: one with one drift rate and no bound, one with two drift rates and no bound, and one with two drift rates and a bound. Unbounded versions of the other two Extrema detection models were not implemented because, in the absence of a bound, they would trivially default to a random guess on all trials, or a decision based on just the last sample, which cannot provide accuracy improvements across the first 3 durations.

#### SnapShot with Guess

In this model, a single, randomly-selected sample of momentary evidence was used to determine the decision. In unbounded variants, the sign of this sample determined the choice regardless of its magnitude. In bounded variants, the sample determined the choice only if it exceeded the bound and a random guess was produced instead when the sample fell short of the bound. Again, three versions of this model were run, respectively with one drift rate and no bound, two drift rates and no bound, and two drift rates with a bound.

### Neurally-constrained Models

The models above were fitted to behaviour alone, and several of them captured accuracy equally well. To resolve this ambiguity, three of these successful models for which a means of producing buildup dynamics in the average decision variable could be conceived - the Integration model and two Extrema Detection models - were subsequently fitted jointly to behaviour and to the average CPP. To do this, a model-based simulation of the CPP was needed to compare to the empirical CPP. This simulation was derived in different ways in each model.

In the Integration model, the CPP was simulated as the absolute value of differential cumulative evidence as previously described (Afacan-Seref et al., 2018). The use of the absolute value operator is based on the assumption that the CPP reflects the sum of two neural populations (one for each choice alternative) that activate in a mutually exclusive way to represent positive and negative values of the single, differential decision variable (Kelly et al., 2021). This is consistent with observations that the CPP builds with positive polarity regardless of choice and that the CPP reaches a stereotyped threshold level at response during urgency-free, continuous monitoring decisions, even though it derives from two accumulator processes only one of which ultimately triggers the decision (Kelly & O’Connell, 2013). Based on post-decision CPP dynamics observed in immediate response tasks and easy delayed-response conditions, we assumed that the decision variable falls back to baseline in a stereotyped way upon reaching the bound. For consistency with the non-integration models below, we modelled this fall-back as a quarter-sine wave (from peak to zero-crossing).

In the Extremum-tracking model, the CPP was simulated again as the absolute value of a differential DV, with the DV in this case being an iteratively-updated representation of the most extreme evidence sample encountered so far. As in the integration model, a quarter-sine fall-down from the bound was assumed if and when it was crossed. In the final model based on Extrema detection with a random Guess default, the decision variable is again the momentary evidence itself, subjected to bounds, but here it is assumed that a centroparietal signal is only evoked upon such bound crossings. We assumed that the trial-averaged CPP was the result of averaging single-trial discrete event waveforms that were triggered by, and therefore time-locked to, decision termination. We modelled this decision termination-flagging event as a half (upper) sinusoidal function whose width is a free parameter and whose amplitude is fixed, starting at the bound crossing time. When averaged, these half-sines can resemble any arbitrary waveform depending on the distribution of bound-crossing times. In theory, this formulation allows the Extrema Guess model to produce a signal with ramp-like properties similar to the CPP without accumulating evidence. For trials where no bound was reached, the CPP was simulated as zero throughout the evidence period. For consistency with this last, ‘Extremum-flagging’ model, the fall-down dynamics of the simulated CPP after each bound-crossing for the Integration and Extremum-tracking models was assumed to take the shape of a quarter-sine function (peak to zero-crossing), whose width was similarly set as a free parameter in the neurally-constrained fits. Note that the ‘FlagWidth’ parameter referred to half of the sinusoidal period across all 3 models for consistency, and so the actual duration of the peak-to-baseline fall-down in the Integration and Extremum-tracking models was half of the estimated value of this parameter. As in the Integration mode, the absolute value of each single-trial DV waveform was taken before averaging in both Extrema detection models.

Based on the above simulations of the CPP, neurally constrained models were implemented by combining the same G^2^ metric quantifying behavioural fit with a metric of CPP fit with variable weighting in a joint objective function to be optimised by the same SIMPLEX algorithm as used previously. Specifically, the joint objective function was given by

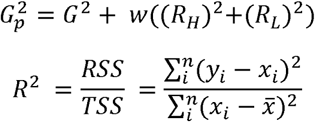

where R^2^ is the residuals sum of squares (RSS) over the total sum of squares (TSS) of the differences between simulated,, and empirical CPP,, waveforms, computed for the high (subscript H) and low (L) contrast conditions. The 4 low-contrast conditions were averaged so that the models would not be informed about amplitude variations across them, allowing these variations to be used as an independent evaluation of model fit (Figure 4-Figure Supplement 2, and Figure Supplement 5 for basic 3-parameter models). The neural weighting was set to each of the values w= 0.1, 1, 10, 100, 1000 so that we could conduct a sensitivity analysis of how model fits varied with the degree of emphasis placed on the neural fit. To compute the neural R^2^ term, we had to first apply appropriate time-windowing to exclude segments of the grand average CPP waveform that did not clearly map onto the decision variable in the model, and normalize the real and simulated CPP so that they would have the same units. Specifically, we extracted windows of the observed CPP waveforms starting from 200 ms, the point at which the low and high-contrast CPP traces begin to diverge and presumably therefore the moment of initial impact of evidence, up to 900 ms in the high-contrast condition and up to 1600 ms in the low contrast conditions. The earlier end time for the high-contrast condition was chosen to avoid the bulk of a separate negative-going potential that appeared in the latter half of the stimulus-evaluation period in that condition, whose relationship to the decision process was not obvious and which would therefore risk biasing the measurement of the CPP during that time window. We then shifted the waveforms down so that they began at zero amplitude and then divided by the mean across all sample points to normalize them. The same normalization procedure was applied to the model-simulated CPPs, which gave the simulated and empirical waveforms comparable units.

In extracting the to-be-modelled CPP waveform starting at the onset of evidence-driven buildup, we remove the need to estimate transmission or other non-decision delays, which had also been unnecessary in behaviour-only modelling due to not fitting reaction time. Without loss of generality, we could therefore assume for simplicity that evidence accumulation and extremum-detection processes began without delay at evidence onset, and that the ‘flag’ signal began immediately upon bound crossing (Figure 4D). It is important to note that this measure to remove unnecessary timing parameter estimation still allows for decision variable onset delays and flag-onset delays that may occur in reality. In all three models, we allowed the width of the sine-function describing the post-bound fall-down (quarter-sine) and flag (half-sine) to vary as a free parameter, on the basis that we cannot estimate these single-trial quantities from observed average waveforms without knowing bound-crossing times.

Although motor preparation signals were not used in the model fits, it was additionally of interest to simulate these signals from all models for qualitative comparison with Mu/Beta lateralisation (Figure 3C). To do this, we assumed that the relative motor preparation directly reflected the differential decision variable but with the distinction that it sustained at the level of the bound upon reaching it, rather than immediately falling like the CPP. From this perspective of motor preparation, the Extremum-flagging model can be considered highly similar in nature to the discrete stepping model proposed by Latimer et al., (2015), which features stochastically occurring, discrete steps in neural activity reflecting transitions from an undecided to a decided state. In Figure 4N-P, to replicate the same temporal blurring that results from the Short-time Fourier Transform (STFT) in the empirical data, we smoothed the simulated motor preparation signals by convolving with a boxcar function of the same duration as the STFT window (320 ms).

Beyond the core parameters of drift rate and bound, sequential sampling models are typically endowed with additional parameters to enable them to achieve a comprehensive behavioural fit (Ratcliff et al., 2016). We therefore explored a range of additional parameters and their impact on the neurally-constrained fits. Specifically, for each of the three model classes - Integration, Extremum-tracking and Extremum-flagging - we included one extra model feature at a time, alongside the two drift rates and bound of the basic model. We describe each additional feature in turn:

### Sampling onset time (SampT)

When sensory evidence is weak, it is difficult to distinguish evidence from noise and this can lead to variability in the onset time of sequential sampling (Devine et al., 2019). Though subtle, the CPP in this task appeared to begin to rise before the onset of evidence, suggesting a potentially early sampling onset (Figure 3B). SampT was a free parameter that was allowed to take a value before or after time zero, which in the models marked the time when informative contrast-difference evidence first became available for sampling. Figure 4 - Figure Supplement 1 shows that this extra parameter did not substantively improve the fit of either of the competitive models.

### Starting point variability (sz)

Variability in the starting point of the decision variable allows for random trial-to-trial fluctuations in bias (Bogacz et al., 2006; Purcell et al., 2012). We implemented starting point variability as a uniform distribution of starting levels centred at the neutral position between the correct/error bounds whose standard deviation was the free parameter sz. Figure 4 - Figure Supplement 1 illustrates that adding this extra parameter substantially improved the fit of the integration model. Interestingly, this nonzero starting-point variability is supported by the motor signal activity at the stimulus onset (Figure 5), although the model was not informed about this empirical effect.

### Time-varying bound (,*t_0.5_*)

While constant bounds offer a parsimonious account of the decision criterion, bounds that change the criterion amount of evidence required for a decision as a function of elapsed time in the trial have been shown to better capture neural and behavioural data in certain cases, such as when urgency to respond is imposed through deadlines (Kelly et al., 2021), and although responses were deferred until after the long stimulus in the current task, a tight deadline of 0.5 sec was imposed at that point, which in combination with the predominant very weak evidence may have led to an experience of urgency. Such dynamically changing bounds have been shown to be optimal in certain circumstances involving unpredictable task difficulty (Malhotra et al., 2018) and are a critical feature of some extrema detection models (Thura et al., 2012). To determine whether models with time-varying bounds provide better fits to the data than models with fixed bounds, we implemented a time-varying bound using a linear function, given by

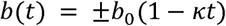

where *b_0_* is the initial boundary separation, time (t), *k* represents the collapse rate, allowed to be positive or negative (increasing bound). We also fit an alternative, nonlinear collapsing bound model with a hyperbolic ratio function, given by:

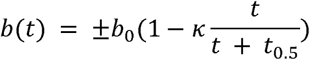

where *t_0.5_* the semi-saturation time. The positive and negative versions of this bound function correspond to the correct and incorrect alternative. This temporal function has been previously used to quantify the hypothetical urgency signal in physiological data (Churchland et al., 2008; Hanks et al., 2011; Voskuilen et al., 2016), except that the amount of collapse is expressed as a fraction of the total boundary separation rather than as an absolute value, which simplified fitting and led to more interpretable parameter estimates. Our results indicate that incorporating time-varying bounds improved the model fits for all the models (Figure 4 - Figure Supplement 1). The optimal bound change for the two extrema-detection models was always a decreasing one (positive values of *k*; Table S3&S4).

### Between trial drift rate variability (eta)

Although our stimulus does not include fluctuations in contrast, the internal representation of this contrast may fluctuate from trial to trial, with factors such as alertness and attention (Ratcliff et al., 2016). To allow for such between-trial variability in encoding sensory information, an extra parameter for trial-to-trial variability in drift rate was added, captured as Gaussian variability with standard deviation eta, centred on the drift rate parameters. This additional parameter improved the fit for integration model slightly (lower G^2^+Penalty) while did not improve the fit for Extremum-flagging model (Tables S3, S5-8).

### Accumulation duration (Dur)

A central question in this study was whether judgements about prolonged static sensory evidence involve sequential sampling over the full evidence period, or only a portion of it. To explore this possibility in all three models, we implemented the models with one extra parameter (‘Dur’) that determined the time window starting from the onset of evidence encoding during which evidence was sequentially sampled for decision formation. The results revealed that the basic Integration model did improve with the addition of this duration parameter, but the estimate of this duration needed to fit the neural dynamics was less than 800 ms, therefore precluding reproduction of the significant increase in behavioural accuracy from the 800-ms to 1600-ms duration condition (Tables S3, S5-8).

### Leaky accumulation

Another way that integration of evidence can be limited is through leak, whereby older samples are discounted in the cumulative total, and the decision variable gradually decays to zero in the absence of sufficient evidence. We implemented leaky accumulation through an additional leak parameter which scales-down the weighting of previous evidence relative to the current sample, according to:

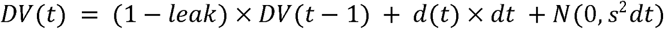

The results demonstrated minimal information loss over time (leak = 0.0002 at w=10), with only a small benefit to fit from including the parameter, in agreement with previous findings of low leakage (Kiani et al., 2008).

The fitting procedure for the neurally-constrained models followed the same general strategy as the model fits to behaviour alone, starting with evaluation of objective function at each point of a comprehensive grid, but now running the SIMPLEX algorithm from the 100 best grid points as starting parameter vectors. In the most complex model, which extends the basic model by including starting point variability and a non-linear collapsing decision bound, five grid points were used for each of three parameters (starting point variability, collapse rate and semi-saturation time) instead of 10 to limit computational expense (see Table S4). Running this procedure on all three model classes, we found that whereas the best objective function values for the Integration and Extremum-flagging models were 7.98 and 9.95, respectively (taking w=10 as representative), the best achieved by Extremum-tracking was 15.83. We therefore set Extremum-tracking aside and proceeded to refine the two competitive models, the best performing of which included both collapsing bound and starting point variability. To further ensure convergence on global optima despite the additional complexity, we ran additional fits with starting vectors generated from the parameter estimates of models that had fewer free parameters. Specifically, for the ‘szC’ models (with free parameters for both collapsing bound and starting point variability), we started a fit from the best fitting ‘C’ model and with sz set to zero, half of and equal to its estimated value in the ‘sz’ model. Analogously, we generated two more ‘szC’ starting vectors by taking the best fitting ‘sz’ model parameter values and adding an additional collapse rate equal to zero, equal to half and equal to the collapse rate found in the ‘C’ model.

## Supplementary Information

**Table S1.** Pairwise comparisons of accuracy, showing a significant difference between each pair of consecutive durations

| Duration 1 | Duration 2 | T | dof | p-value (uncorrected) | BF10 |
| --- | --- | --- | --- | --- | --- |
| 0.2s | 0.4s | -3.78647 | 15 | $1.79 \times 10^{-3}$ | 23.199 |
| 0.4s | 0.8s | -3.74 | 15 | $1.97 \times 10^{-3}$ | 21.39 |
| 0.8s | 1.6s | -6.19855 | 15 | $1.70 \times 10^{-5}$ | 1372 |

**Table S2.** Pairwise comparisons of mean RT. There was a significant difference in reaction time between each pair of consecutive durations except for the shortest durations (0.2s vs 0.4s).

| Duration 1 | Duration 2 | T | dof | p-value (uncorrected) | BF10 |
| --- | --- | --- | --- | --- | --- |
| 0.2s | 0.4s | 1.40869 | 15 | $1.8 \times 10^{-1}$ | 0.587 |
| 0.4s | 0.8s | 17.8368 | 15 | $1.60 \times 10^{-11}$ | $4.41 \times 10^8$ |
| 0.8s | 1.6s | 2.49377 | 15 | $2.48 \times 10^{-2}$ | 2.62 |

**Table 1 - Table Supplement 1:** the mean and standard deviation parameter estimates of models across the 10 fits to behaviour only. ‘UIntg’ refers to the unbounded Integration model, ‘BIntg’ refers to the bounded Integration model, ‘UExtE’ refers to the unbounded Extremum-tracking, ‘BExtE’ refers to Extremum-tracking, and ‘BExtG’ refers to Extremum-flagging which has a default guess. ‘B’ refers to the bound and D1 and D2 are the drift rates in the low- and high-contrast conditions, respectively. The parameter estimates tend to vary by no more than 2-5% except for the bound estimates in the Snapshot and Extremum-tracking models, in line with their inability to distinguish whether there is a bound or not.

|  | **UIntg** | **BIntg** | **UExtE** | **BExtE** | **BExtG** | **BExtL** | **BSpst** |
| --- | --- | --- | --- | --- | --- | --- | --- |
| **B** | - | 0.13±0.0058 | - | 0.02±0.006 | 0.03±0.003 | 0.0143±0.0001 | 0.0027±0.0035 |
| **D1** | 0.056±0.0007 | 0.056±0.001 | 0.379±0.0013 | 0.380±0.009 | 6.8±1.36 | 0.47±0.017 | 2.38±1.12 |
| **D2** | 0.206±0.0011 | 0.22±0.006 | 1.61±0.0091 | 1.63±0.05 | 9.6±1.4 | 3.64±2.4 | 7.58±2.19 |

**Table S3.** Neurally constrained Integration, Extremum-tracking and Extremum-flagging models with various extra model features at neural constraint weighting **w = 10**. ‘BIntg’ refers to the bounded Integration model, ‘BExtE’ refers to Extremum-tracking, and ‘BExtG’ refers to Extremum-flagging which has a default guess. ‘FlagWidth’ refers to the width of half-sine while t05 and K are semi-saturation constant and collapsing rate parameters of the collapsing bound function.

| w = 10 | G<sup>2</sup>+Penalty | B | D1 | D2 | FlagWidth | extraParam | t05, K |
| --- | --- | --- | --- | --- | --- | --- | --- |
| **Blntg** | 7.9864 | 0.3591 | 0.0566 | 1.0991 | 0.6688 | - | - |
| **Blntg_eta** | 7.6351 | 0.3631 | 0.0641 | 1.1498 | 0.7281 | eta = 0.0513 | - |
| **Blntg_sz** | 2.6871 | 0.2988 | 0.1056 | 1.3391 | 0.7507 | sz = 0.5987 | - |
| **Blntg_Dur** | 7.1693 | 0.3374 | 0.0593 | 1.0801 | 0.7441 | Dur = 1.1515 | - |
| **Blntg_sampT** | 7.9435 | 0.3724 | 0.0593 | 1.0998 | 0.6338 | sampT = -0.0873 | - |
| **Blntg_C** | 5.8835 | 0.3578 | 0.0578 | 0.9601 | 0.7161 | - | K = 0.4594 |
| **Blntg_szC** | 2.2533 | 0.4683 | 0.1607 | 1.4049 | 0.2502 | sz = 0.9419 | K = 0.312 |
| **Blntg_szC (NL)** | 2.5622 | 0.2996 | 0.1133 | 1.2131 | 0.6619 | sz = 0.6 | t0.5 = 3.9996,<br>K = 0.8766 |
| **Blntg_Leak** | 7.8308 | 0.3159 | 0.0577 | 1.0194 | 0.7582 | Leak = 0.0002 | - |
| **BExtE** | 19.7663 | 0.035 | 0.3815 | 10.865 | 0.9 | - | - |
| **BExtE_szC** | 15.836 | 0.059 | 0.384 | 16.79 | 0.1 | sz = 0.0015 | K = 0.46 |
| **BExtE_szC(NL)** | 17.869 | 0.04 | 0.3808 | 12.3183 | 0.89 | sz = 0.00002 | t0.5 = 2.4969,<br>K = 0.9101 |
| **BExtG** | 9.9581 | 0.0145 | 0.5079 | 2.0663 | 0.52 | - | - |
| **BExtG_eta** | 10.093 | 0.0143 | 0.4666 | 2.2095 | 0.5705 | eta = 0.0829 | - |
| **BExtG_sz** | 10.2583 | 0.0142 | 0.4742 | 10.173 | 0.7305 | sz =0.0004 | - |
| **BExtG_Dur** | 10.2672 | 0.0142 | 0.4564 | 15.3698 | 0.7231 | Dur = 1.5716 | - |
| **BExtG_sampT** | 5.7477 | 0.0143 | 0.4909 | 15.7325 | 0.7215 | sampT =0.0119 | - |
| **BExtG_C** | 2.877 | 0.015 | 0.4609 | 2.1092 | 0.4379 | - | K = 0.0889 |
| **BExtG_szC** | 2.7589 | 0.015 | 0.4387 | 2.1614 | 0.4462 | sz = 0.000135 | K = 0.1062 |
| **BExtG_szC(NL)** | 2.9558 | 0.015 | 0.4471 | 2.1584 | 0.4507 | sz =0.0005 | t05 = 2.1641;<br>K=0.3177 |
| **BExtL_szC** | 2.9578 | 0.0149 | 0.416 | 2.102 | 0.42 | Sz = 0.00036 | K = 0.109 |

**Table S4.** Allowed parameter ranges for model fits, i.e. the minimum and maximum values that each parameter can take in the SIMPLEX (fminsearch) algorithm. ‘Default value’ indicates parameter settings used when the parameter is fixed rather than free. An asterisk * marks the parameters with 5 grid points in the models with both starting point variability and a collapsing bound.

| Parameters | Label | Integration | ExtE-tracking | ExtG-flagging | Default value |
| --- | --- | --- | --- | --- | --- |
| bound | B | [0, 0.65] | [0, 0.06] | [0, 0.04] | - |
| Drift rate (Low) | D1 | [0, 0.6] | [0, 2] | [0, 5] | - |
| Drift rate (High) | D2 | [0, 2] | [0, 20] | [0, 15] | - |
| Between trial Variability | eta | [0, 0.5] | [0, 0.5] | [0, 0.5] | 0 |
| Starting point variability* | sz | [0, 1] | [0, 0.04] | [0, 0.04] | 0 |
| Duration | Dur | [0.2, 1.6] | [0.2, 1.6] | [0.2, 1.6] | 1.6 |
| Accumulation onset Time | sampT | [-0.4, 0.4] | [-0.4, 0.4] | [-0.4, 0.4] | 0 |
| collapse* | k | [-1 1] | [-1, 1] | [-1 1] | 0 |
| Leak | leak | [0, 1] | - | - | 0 |
| FlagWidth | wd | [0.002, 0.9] | [0.002, 0.9] | [0.002, 0.9] | - |
| Semi-saturation time | t05 | [0.1 4] | [0.1 2.5] | [0.1 2.5] | - |

**Table S5.** Neurally constrained Integration, Extremum-tracking and Extremum-flagging models with one extra feature at neural constraint weighting **w = 0.1**. ‘BIntg’ refers to the bounded Integration model, ‘BExtE’ refers to Extremum-tracking, and ‘BExtG’ refers to Extremum-flagging which has a guess default. ‘FlagWidth’ refers to the width of half-sine. t05 and K are semi-saturation constant and collapsing rate parameters of the nonlinear collapsing bound function, whereas the linear collapsing bound function only has a slope parameter K.

|  |  |  |  |  |  |  |  |
| --- | --- | --- | --- | --- | --- | --- | --- |
| $w = 0.1$ | $G^2 + \text{Penalty}$ | B | D1 | D2 | FlagWidth | extraParam | t05, K |
| **Blntg** | 0.3614 | 0.1318 | 0.0566 | 0.2228 | 0.857 | - | - |
| **Blntg_eta** | 0.3404 | 0.2053 | 0.0674 | 0.2642 | 0.8895 | eta = 0.0609 | - |
| **Blntg_sz** | 0.3352 | 0.1443 | 0.0668 | 0.3486 | 0.8923 | sz =0.2108 | - |
| **Blntg_Dur** | 0.3441 | 0.1725 | 0.0591 | 0.2304 | 0.4564 | Dur = 1.3034 | - |
| **Blntg_sampT** | 0.3755 | 0.2453 | 0.0634 | 0.2423 | 0.5171 | sampT=-<br>0.2244 | - |
| **Blntg_C** | 0.3793 | 0.422 | 0.058 | 0.2233 | 0.8931 | - | K = 0.6866 |
| **Blntg_szC** | 0.2328 | 0.3137 | 0.1288 | 0.8927 | 0.8472 | sz = 0.6177 | K = 0.6355 |
| **Blntg_szC (NL)** | 0.1662 | 0.2888 | 0.1221 | 1.3666 | 0.8777 | sz = 0.5784 | t05 = 2.438; K<br>= 2.0444 |
| **Blntg_Leak** | 0.3666 | 0.1291 | 0.0561 | 0.222 | 0.8295 | leak = 0.0 | - |
| **BExtE** | 0.8374 | 0.0347 | 0.3883 | 1.6309 | 0.0044 | - | - |
| **BExtE_szC** | 0.44 | 0.0442 | 0.598 | 4.41 | 0.336 | sz = 0.044 | K = 0.14 |
| **BExtE_szC (NL)** | 0.7045 | 0.0257 | 0.408 | 1.9322 | 0.8911 | sz = 0.0315 | t05=2.156 ;K<br>= -0.6199 |
| **BExtG** | 0.3203 | 0.0155 | 0.8372 | 2.1473 | 0.5367 | - | - |
| **BExtG_eta** | 0.5129 | 0.0195 | 2.7719 | 4.1325 | 0.6719 | eta =0.0996 | - |
| **BExtG_sz** | 0.4193 | 0.015 | 0.6899 | 2.3364 | 0.7015 | sz = 0.0029 | - |
| **BExtG_Dur** | 0.5397 | 0.0195 | 2.8073 | 4.1856 | 0.5624 | Dur = 1.5007 | - |
| **BExtG_sampT** | 0.3384 | 0.016 | 1.0522 | 4.6706 | 0.7095 | sampT = -<br>0.0431 | - |
| **BExtG_C** | 0.2967 | 0.0145 | 0.4045 | 1.8311 | 0.672 | - | K = 0.1044 |
| **BExtG_szC** | 0.2158 | 0.0147 | 0.4033 | 1.7206 | 0.414 | sz = 0.0001 | K = 0.1274 |
| **BExtG_szC (NL)** | 0.2603 | 0.0151 | 0.5023 | 2.0497 | 0.4981 | sz = 0.0029 | t05 = 2.3537;<br>K = 0.5122 |

**Table S6.** Neurally constrained Integration, Extremum-tracking and Extremum-flagging models with one extra feature at neural constraint weighting **w = 1**. ‘BIntg’ refers to the bounded Integration model, ‘BExtE’ refers to Extremum-tracking, and ‘BExtG’ refers to Extremum-flagging which has a default guess. ‘FlagWidth’ refers to the width of half-sine while t05 and K are semi-saturation constant and collapsing rate parameters of the collapsing bound function.

| **w = 1** | **G<sup>2</sup>+Penalty** | **B** | **D1** | **D2** | **FlagWidth** | **ExtraParam** | **t05, K** |
| --- | --- | --- | --- | --- | --- | --- | --- |
| **Blntg** | 2.6731 | 0.1337 | 0.0559 | 0.2232 | 0.8663 | - | - |
| **Blntg_eta** | 2.3237 | 0.1783 | 0.069 | 0.2741 | 0.6237 | eta = 0.0635 | - |
| **Blntg_sz** | 0.6277 | 0.1893 | 0.085 | 0.9504 | 0.8973 | sz = 0.3719 | - |
| **Blntg_Dur** | 2.5003 | 0.1621 | 0.059 | 0.2358 | 0.5866 | Dur = 1.2735 | - |
| **Blntg_sampT** | 2.2597 | 0.1901 | 0.0686 | 0.6587 | 0.8998 | sampT= -0.3973 | - |
| **Blntg_C** | 2.6294 | 0.1342 | 0.0564 | 0.2284 | 0.8679 | - | K = 0.009 |
| **Blntg_szC** | 0.5706 | 0.2293 | 0.0939 | 0.9363 | 0.751 | sz = 0.4533 | K = 0.2339 |
| **Blntg_szC(NL)** | 0.598 | 0.2869 | 0.1093 | 1.1275 | 0.6546 | sz = 0.5732 | t05= 3.7864 ;<br>K =1.4184 |
| **Blntg_Leak** | 2.592 | 0.1367 | 0.0563 | 0.2234 | 0.851 | Leak = 0.0001 | - |
| **BExtE** | 4.4127 | 0.0344 | 0.3874 | 1.6424 | 0.1795 | - | - |
| **BExtE_szC** | 2.69 | 0.07 | 0.59 | 4.82 | 0.47 | sz = 0.04 | K = 0.3 |
| **BExtE_szC (NL)** | 2.0551 | 0.0395 | 0.5155 | 11.0308 | 0.8852 | sz = 0.0397 | t05 = 1.8842;<br>K = -1.8603 |
| **BExtG** | 1.2939 | 0.015 | 0.6599 | 1.959 | 0.5004 | - | - |
| **BExtG_eta** | 3.8019 | 0.0195 | 2.7484 | 4.5935 | 0.5871 | eta = 0.4978 | - |
| **BExtG_sz** | 2.7936 | 0.0171 | 1.5133 | 4.5796 | 0.8999 | sz = 0.0093 | - |
| **BExtG_Dur** | 2.1145 | 0.0159 | 1.037 | 2.37 | 0.5268 | Dur = 1.4749 | - |
| **BExtG_sampT** | 1.041 | 0.0144 | 0.531 | 12.6837 | 0.7459 | sampT = -0.0118 | - |
| **BExtG_C** | 0.6235 | 0.0147 | 0.4147 | 1.8688 | 0.4494 | - | K = 0.1042 |
| **BExtG_szC** | 0.6313 | 0.0146 | 0.4185 | 1.8496 | 0.4559 | sz = 0.00014 | K = 0.0983 |
| **BExtG_szC (NL)** | 1.0876 | 0.0154 | 0.6616 | 2.2876 | 0.4427 | sz = 0.0042 | t05 = 2.4824;<br>K = 0.3838 |

**Table S7.**
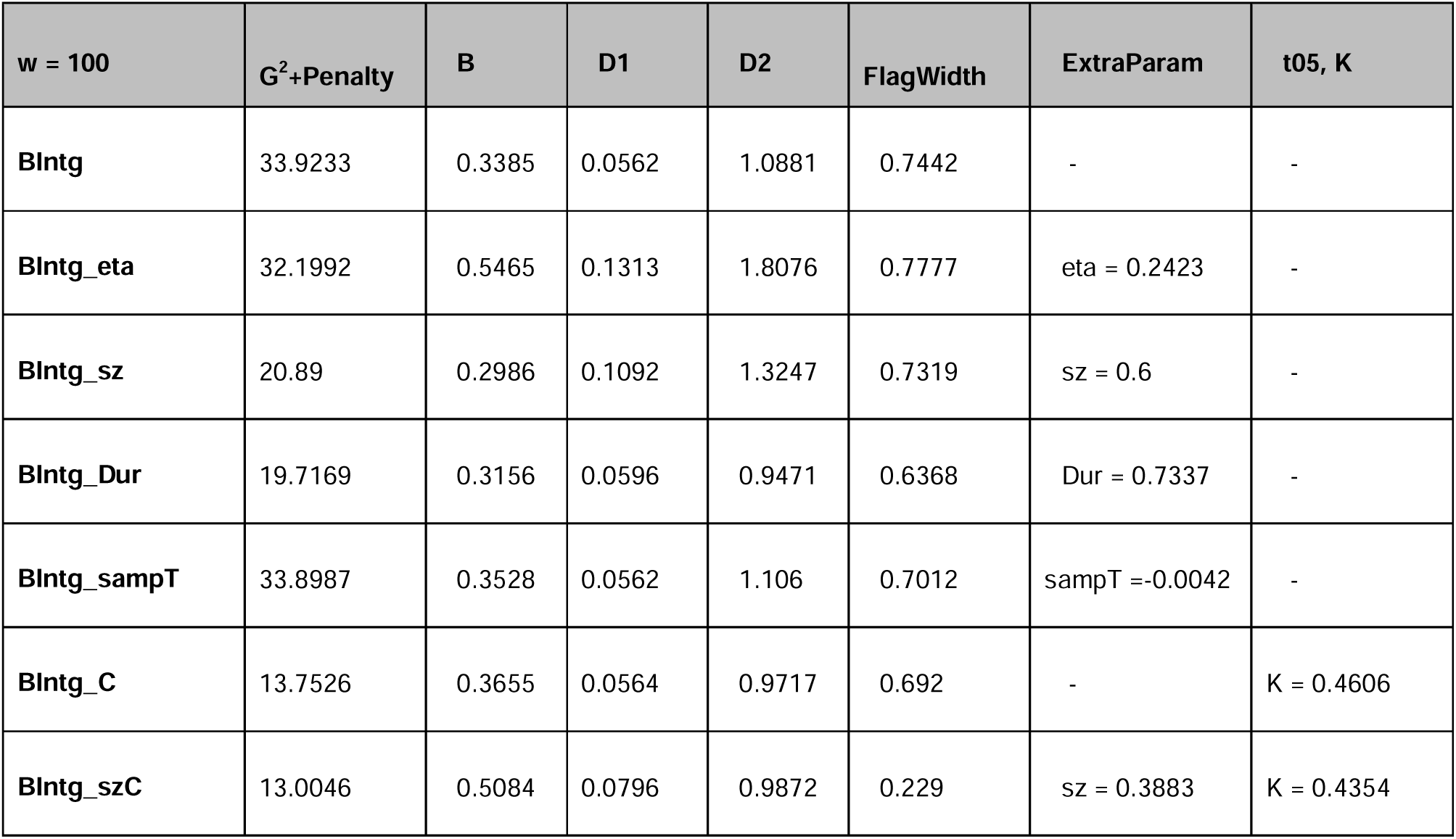
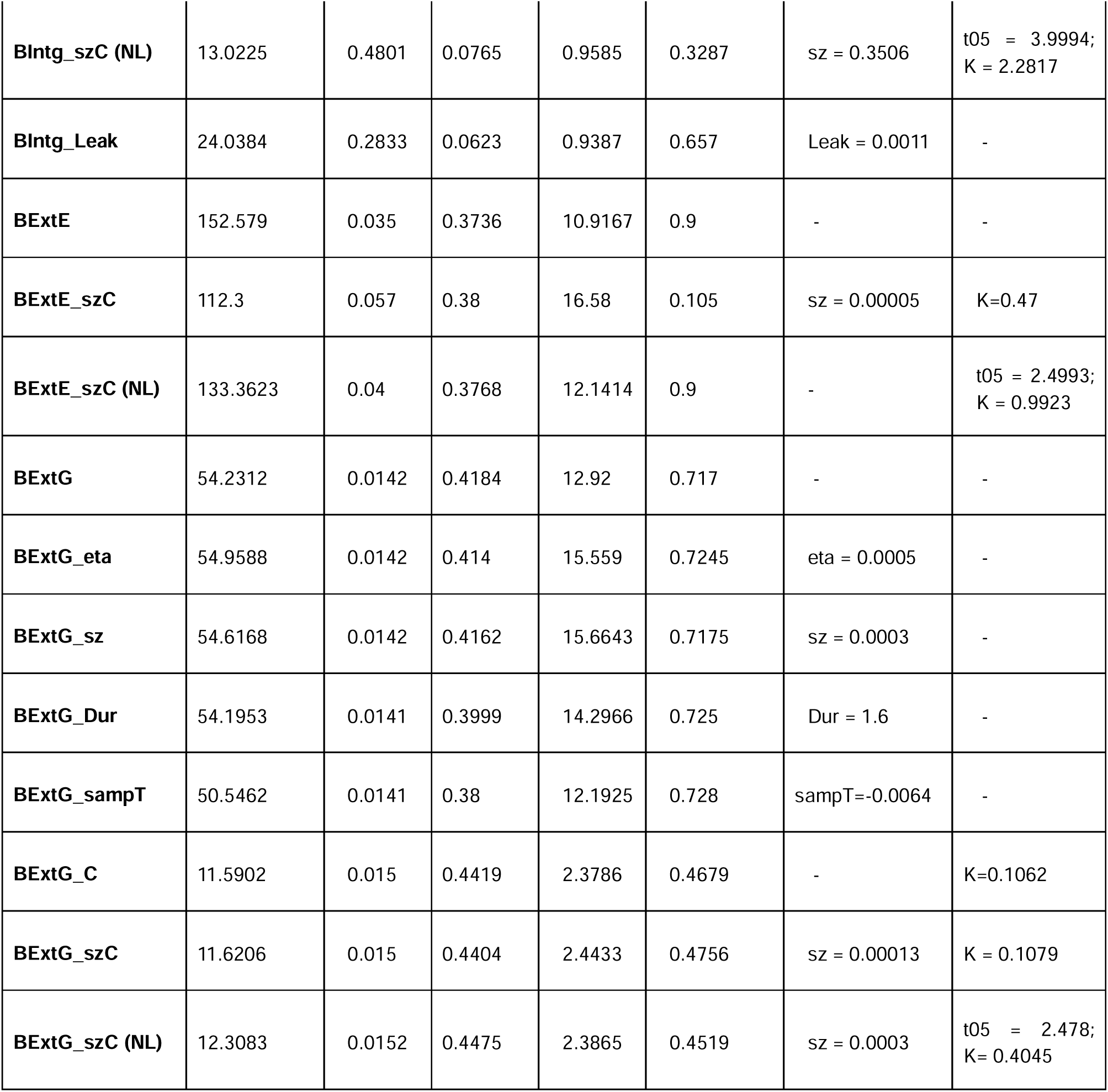
Neurally constrained Integration, Extremum-tracking and Extremum-flagging models with one extra feature at neural constraint weighting **w = 100**. ‘BIntg’ refers to the bounded Integration model, ‘BExtE’ refers to Extremum-tracking, and ‘BExtG’ refers to Extremum-flagging which has a guess default. ‘FlagWidth’ refers to the width of half-sine while t05 and K are semi-saturation constant and collapsing rate parameters of the collapsing bound function.

| **w = 100** | **G<sup>2</sup>+Penalty** | **B** | **D1** | **D2** | **FlagWidth** | **ExtraParam** | **t05, K** |
| --- | --- | --- | --- | --- | --- | --- | --- |
| **Blntg** | 33.9233 | 0.3385 | 0.0562 | 1.0881 | 0.7442 | - | - |
| **Blntg_eta** | 32.1992 | 0.5465 | 0.1313 | 1.8076 | 0.7777 | eta = 0.2423 | - |
| **Blntg_sz** | 20.89 | 0.2986 | 0.1092 | 1.3247 | 0.7319 | sz = 0.6 | - |
| **Blntg_Dur** | 19.7169 | 0.3156 | 0.0596 | 0.9471 | 0.6368 | Dur = 0.7337 | - |
| **Blntg_sampT** | 33.8987 | 0.3528 | 0.0562 | 1.106 | 0.7012 | sampT = -0.0042 | - |
| **Blntg_C** | 13.7526 | 0.3655 | 0.0564 | 0.9717 | 0.692 | - | K = 0.4606 |
| **Blntg_szC** | 13.0046 | 0.5084 | 0.0796 | 0.9872 | 0.229 | sz = 0.3883 | K = 0.4354 |
| **BIntg_szC (NL)** | 13.0225 | 0.4801 | 0.0765 | 0.9585 | 0.3287 | sz = 0.3506 | t05 = 3.9994;<br>K = 2.2817 |
| **BIntg_Leak** | 24.0384 | 0.2833 | 0.0623 | 0.9387 | 0.657 | Leak = 0.0011 | - |
| **BExtE** | 152.579 | 0.035 | 0.3736 | 10.9167 | 0.9 | - | - |
| **BExtE_szC** | 112.3 | 0.057 | 0.38 | 16.58 | 0.105 | sz = 0.00005 | K=0.47 |
| **BExtE_szC (NL)** | 133.3623 | 0.04 | 0.3768 | 12.1414 | 0.9 | - | t05 = 2.4993;<br>K = 0.9923 |
| **BExtG** | 54.2312 | 0.0142 | 0.4184 | 12.92 | 0.717 | - | - |
| **BExtG_eta** | 54.9588 | 0.0142 | 0.414 | 15.559 | 0.7245 | eta = 0.0005 | - |
| **BExtG_sz** | 54.6168 | 0.0142 | 0.4162 | 15.6643 | 0.7175 | sz = 0.0003 | - |
| **BExtG_Dur** | 54.1953 | 0.0141 | 0.3999 | 14.2966 | 0.725 | Dur = 1.6 | - |
| **BExtG_sampT** | 50.5462 | 0.0141 | 0.38 | 12.1925 | 0.728 | sampT=-0.0064 | - |
| **BExtG_C** | 11.5902 | 0.015 | 0.4419 | 2.3786 | 0.4679 | - | K=0.1062 |
| **BExtG_szC** | 11.6206 | 0.015 | 0.4404 | 2.4433 | 0.4756 | sz = 0.00013 | K = 0.1079 |
| **BExtG_szC (NL)** | 12.3083 | 0.0152 | 0.4475 | 2.3865 | 0.4519 | sz = 0.0003 | t05 = 2.478;<br>K = 0.4045 |

**Table S8.**
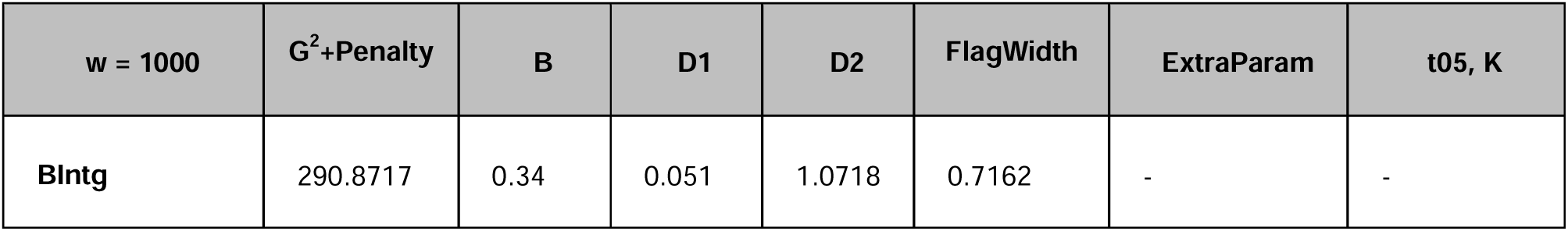
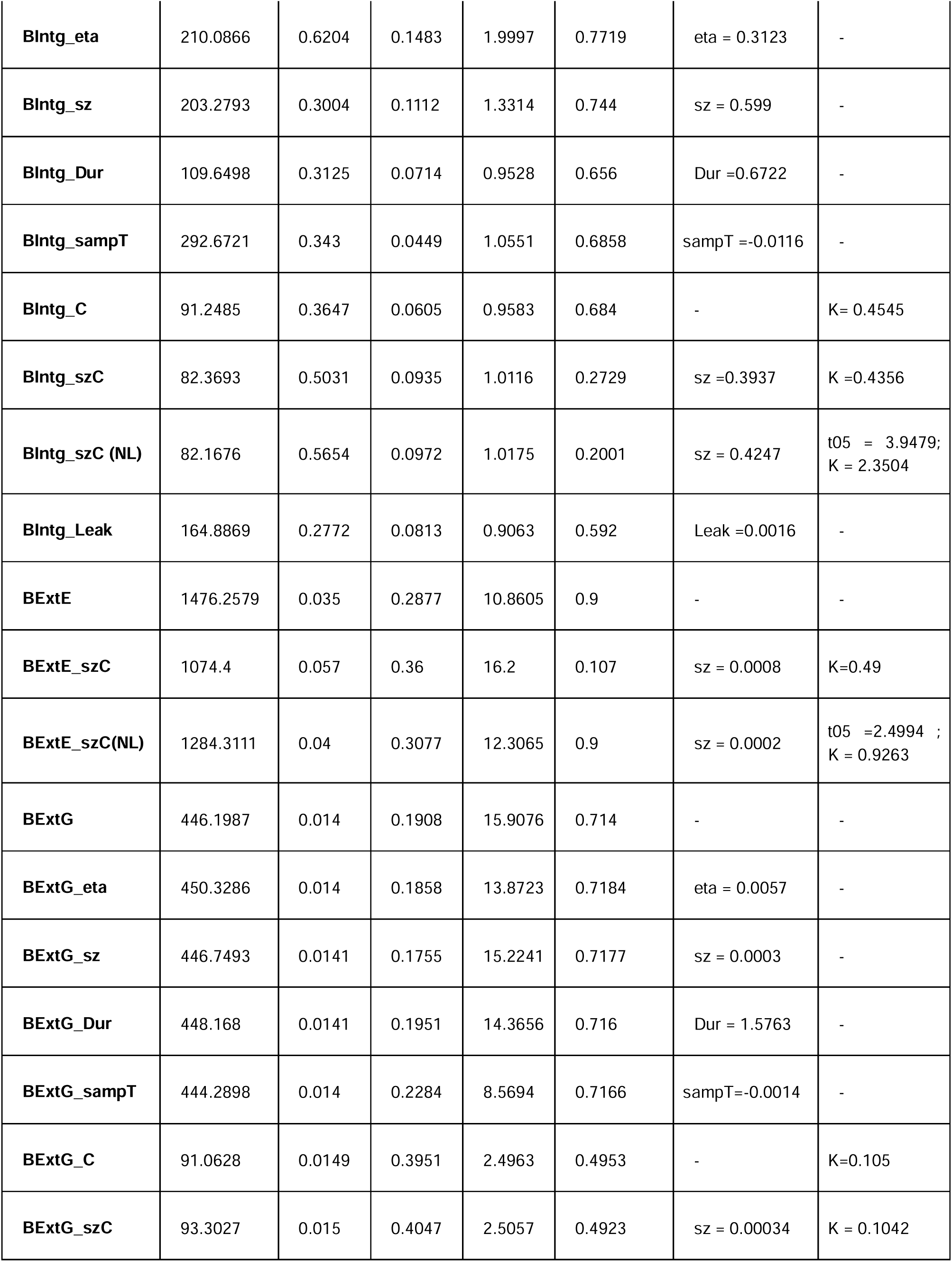
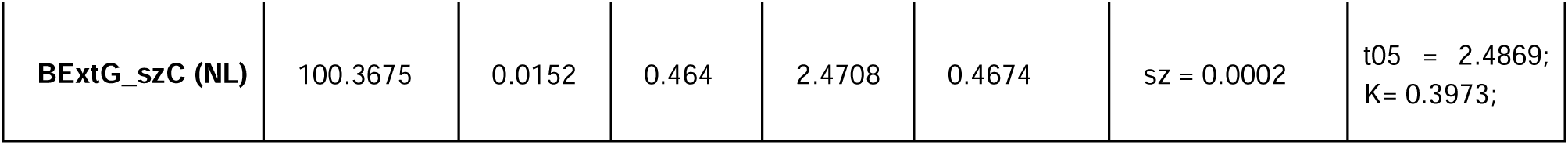
Neurally constrained Integration, Extremum-tracking and Extremum-flagging models with one extra feature at neural constraint weighting **w = 1000**. ‘BIntg’ refers to the bounded Integration model, ‘BExtE’ refers to Extremum-tracking, and ‘BExtG’ refers to Extremum-flagging which has a guess default. ‘FlagWidth’ refers to the width of half-sine while t05 and K are semi-saturation constant and collapsing rate parameters of the collapsing bound function.

**Figure 2 - Figure supplement 1.**
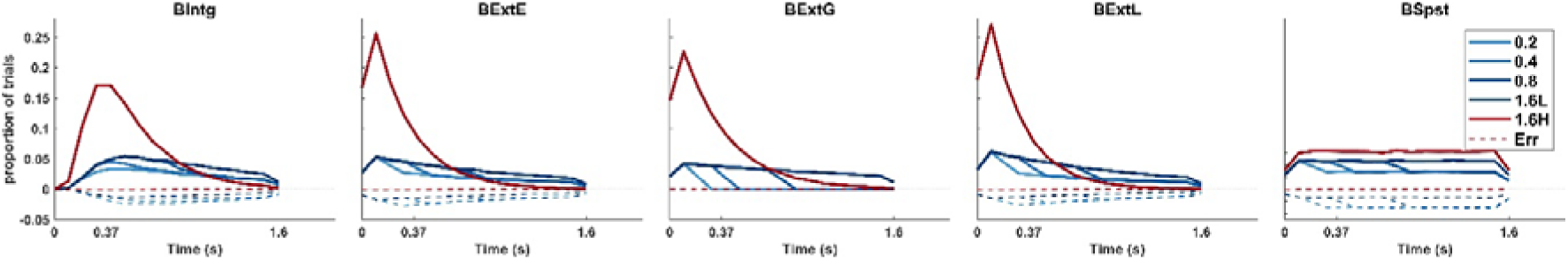
Histograms of Bound-crossing times per condition in behaviour-only models that included a bound. The integration model, **BIntg**, allows 55-60% of trials to exceed the bound in the low-contrast conditions and nearly all trials in the high-contrast conditions, with the relative proportion of correct vs error bound crossing scaling with duration. A similar pattern is observed in the **BExtE** and **BExtL** extrema detection models. The **BExtG** model produces a good fit by setting a relatively high bound and assuming a drift rate for low-contrast conditions that is almost as high as the high-contrast condition. This produces no error bound crossings and an almost uniform distribution of correct bound crossing times during the evidence periods for hard trials, which produces a linearly increasing ratio of correct bound crossings to guesses across durations, in turn yielding a linear increase in accuracies. The Snapshot model, **BSpt**, by construction, produces a bound crossing distribution that is uniform across all time for which there is evidence.

**Figure 2 - Figure Supplement 2.**
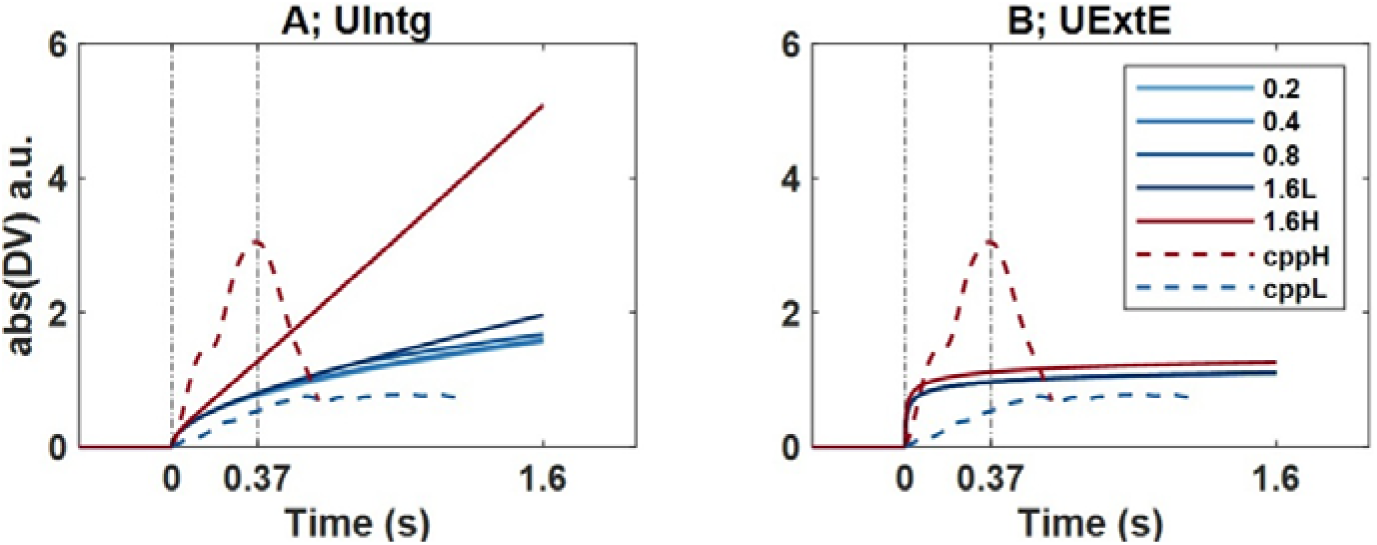
Simulated decision variable waveforms of the unbounded integration (A; UIntg) and non-integration (B; UExtE) models that produced good fits to behaviour alone, illustrating that neither of these models can reproduce the peak and falling-down of the empirical CPP signal (dashed) in the high-contrast condition. For consistency with the neurally-constrained modelling analysis, empirical traces show the mean CPP segments that are used as constraints, with low-contrast conditions averaged (‘cppL’; See Neurally-constrained model-fitting in Results), whereas all five conditions are simulated for the models.

**Figure 3 - Figure Supplement 1.**
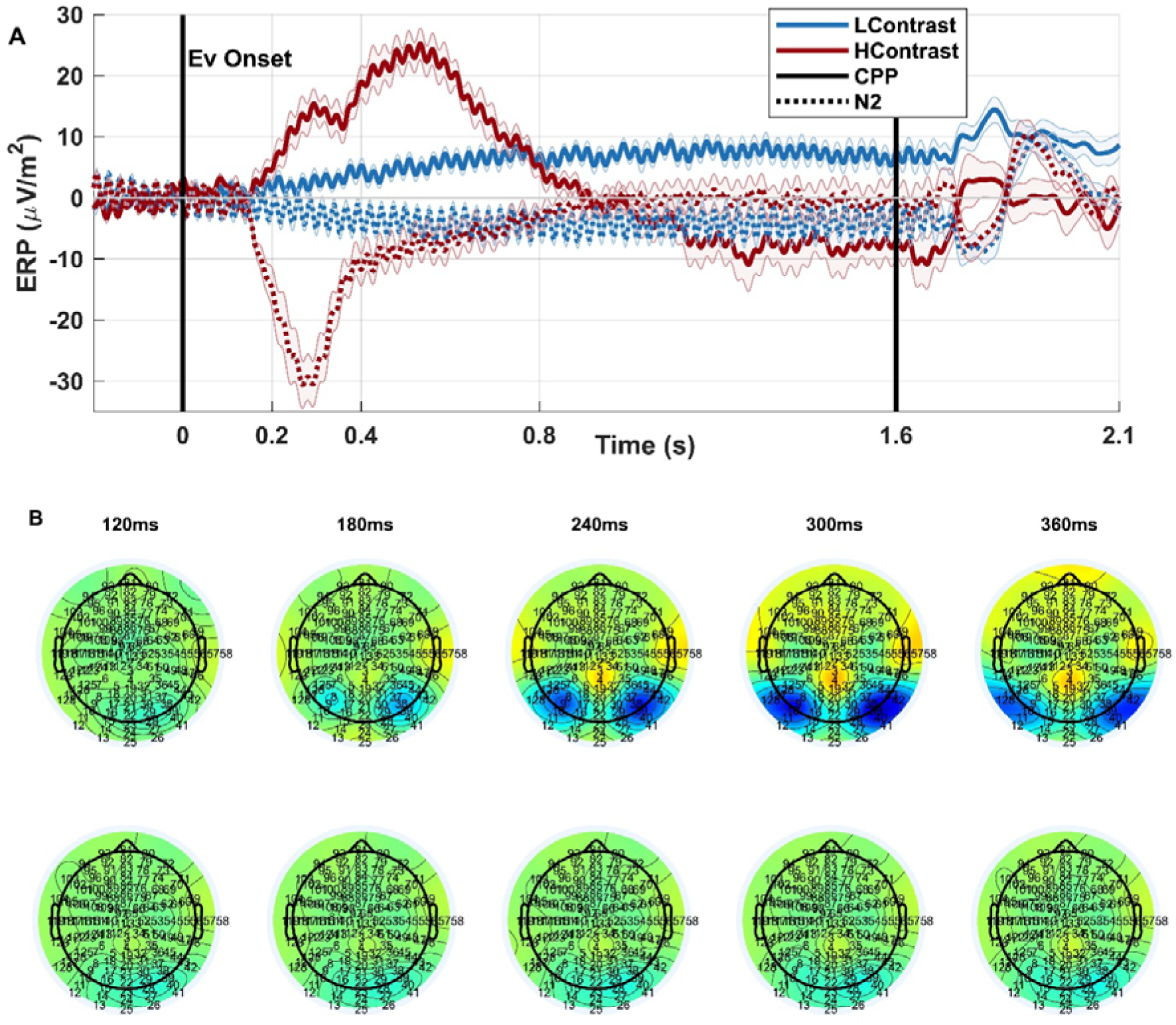
(A) Centroparietal positivity (CPP) waveform without the 5-Hz cutoff smoothing filter applied as in Fig 3 and Fig 4, demonstrating a bimodal morphology that is not present in the low-contrast conditions (shown here averaged across durations). For comparison, the waveforms from bilateral occipital sites are superimposed in dashed lines, showing that the initial peak in centroparietal buildup is contemporaneous with the peak of the N2 component. We verified that this pattern was visible in several individual subjects. (B) Series of topographies from the onset of activity through the dip in centroparietal buildup, for the high-contrast (top) and averaged low-contrast conditions (bottom), again indicating that the initial peak and dip of the high-contrast centroparietal waveform likely contains a contribution from the positive end of dipolar activity generating the occipital N2. Each labelled timepoint represents the centre of a 60-ms window across which the amplitude is averaged.

**Figure 4 - Figure Supplement 1.**
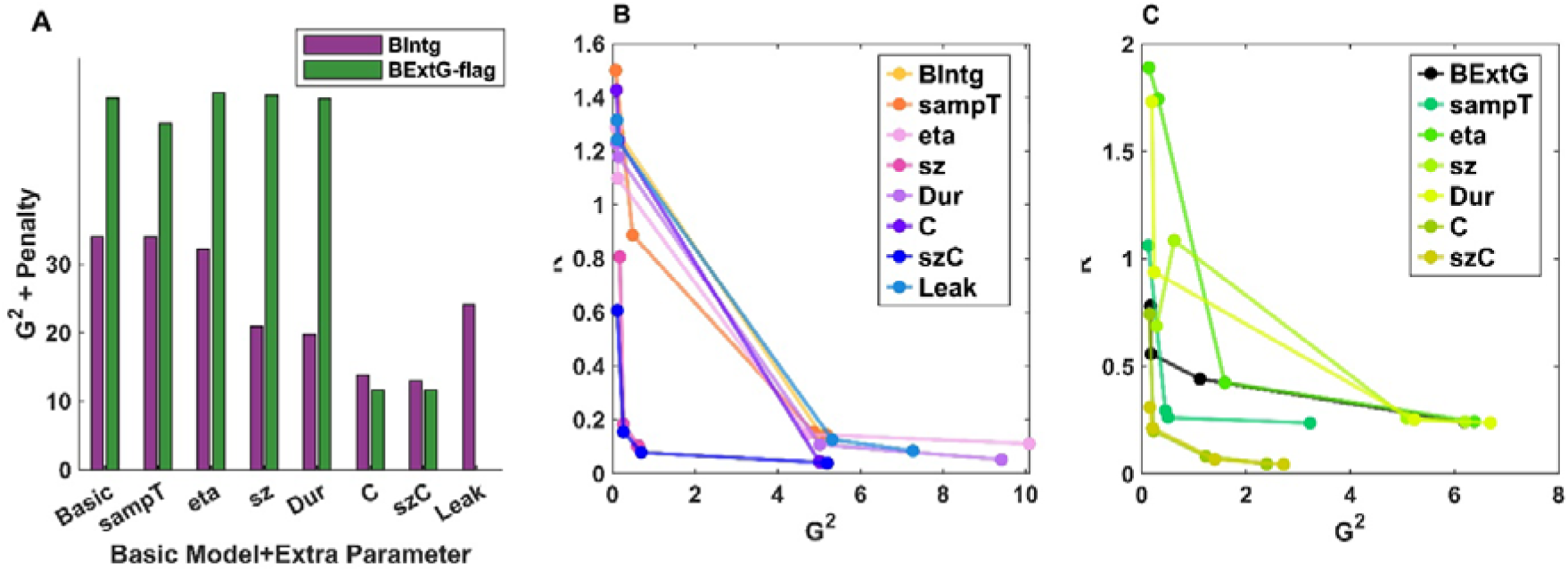
Impact of including additional, plausible free parameters to the basic neurally-informed, integration and Extremum-flagging non-integration models, taking the neural constraint weighting of 100 as a representative case. (A) Full objective function value (G^2^+neural penalty term) across models at weighting of 100. For the integration models, both starting point variability (‘sz’) and the collapsing bound improved the fit, while the competitive Extremum-flagging model (BExtG-flag) underwent the greatest fit improvement with the addition of a collapsing bound. B,C) neural-behavioural fit plots illustrating the trade-off of fitting to both behaviour and the CPP for the additional-parameter models within each model class. For all models, the lowest weighting starts in the upper left part of the plot, and the trace moves downwards as better neural fits are achieved by weighting that aspect more, at some point curving rightwards as behavioural fit is compromised in favour of neural fit. (B) The integration model including starting point variability achieved a better trade-off in fitting jointly to behaviour and the CPP waveforms than any other single additional parameter (fit points reaching closer to the origin). Additionally adding a collapsing bound further improves neural fits (lower R^2^). (C) The Extremum-flagging model achieved a comparable trade-off, with good neural and behavioural fits, in this case benefitting more from a collapsing bound than starting point variability. “sampT” denotes the basic model with extra parameter as “sampling onset time”; similarly “eta” represents between-trial drift rate variability; “sz” indicates starting point variability; “Dur” corresponds to accumulation duration; “C” refers to models with a collapsing decision bound; “szC” represents models incorporating both starting point variability and a collapsing bound; and “Leak” denotes models with leaky accumulation.

**Figure 4 - Figure Supplement 2.**
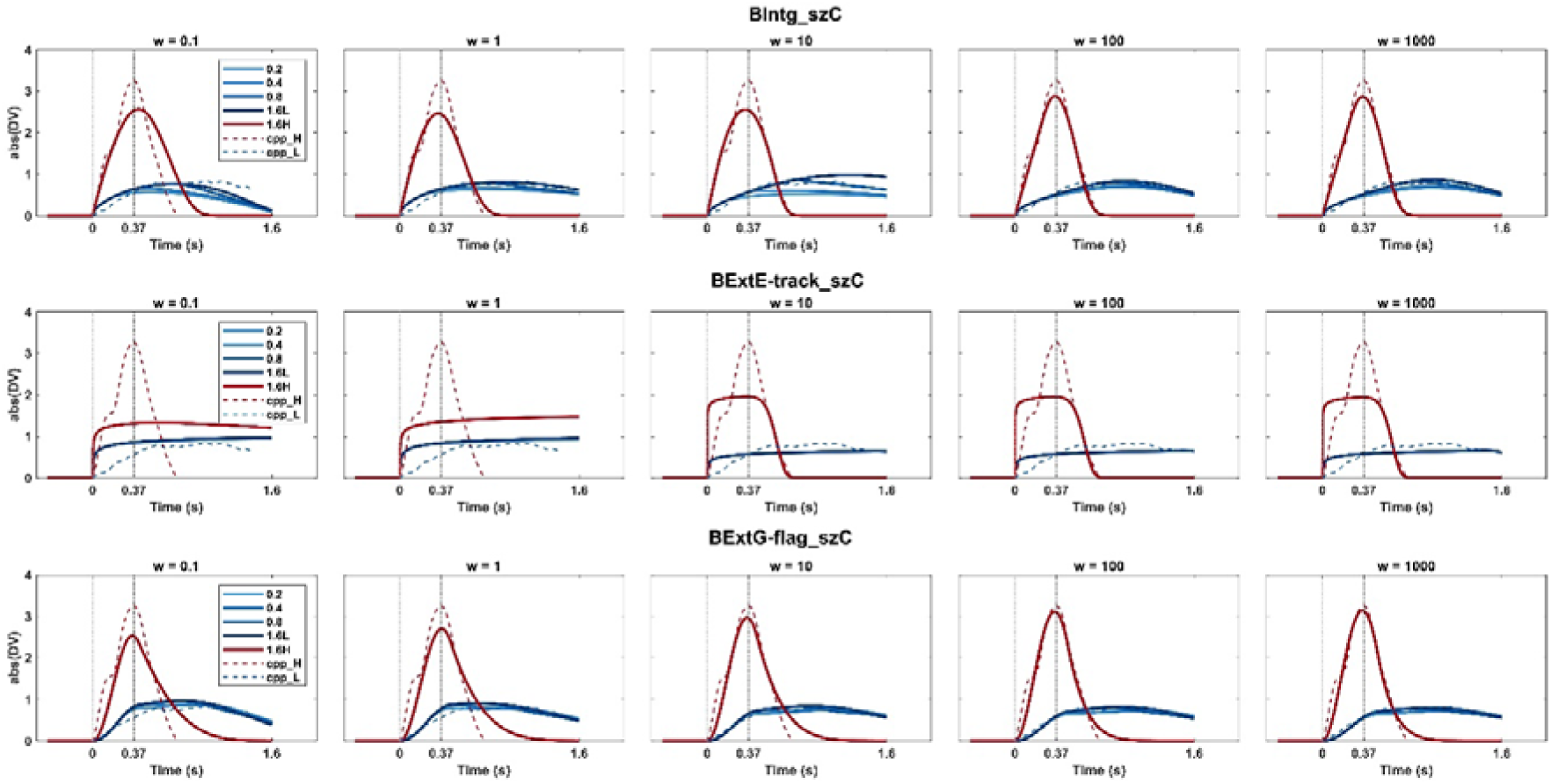
Simulated average CPP using the estimated parameters of the bounded Integration (top row), Extrema-tracking (middle), and Extremum-flagging model (bottom) for the full range of neural constraint weightings, all with starting-point variability and collapsing bounds included (same models as shown in Figure 4).

**Figure 4 - Figure Supplement 3.**
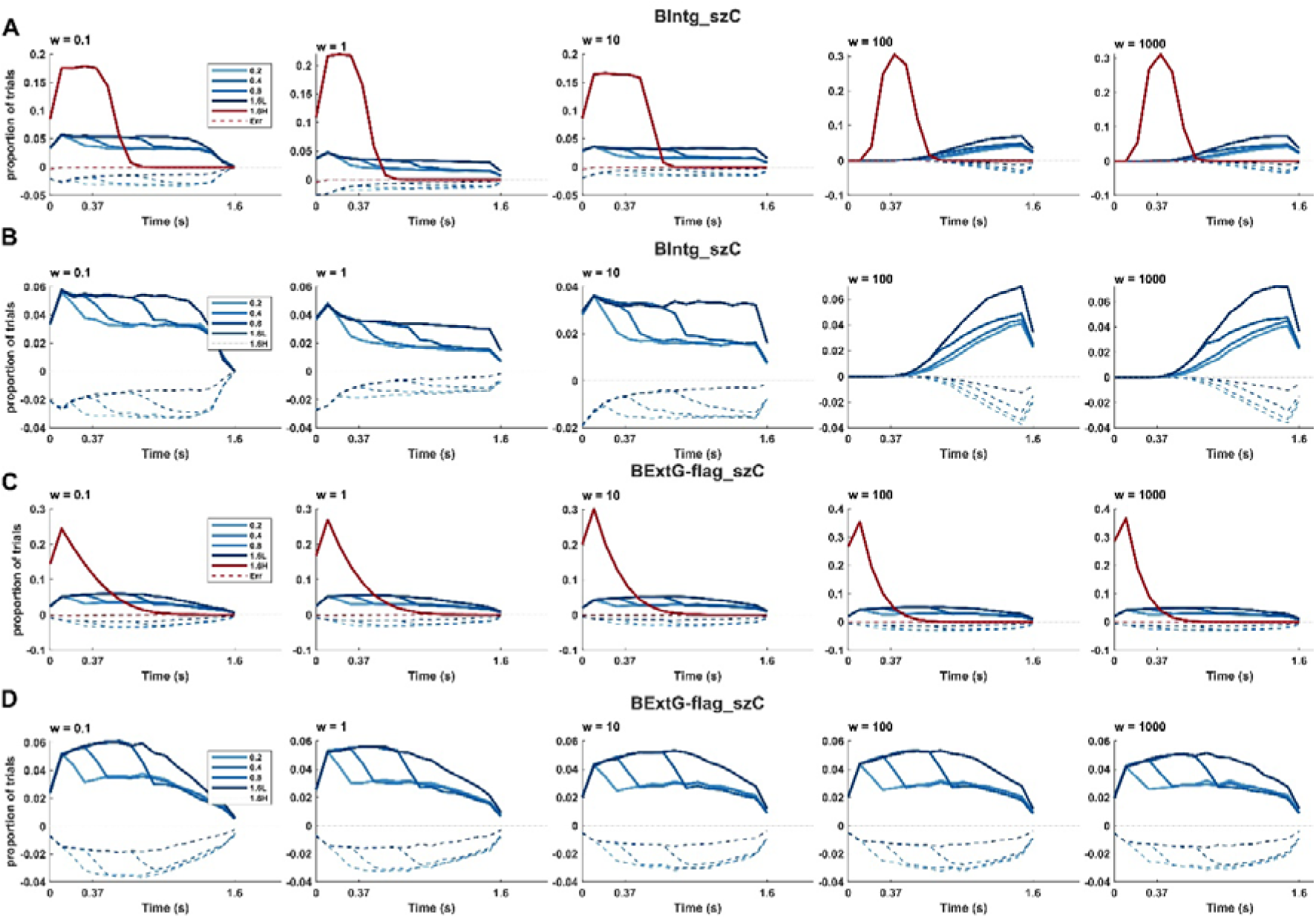
Bound crossing frequencies predicted by the Integration and Extremum-flagging models with starting-point variability and collapsing bounds. (A) Bound crossing frequencies for all easy and hard trials of the Integration model, with a zoomed-in version of just the low-contrast conditions in (B) for better visibility. Error bound crossings are shown in dashed lines against the negative y-axis. In general, the overall proportion of hard trial with bound crossings decreased markedly with increasing neural constraint weighting for the Integration model, from approximately 0.95 at w=0.1 to 0.42 at w=1000. (C & D) Bound crossings of the collapsing Extremum-flagging model. These decreased much less with increasing neural constraint weighting, from 0.92 at w=0.1 to 0.85 at w=1000.

**Figure 4 - Figure Supplement 4.**
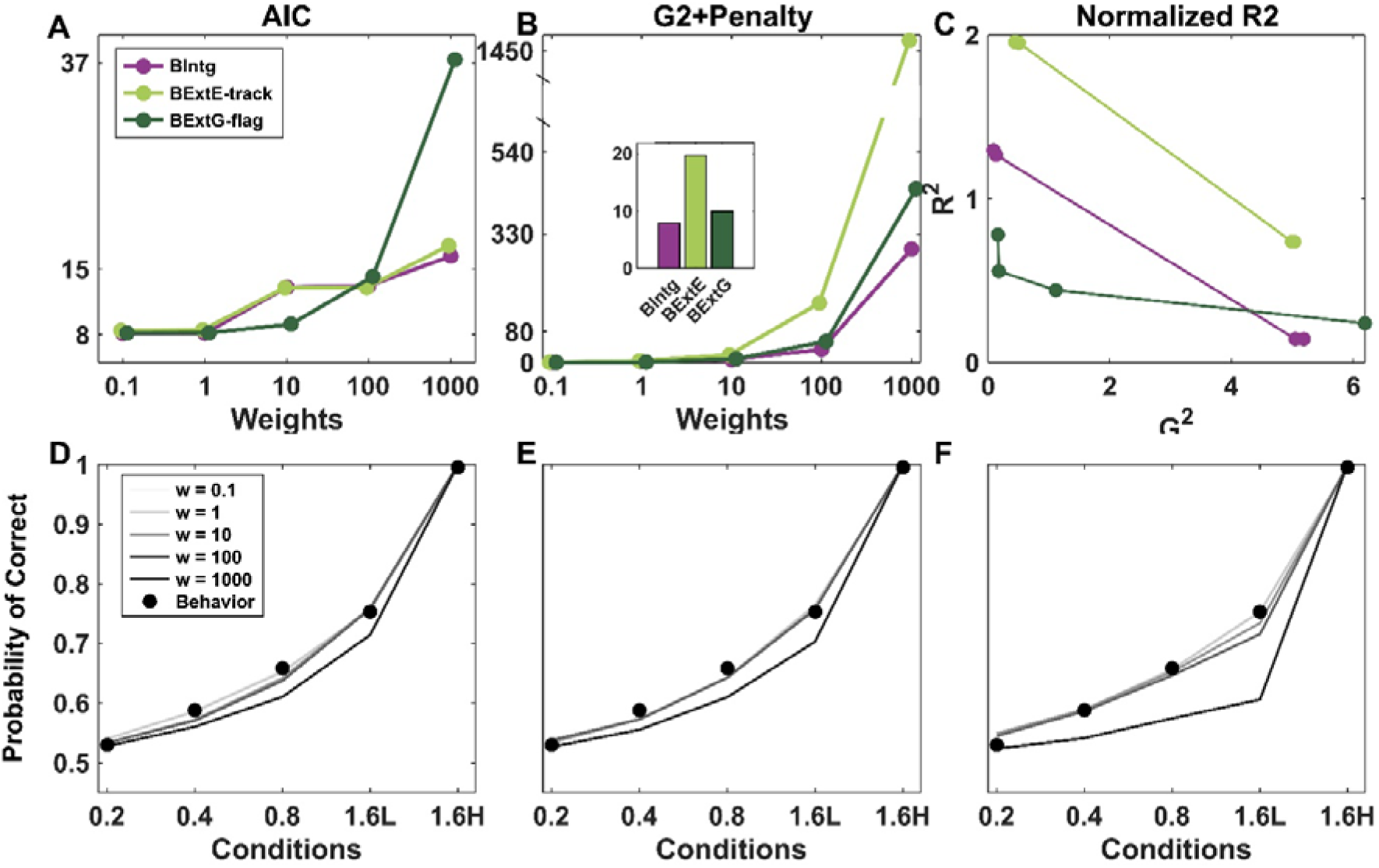
Neurally constrained for the basic models (with no additional parameters beyond two drift rates and a constant bound). (A) Akaike’s Information Criterion (AIC) derived from only the behavioural component of the neurally-constrained fit, plotted as a function of the weighting of CPP-waveform constraint. With increasing emphasis on neural data in the fit, the fit to behaviour was compromised to varying extents, and the Extremum-flagging model was best at retaining good behavioural fits at lower neural weights. (B) Overall neurally-constrained objective function (behavioural G^2^ value plus weighted neural penalty quantifying divergence in observed vs simulated CPP) as a function of neural constraint weighting. For visualisation purposes, the inset compares the objective function values to the ‘basic’ models at w = 10. The integration model outperforms at the neural weights w>1 (C) Neural fit (R^2^) plotted against behavioural fit (G^2^), illustrating a trade-off between how well the models can capture behaviour and the CPP up to w = 100. The closer the joint values come to the origin the better they are at simultaneously fitting both. (D-F) Predicted behavioural accuracy for a range of neural constraint weightings (‘w’) for (D) Integration (E) the Extremum-tracking and (F) the Extremum-flagging model.

**Figure 4 - Figure Supplement 5.**
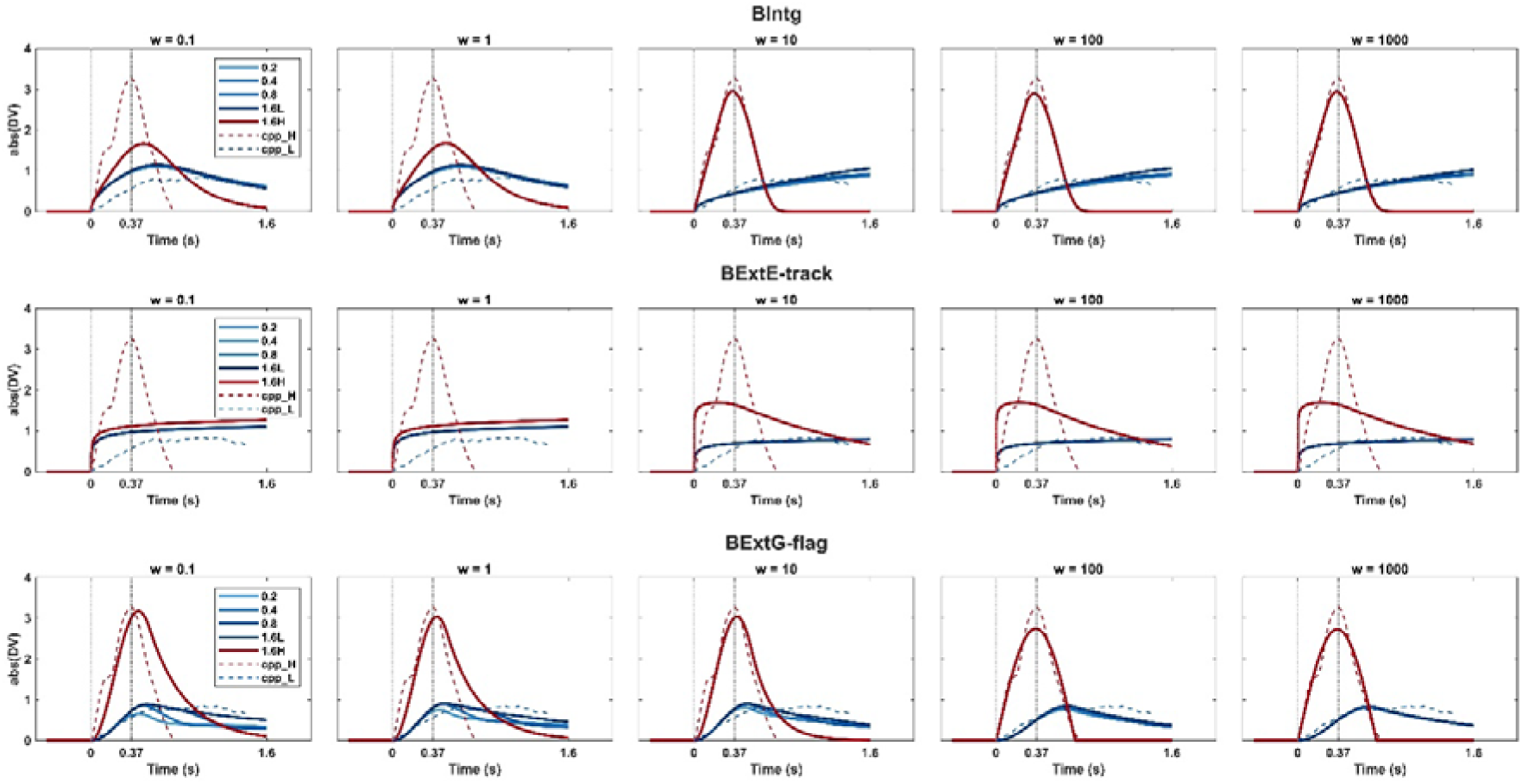
Simulated CPP signals for all basic models (with no additional parameters beyond two drift rates and a constant bound), for each of the neural constraint weightings tested. The simulated waveforms over the neural weights showed that both the integration model, **BIntg**, and Extremum-flagging model, **BExtG-flag**, are qualitatively successful in reproducing the CPP waveforms. The **BExtE-track** model sacrifices the fit to the easy condition to have a better prediction on the weak contrast conditions (The blue line fits better to the real CPP over the higher weights although the falling-down peak in the easy condition in red is not captured).

**Figure 4 - Figure Supplement 6.**
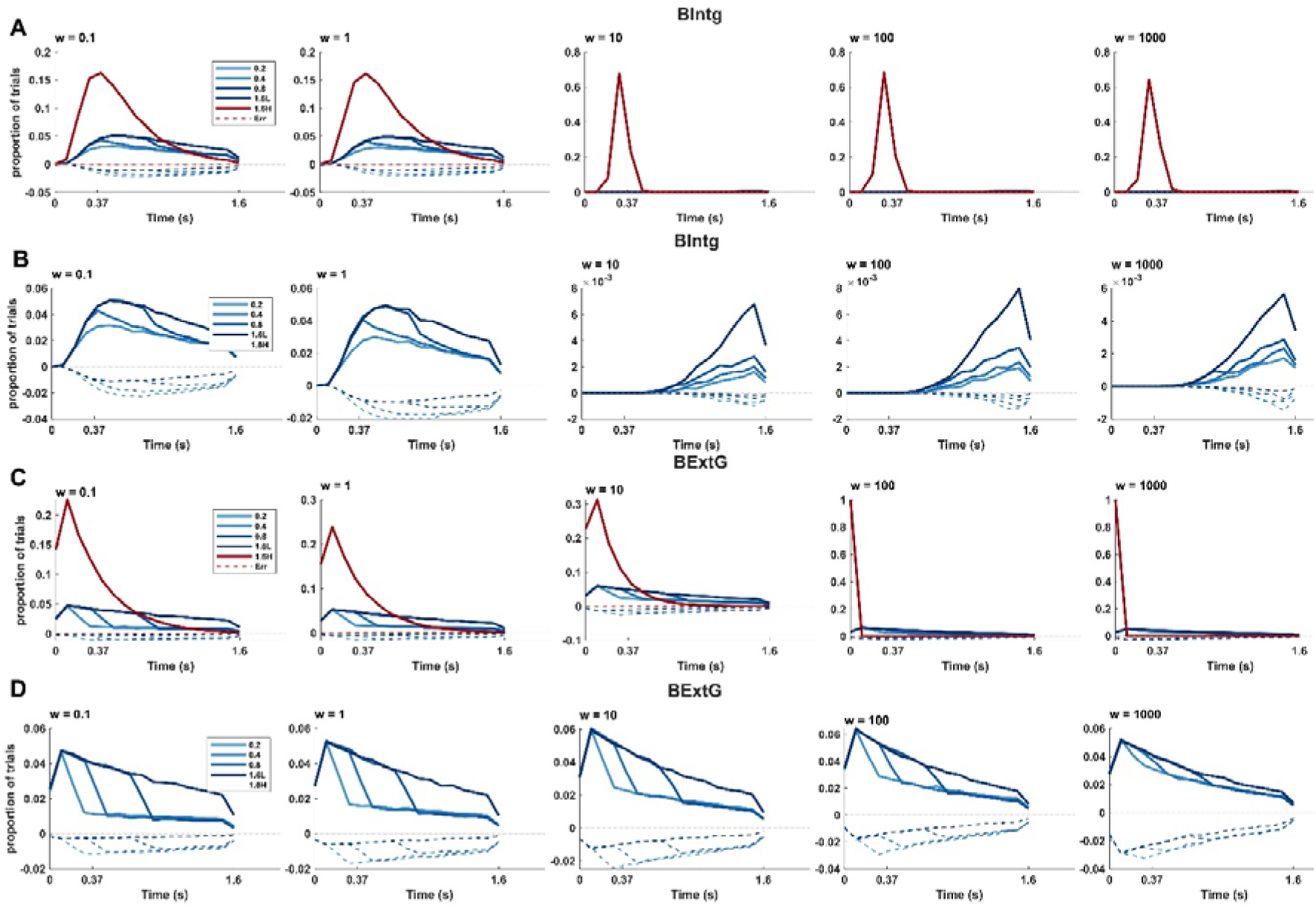
Bound crossing frequencies in correct and error trials of “basic” neurally constrained models with no additional parameters beyond two drift rates and a constant bound. (**A**) Bound crossings of all easy and hard conditions for the Integration model. The portion of the hard trials crossing the bound decrease across the neural weights. In contrast, easier trials are able to surpass the bound at higher neural constraint weightings. The drastic change occurred at w > 1. Further, the number of error trials reduces by focusing more on the neural weights. (B) The bound crossings of only the low-contrast conditions in the bounded integration, BIntg, are shown again on their own for better visibility. The bound crossings are delayed in the higher neural weights toward the later time in the trials. (**C**) Bound crossings for all conditions of the Extremum-flagging model. Similar to the Integration model, a transition happened at w>1 where reproducing the early CPP peak improved. (D) A replotting of only the low-contrast conditions of the BExtG-flag model for visibility.

**Figure 4 - Figure Supplement 7.**
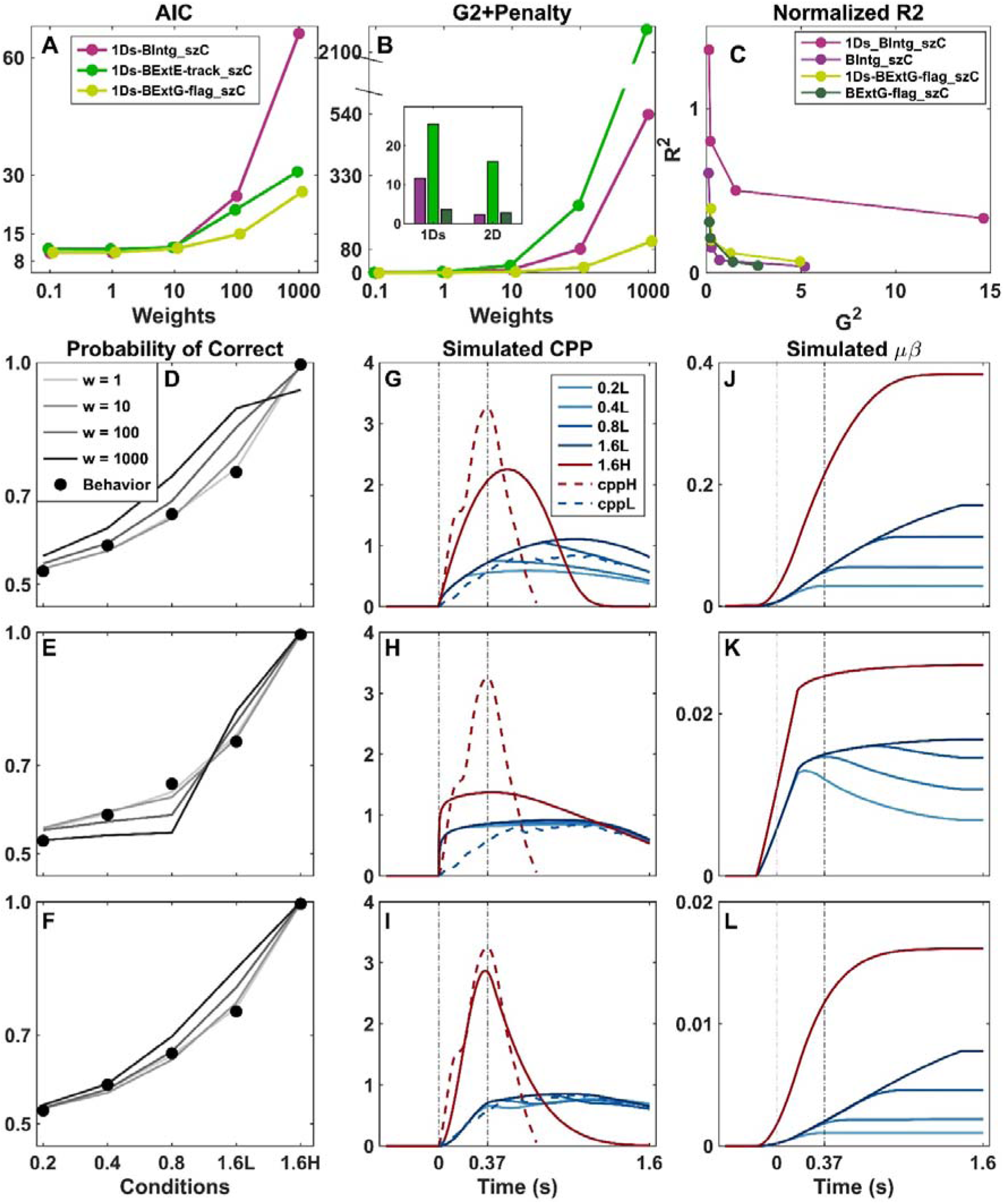
Neurally constrained models and their predictions for models with 4-fold scaled drift rate to match physical contrasts. (A-B) Sensitivity analysis showed that the Extremum-flagging models outperformed the integration and Extremum-tracking models in capturing behaviour at higher neural weights. In line with the fact that the Integration model estimates a much higher ratio of drift rates when allowed to do so, it provides a poor fit for a constrained 4-fold scaling of drift rate. The inset in panel B compares the G^2^ plus penalty of these one-drift-rate-scaled (“1Ds”) models compared to the corresponding models with two free drift rates (“2D”) at w = 10. (C) The trade-off shows that the Integration and Extremum-flagging models with two free drift rates outperform their corresponding 4-fold scaled models, but the 4-fold scaled Integration model has particular difficulty capturing the CPP dynamics. (D-F) All models compensate for accuracy at higher weights to provide better fit to the neural data. The 4-fold scaled drift rate is not enough for the Integration model to capture accuracy in the high contrast trials. (G-I) Simulated normalised average CPP for the models at w = 10. The simulated normalised average CPP (solid lines) is shown for all five conditions. The empirical data is also shown (dashed lines) with the waveforms for the four low-contrast conditions averaged together. (J-L) Simulated motor preparation. Same as (G-I) except showing the trial-averaged differential DV (without taking the absolute value) and having the signal sustain beyond bound crossing.

**Figure 4 - Figure Supplement 8.**
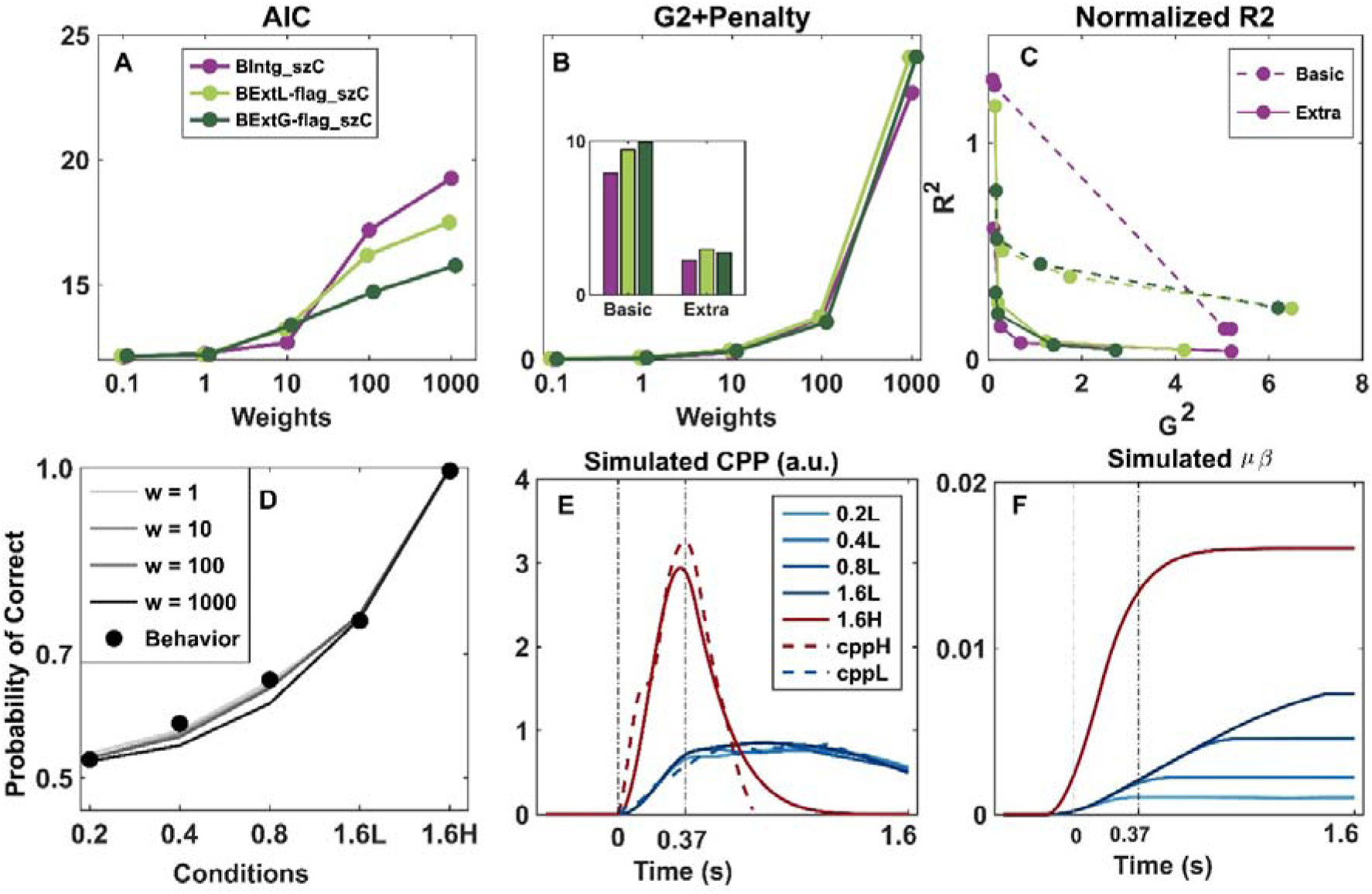
Comparison of Extremum-flagging model with a default of taking the last sample (‘BExtL’)) vs taking a random guess (‘BExtG’), when a bound is not reached during the stimulus. Apart from a tendency to sacrifice the fit to accuracy more at higher neural weightings, the model works very similarly for the two alternative default strategies. Parameter estimates were also similar at w = 10.

