## Supplementals for "Probing the role of sequential sampling and integration in decisions about protracted, noiseless stimuli"

### Supplementary Information

**Table S1** Pairwise comparisons of accuracy, showing a significant difference between each pair of consecutive durations

| **Duration 1** | **Duration 2** | **T** | **dof** | **p-value (uncorrected)** |
| --- | --- | --- | --- | --- |
| **0.2s** | 0.4s | -3.78647 | 15 | 1.79 x 10^-3^ |
| **0.4s** | 0.8s | -3.74 | 15 | 1.97 x 10^-3^ |
| **0.8s** | 1.6s | -6.19855 | 15 | 1.70 x 10^-5^ |

**Table S2** Pairwise comparisons of mean RT. There was a significant difference in reaction time between each pair of consecutive durations except for the shortest durations (0.2s vs 0.4s).

| **Duration 1** | **Duration 2** | **T** | **dof** | **p-value (uncorrected)** |
| --- | --- | --- | --- | --- |
| **0.2s** | 0.4s | 1.40869 | 15 | 1.8 x 10^-1^ |
| **0.4s** | 0.8s | 17.8368 | 15 | 1.60 x 10^-11^ |
| **0.8s** | 1.6s | 2.49377 | 15 | 2.48 x 10^-2^ |

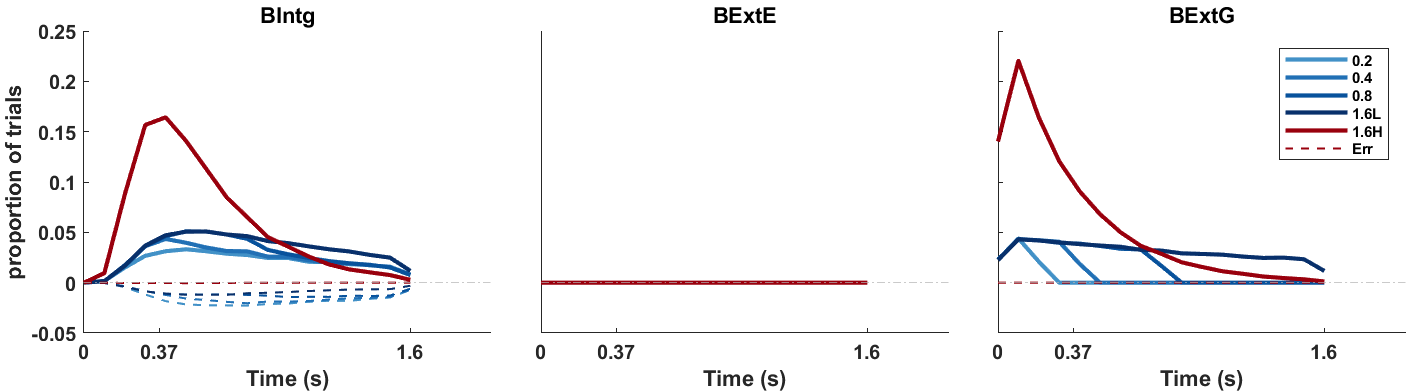

**Figure S1**. Histograms of Bound-crossing times per conditions in behavior-only models that included a bound. The integration model, **BIntg**, allows 55-60% of trials to exceed the bound in the low-contrast conditions and nearly all trials in the high-contrast conditions, with the relative proportion of correct vs error bound crossing scaling with duration. In the **BExtE** model, the bound is set so high that it is never crossed, thus functioning in the same way as the unbounded model. This is because it can only achieve the full degree of accuracy improvement with duration when the opportunity for correct revision of the running extremum derives from the full remaining non-zero evidence. Bound crossings would reduce these revision opportunities. The **BExtG** model produces a good fit by setting a relatively high bound and assuming a drift rate for low-contrast conditions that is almost as high as the high-contrast condition. This produces no error bound crossings and an almost uniform distribution of correct bound crossing times during the evidence periods for hard trials, which produces a linearly increasing ratio of correct bound crossings to guesses across durations, in turn yielding a linear increase in accuracies.

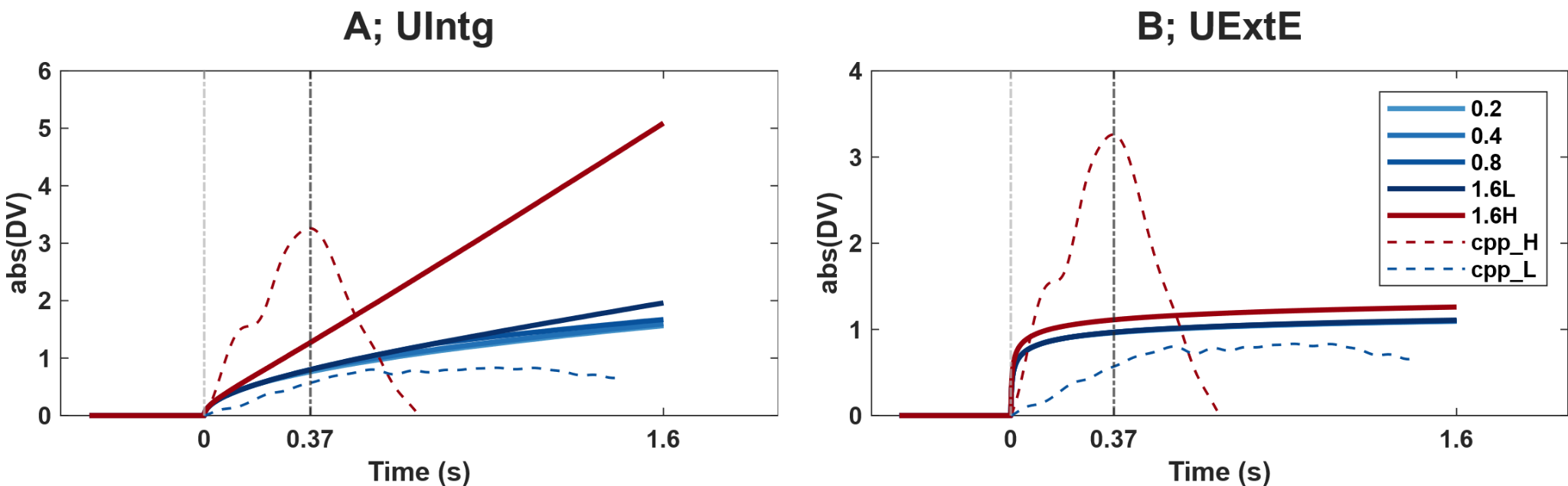

**Figure S2** Simulated decision variable waveforms of the unbounded integration (A; UIntg) and non-integration (B; UExtE) models that produced good fits to behaviour alone, illustrating that neither of these models can reproduce the peak and falling-down of the empirical CPP signal (dashed) in the high-contrast condition. For consistency with the neurally-constrained modelling analysis, empirical traces show the mean CPP segments that are used as constraints, with low-contrast conditions averaged (‘cppL;’ See Neurally-constrained model-fitting in Results), whereas all five conditions are simulated for the models.

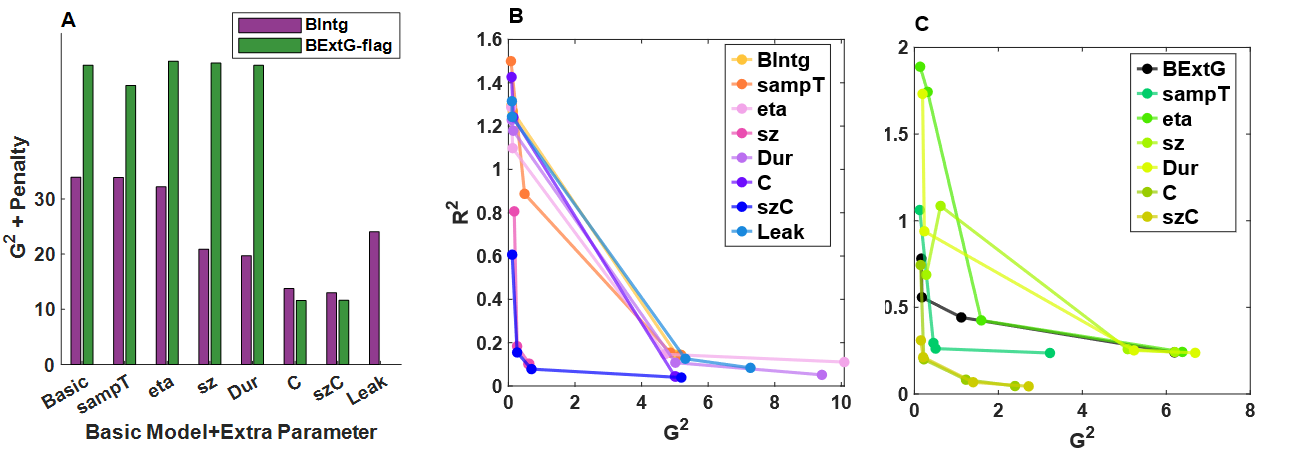

**Figure S3** Impact of including additional, plausible free parameters to the basic neurally-informed, integration and Extremum-flagging non-integration models, taking the neural constraint weighting of 100 as a representative case. A) full objective function value (G^2^+neural penalty term) across models at weighting of 100. For the integration models, both starting point variability (‘sz’) and the collapsing bound improved the fit, while the competitive Extremum-flagging model (BExtG-flag) underwent the greatest fit improvement with the addition of a collapsing bound. B,C) neural-behavioural fit plots illustrating the trade-off of fitting to both behavior and the CPP for the additional-parameter models within each model class. For all models, the lowest weighting starts in the upper left part of the plot, and the trace moves downwards as better neural fits are achieved by weighting that aspect more, at some point curving rightwards as behavioural fit is compromised in favour of neural fit. B) The integration model including starting point variability achieved a better trade-off in fitting jointly to behaviour and the CPP waveforms than any other single additional parameter (fit points reaching closer to origin). Additionally adding a collapsing bound further improves neural fits (lower R^2^). C) The Extremum-flagging model achieved a comparable trade-off, with good neural and behavioural fits, in this case benefitting more from a collapsing bound than starting point variability. “sampT” denotes the basic model with extra parameter as “sampling onset time”; similarly “eta” represents between-trial drift rate variability; “sz” indicates starting point variability; “Dur” corresponds to accumulation duration; “C” refers to models with a collapsing decision bound; “szC” represents models incorporating both starting point variability and a collapsing bound; and “Leak” denotes models with leaky accumulation.

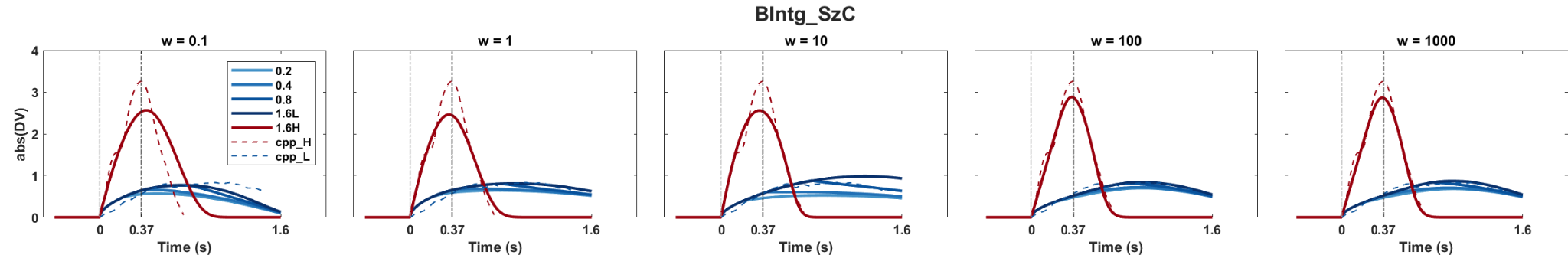

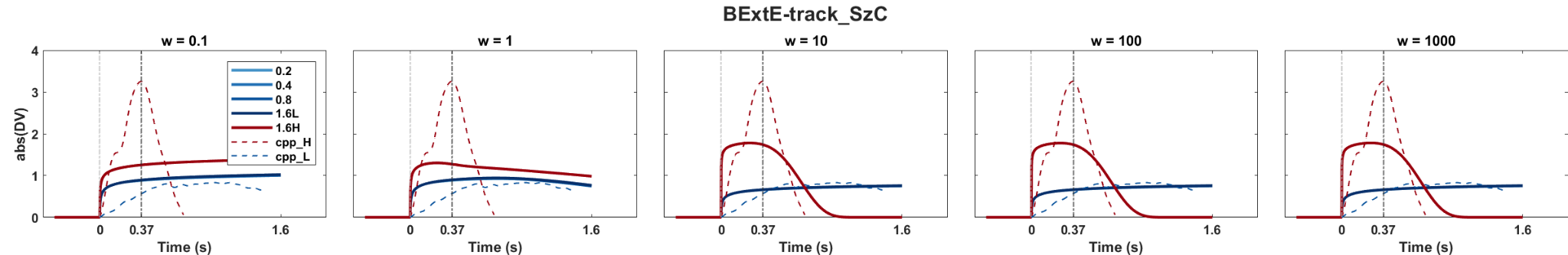

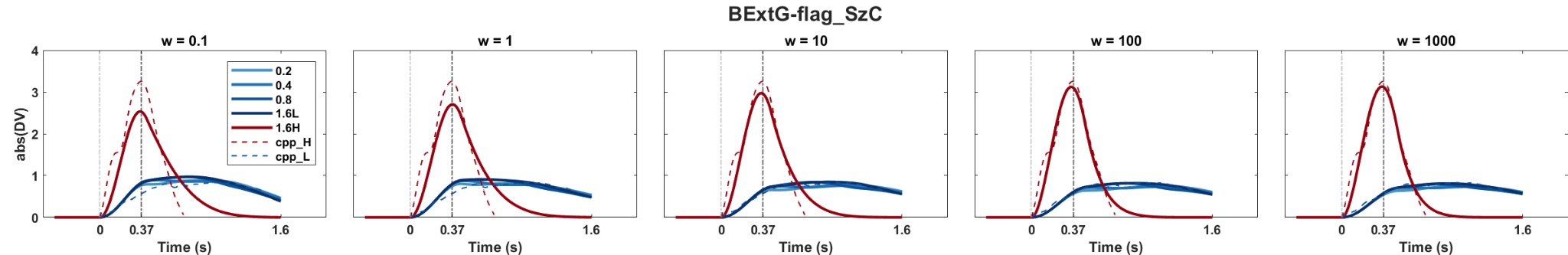

**Figure S4** Simulated average CPP using the estimated parameters of the bounded Integration (top row), extrema-tracking (middle), and Extremum-flagging model (bottom) for the full range of neural constraint weightings, all with starting-point variability and collapsing bounds included (same models as shown in Figure 4).

**Table S3** Neurally constrained Integration, Extremum-tracking and Extremum-flagging models with various extra model features at neural constraint weighting **w = 10**. ‘BIntg’ refers to the bounded Integration model, ‘BExtE’ refers to Extremum-tracking, and ‘BExtG’ refers to Extremum-flagging which has a default guess.  ‘FlagWidth’ refers to the width of half-sine while t05 and K are semi-saturation constant and collapsing rate parameters of the collapsing bound function.

| **w = 10** | **G^2^+Penalty** | **B** | **D1** | **D2** | **FlagWidth** | **extraParam** | **t05, K** |
| --- | --- | --- | --- | --- | --- | --- | --- |
| **BIntg** | 7.9864 | 0.3591 | 0.0566 | 1.0991 | 0.6688 | - | - |
| **BIntg_eta** | 7.6351 | 0.3631 | 0.0641 | 1.1498 | 0.7281 | eta = 0.0513 | - |
| **BIntg_sz** | 2.6871 | 0.2988 | 0.1056 | 1.3391 | 0.7507 | sz = 0.5987 | - |
| **BIntg_Dur** | 7.1693 | 0.3374 | 0.0593 | 1.0801 | 0.7441 | Dur = 1.1515 | - |
| **BIntg_sampT** | 7.9435 | 0.3724 | 0.0593 | 1.0998 | 0.6338 | accT =-0.0873 | - |
| **BIntg_C** | 5.8835 | 0.3578 | 0.0578 | 0.9601 | 0.7161 | - | K =0.4594 |
| **BIntg_szC** | 2.2533 | 0.4683 | 0.1607 | 1.4049 | 0.2502 | sz = 0.9419 | K = 0.312 |
| **BIntg_szC (NL)** | 2.5622 | 0.2996 | 0.1133 | 1.2131 | 0.6619 | sz = 0.6 | t0.5 =3.9996, K =0.8766 |
| **BIntg_Leak** | 7.8308 | 0.3159 | 0.0577 | 1.0194 | 0.7582 | Leak=0.0002 | - |
| **BExtE** | 19.7663 | 0.035 | 0.3815 | 10.865 | 0.9 | - | - |
| **BExtE_szC** | 17.7528 | 0.04 | 0.3813 | 12.3253 | 0.9 | sz = 0.0004 | K = 0.3205 |
| **BExtE_szC(NL)** | 17.869 | 0.04 | 0.3808 | 12.3183 | 0.89 | sz =0.00002 | t0.5 =2.4969, K =0.9101 |
| **BExtG** | 9.9581 | 0.0145 | 0.5079 | 2.0663 | 0.52 | - | - |
| **BExtG_eta** | 10.093 | 0.0143 | 0.4666 | 2.2095 | 0.5705 | eta = 0.0829 | - |
| **BExtG_sz** | 10.2583 | 0.0142 | 0.4742 | 10.173 | 0.7305 | sz =0.0004 | - |
| **BExtG_Dur** | 10.2672 | 0.0142 | 0.4564 | 15.3698 | 0.7231 | Dur = 1.5716 | - |
| **BExtG_sampT** | 5.7477 | 0.0143 | 0.4909 | 15.7325 | 0.7215 | sampT =0.0119 | - |
| **BExtG_C** | 2.877 | 0.015 | 0.4609 | 2.1092 | 0.4379 | - | K = 0.0889 |
| **BExtG_szC** | 2.7589 | 0.015 | 0.4387 | 2.1614 | 0.4462 | sz = | K = 0.1062 |
| **BExtG_szC(NL)** | 2.9558 | 0.015 | 0.4471 | 2.1584 | 0.4507 | sz =0.0005 | t05 = 2.1641; K=0.3177 |

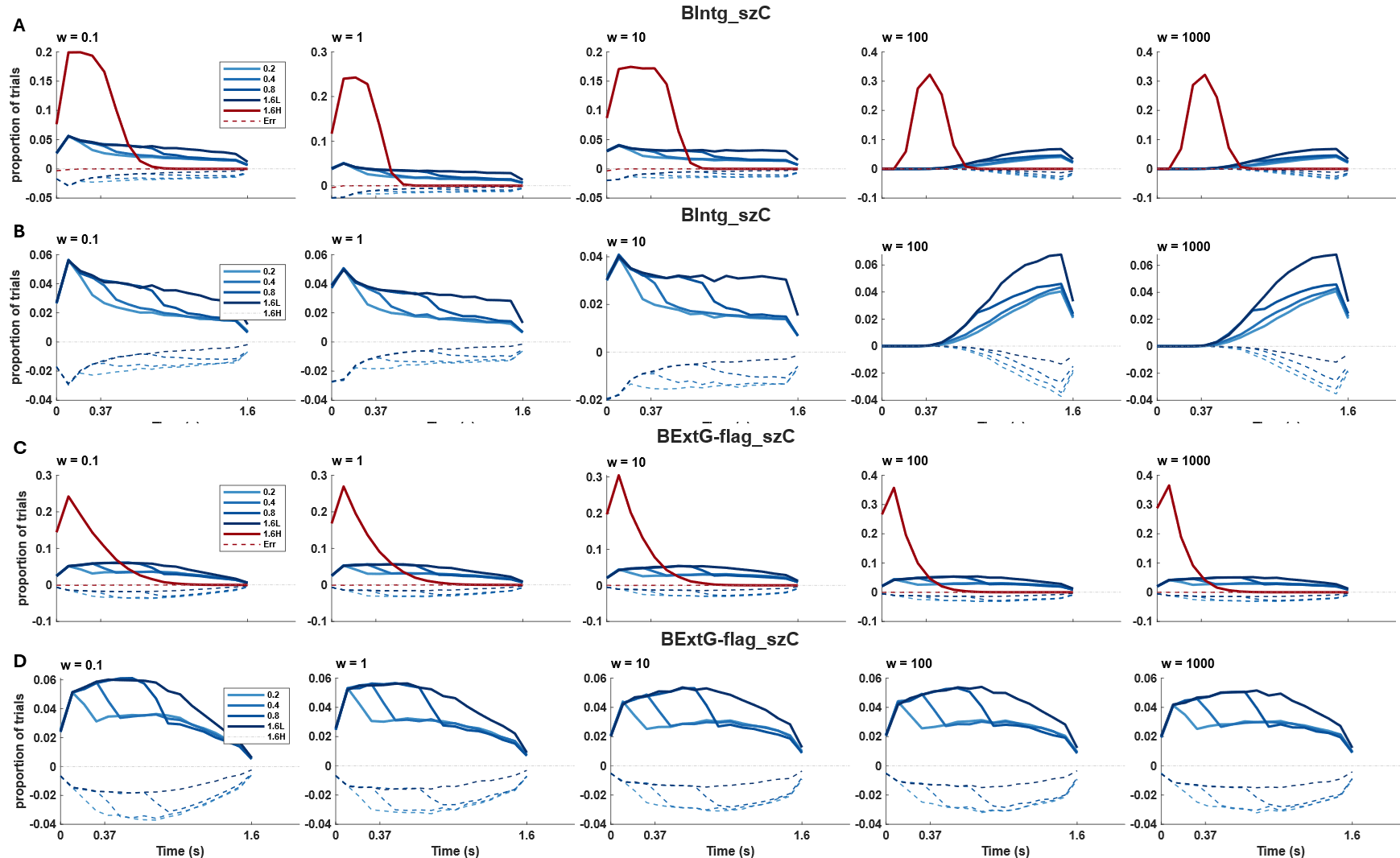

**Figure S5.** Bound crossing frequencies predicted by the Integration and Extremum-flagging models with starting-point variability and collapsing bounds. (A) bound crossing frequencies for all easy and hard trials of the Integration model, with a zoomed-in version of just the low-contrast conditions in (B) for better visibility. Error bound crossings are shown in dashed lines against the negative y-axis. In general, the overall proportion of hard trial with bound crossings decreased markedly with increasing neural constraint weighting for the Integration model, from approximately 0.95 at w=0.1 to 0.42 at w=1000. (C & D) bound crossings of the collapsing Extremum-flagging model. These decreased much less with increasing neural constraint weighting, from 0.92 at w=0.1 to 0.85 at w=1000.

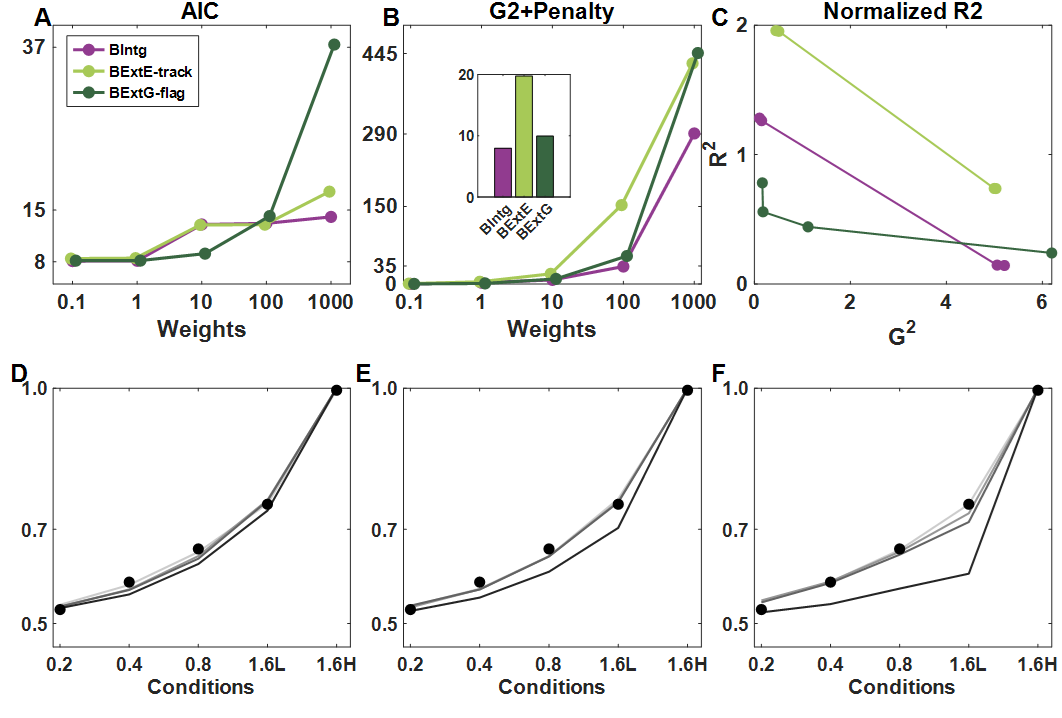

**Figure S6**  Neurally constrained for the basic models (with no additional parameters beyond two drift rates and a constant bound). (**A**) Akaike’s Information Criterion (AIC) derived from only the behavioural component of the neurally-constrained fit, plotted as a function of the weighting of CPP-waveform constraint. With increasing emphasis on neural data in the fit, the fit to behavior was compromised to varying extents, and the Extremum-flagging model was best at retaining good behavioral fits at lower neural weights. (**B**) Overall neurally-constrained objective function (behavioral G^2^ value plus weighted neural penalty quantifying divergence in observed vs simulated CPP) as a function of neural constraint weighting. For the visualisation purposes, the inset compares the objective function values to the ‘basic’ models at w = 10. The integration model outperforms at the neural weights w>1 (**C**) Neural fit (R^2^) plotted against behavioral fit (G^2^), illustrating a trade-off between how well the models can capture behavior and the CPP up to w = 100. The closer the joint values come to the origin the better they are at simultaneously fitting both. (**D-F**) Predicted behavioral accuracy for a range of neural constraint weightings (‘w’) for (D) Integration (E) the Extremum-tracking and (F) the Extremum-flagging model.

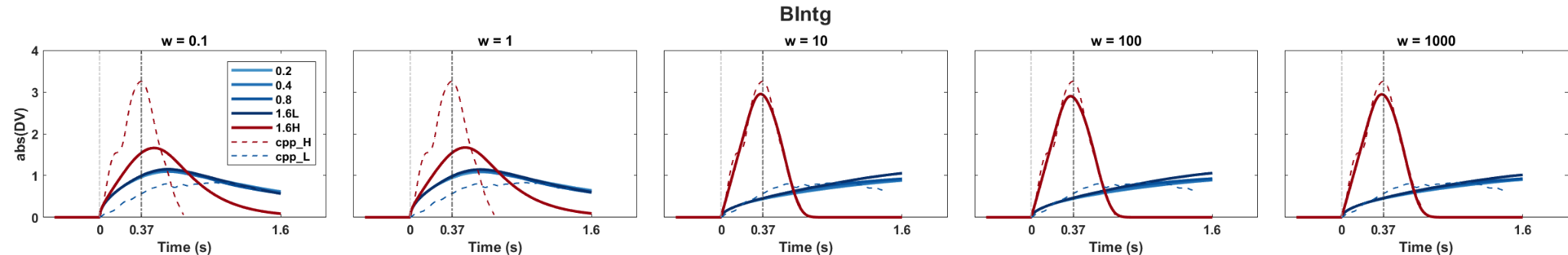

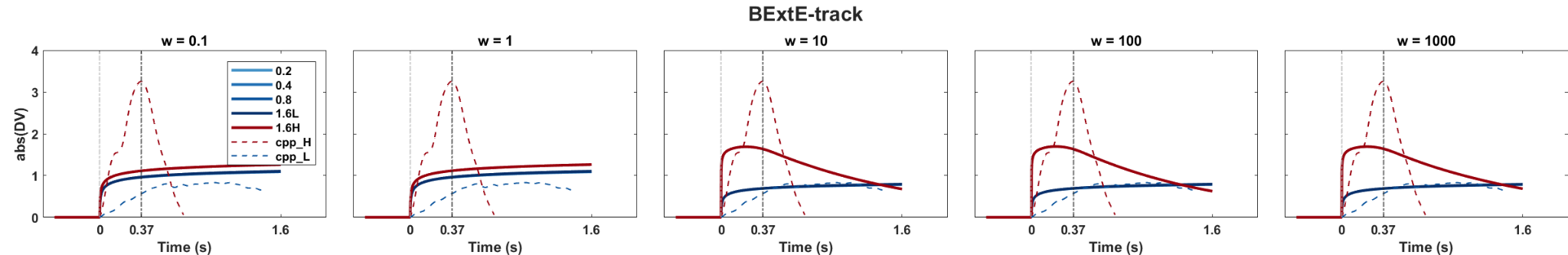

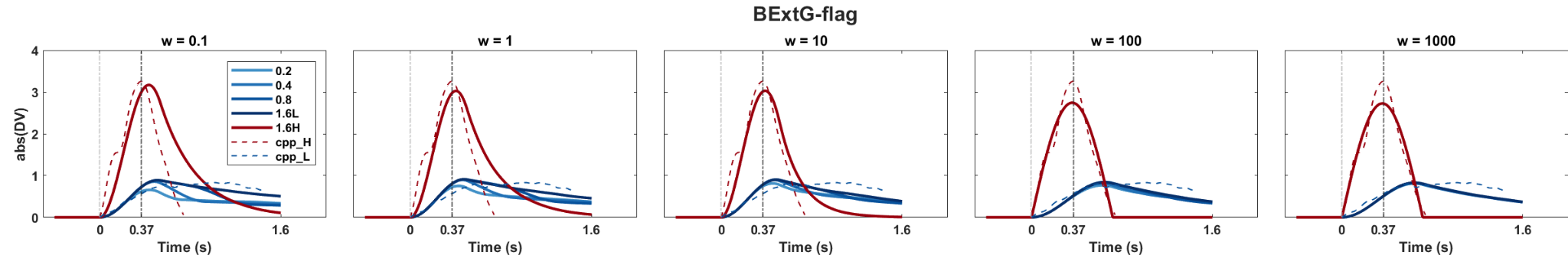

**Figure S7** Simulated CPP signals for all basic models (with no additional parameters beyond two drift rates and a constant bound), for each of the neural constraint weightings tested. The simulated waveforms over the neural weights showed that both the integration model, **BIntg**, and Extremum-flagging model, **BExtG-flag**, qualitatively are successful in reproducing the CPP waveforms. The **BExtE-track** model sacrifices the fit to the easy condition to have a better prediction on the weak contrast conditions (The blue line fits better to the real CPP over the higher weights although the falling-down peak in the easy condition in red is not captured).

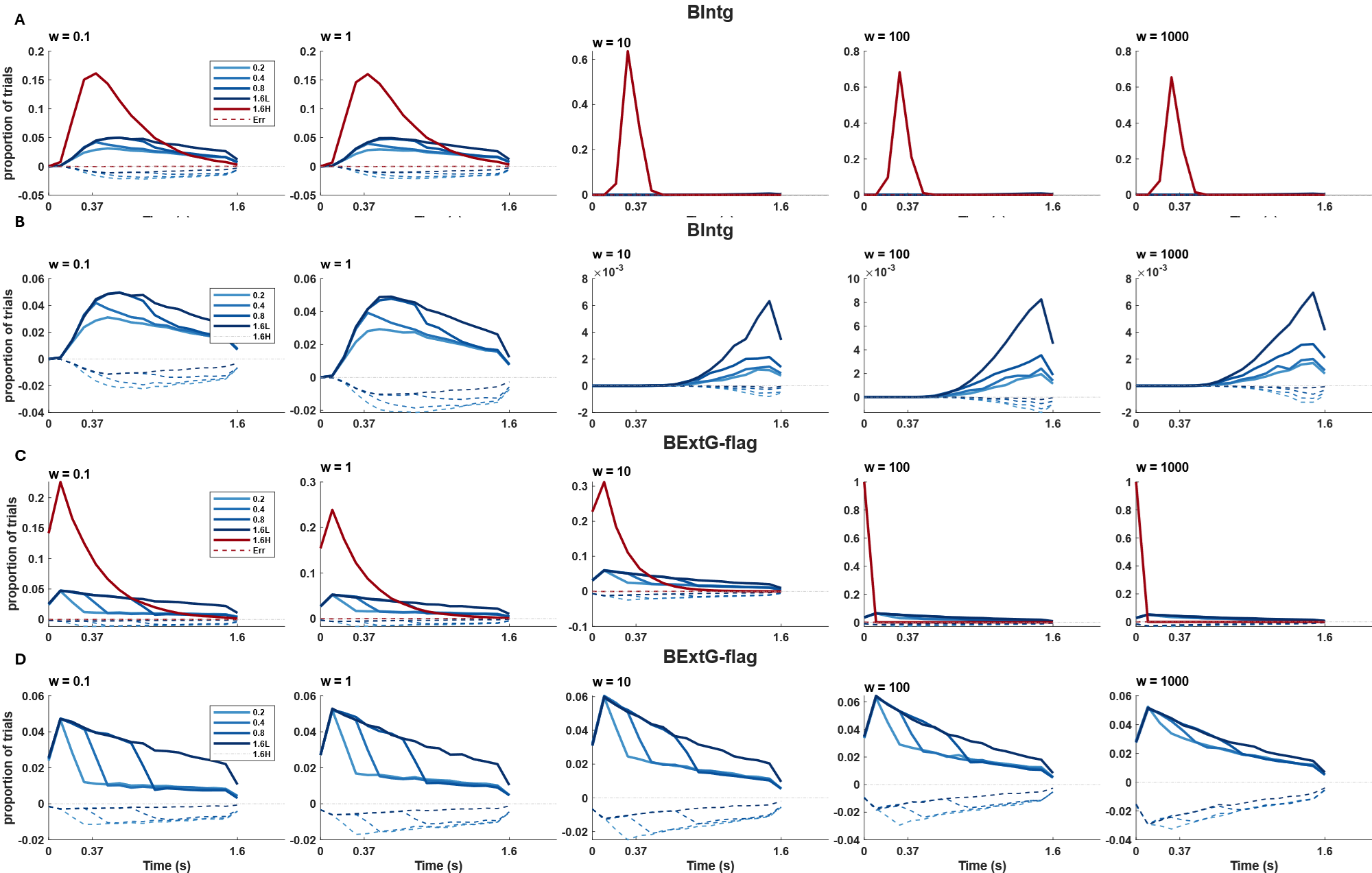

**Figure S8** Bound crossing frequencies in correct and error trials of “basic” neurally constrained models with no additional parameters beyond two drift rates and a constant bound. (**A**) bound crossings of all easy and hard conditions for the Integration model. The portion of the hard trials crossing the bound decrease across the neural weights. In contrast, more easy trials are able to surpass the bound at higher neural constraint weightings. The drastic change occurred at w > 1. Further, the number of error trials reduces by focusing more on the neural weights. (**B**) The bound crossings of only the low-contrast conditions in the bounded integration, Intg, are shown again on their own for better visibility. The bound crossings are delayed in the higher neural weights toward the later time in the trials. (**C**) Bound crossings for all conditions of the Extremum-flagging model. Similar to the Integration model, a transition happened at w>1 where reproducing the early CPP peak improved. (**D**) A replotting of only the low-contrast conditions of the ExtG-flag model for visibility.

**Table S4**. Allowed parameter ranges for model fits, i.e. the minimum and maximum values that each parameter can take in the SIMPLEX (fminsearch) algorithm. ‘Default value’ indicates parameter settings used when the parameter is fixed rather than free. An asterisk * marks the parameters with 5 grid points in the models with both starting point variability and a collapsing bound.

| **Parameters** |  | **Integration** | **ExtE-tracking** | **ExtG-flagging** | **Inactive Default value** |
| --- | --- | --- | --- | --- | --- |
| **bound** | B | [0, 0.65] | [0, 0.04] | [0, 0.04] | - |
| **Drift rate (Low)** | D1 | [0, 0.6] | [0, 2] | [0, 7] | - |
| **Drift rate (High)** | D2 | [0, 2] | [0, 15] | [0, 15] | - |
| **Between trial Variability** | eta | [0, 0.5] | [0, 0.5] | [0, 0.5] | 0 |
| **Starting point*** | sz | [0, 1] | [0, 0.04] | [0, 0.04] | 0 |
| **Duration** | Dur | [0.2, 1.6] | [0.2, 1.6] | [0.2, 1.6] | 1.6 |
| **Accumulation onset Time** | sampT | [-0.4, 0.4] | [-0.4, 0.4] | [-0.4, 0.4] | 0 |
| **collapse*** | k | [-1 1] | [-1, 1] | [-1 1] | 0 |
| **Leak** | leak | [0, 1] | - | - | 0 |
| **FlagWidth** | wd | - | - | [0.002, 0.9] | - |
| **Semi-satuartion time** | t05 | [0.1 4] | [0.1 2.5] | [0.1 2.5] | - |

**Table S5** Neurally constrained Integration, Extremum-tracking and Extremum-flagging models with one extra feature at neural constraint weighting **w = 0.1**. ‘BIntg’ refers to the bounded Integration model, ‘BExtE’ refers to Extremum-tracking, and ‘BExtG’ refers to Extremum-flagging which has a guess default. ‘FlagWidth’ refers to the width of half-sine. t05 and K are semi-saturation constant and collapsing rate parameters of the nonlinear collapsing bound function, whereas the linear collapsing bound function only has a slope parameter K.

| **w = 0.1** | **G^2^+Penalty** | **B** | **D1** | **D2** | **FlagWidth** | **extraParam** | **t05, K** |
| --- | --- | --- | --- | --- | --- | --- | --- |
| **BIntg** | 0.3614 | 0.1318 | 0.0566 | 0.2228 | 0.857 | - | - |
| **BIntg_eta** | 0.3404 | 0.2053 | 0.0674 | 0.2642 | 0.8895 | eta = 0.0609 | - |
| **BIntg_sz** | 0.3352 | 0.1443 | 0.0668 | 0.3486 | 0.8923 | sz =0.2108 | - |
| **BIntg_Dur** | 0.3441 | 0.1725 | 0.0591 | 0.2304 | 0.4564 | Dur = 1.3034 | - |
| **BIntg_sampT** | 0.3755 | 0.2453 | 0.0634 | 0.2423 | 0.5171 | sampT=-0.2244 | - |
| **BIntg_C** | 0.3793 | 0.422 | 0.058 | 0.2233 | 0.8931 | - | K = 0.6866 |
| **BIntg_szC** | 0.2328 | 0.3137 | 0.1288 | 0.8927 | 0.8472 | sz = 0.6177 | K = 0.6355 |
| **BIntg_szC (NL)** | 0.1662 | 0.2888 | 0.1221 | 1.3666 | 0.8777 | sz = 0.5784 | t05 = 2.438; K = 2.0444 |
| **BIntg_Leak** | 0.3666 | 0.1291 | 0.0561 | 0.222 | 0.8295 | leak = 0.0 | - |
| **BExtE** | 0.8374 | 0.0347 | 0.3883 | 1.6309 | 0.0044 | - | - |
| **BExtE_szC** | 0.5344 | 0.04 | 0.5283 | 3.41 | 0.8986 | sz = 0.04 | K = -0.9983 |
| **BExtE_szC (NL)** | 0.7045 | 0.0257 | 0.408 | 1.9322 | 0.8911 | sz = 0.0315 | t05=2.156 ;K = -0.6199 |
| **BExtG** | 0.3203 | 0.0155 | 0.8372 | 2.1473 | 0.5367 | - | - |
| **BExtG_eta** | 0.5129 | 0.0195 | 2.7719 | 4.1325 | 0.6719 | eta =0.0996 | - |
| **BExtG_sz** | 0.4193 | 0.015 | 0.6899 | 2.3364 | 0.7015 | sz = 0.0029 | - |
| **BExtG_Dur** | 0.5397 | 0.0195 | 2.8073 | 4.1856 | 0.5624 | Dur = 1.5007 | - |
| **BExtG_sampT** | 0.3384 | 0.016 | 1.0522 | 4.6706 | 0.7095 | sampT = -0.0431 | - |
| **BExtG_C** | 0.2967 | 0.0145 | 0.4045 | 1.8311 | 0.672 | - | K = 0.1044 |
| **BExtG_szC** | 0.2158 | 0.0147 | 0.4033 | 1.7206 | 0.414 | sz = 0.0001 | K = 0.1274 |
| **BExtG_szC (NL)** | 0.2603 | 0.0151 | 0.5023 | 2.0497 | 0.4981 | sz = 0.0029 | t05 = 2.3537; K = 0.5122 |

**Table S6** Neurally constrained Integration, Extremum-tracking and Extremum-flagging models with one extra feature at neural constraint weighting **w = 1**. ‘BIntg’ refers to the bounded Integration model, ‘BExtE’ refers to Extremum-tracking, and ‘BExtG’ refers to Extremum-flagging which has a default guess. ‘FlagWidth’ refers to the width of half-sine while t05 and K are semi-saturation constant and collapsing rate parameters of the collapsing bound function.

| **w = 1** | **G^2^+Penalty** | **B** | **D1** | **D2** | **FlagWidth** | **ExtraParam** | **t05, K** |
| --- | --- | --- | --- | --- | --- | --- | --- |
| **BIntg** | 2.6731 | 0.1337 | 0.0559 | 0.2232 | 0.8663 | - | - |
| **BIntg_eta** | 2.3237 | 0.1783 | 0.069 | 0.2741 | 0.6237 | eta = 0.0635 | - |
| **BIntg_sz** | 0.6277 | 0.1893 | 0.085 | 0.9504 | 0.8973 | sz = 0.3719 | - |
| **BIntg_Dur** | 2.5003 | 0.1621 | 0.059 | 0.2358 | 0.5866 | Dur = 1.2735 | - |
| **BIntg_sampT** | 2.2597 | 0.1901 | 0.0686 | 0.6587 | 0.8998 | sampT= -0.3973 | - |
| **BIntg_C** | 2.6294 | 0.1342 | 0.0564 | 0.2284 | 0.8679 | - | K = 0.009 |
| **BIntg_szC** | 0.5706 | 0.2293 | 0.0939 | 0.9363 | 0.751 | sz = 0.4533 | K = 0.2339 |
| **BIntg_szC(NL)** | 0.598 | 0.2869 | 0.1093 | 1.1275 | 0.6546 | sz = 0.5732 | t05= 3.7864 ; K =1.4184 |
| **BIntg_Leak** | 2.592 | 0.1367 | 0.0563 | 0.2234 | 0.851 | Leak = 0.0001 | - |
| **BExtE** | 4.4127 | 0.0344 | 0.3874 | 1.6424 | 0.1795 | - | - |
| **BExtE_szC** | 3.2056 | 0.0356 | 0.5956 | 4.448 | 0.7819 | sz = 0.0393 | K = -0.2023 |
| **BExtE_szC (NL)** | 2.0551 | 0.0395 | 0.5155 | 11.0308 | 0.8852 | sz = 0.0397 | t05 = 1.8842; K = -1.8603 |
| **BExtG** | 1.2939 | 0.015 | 0.6599 | 1.959 | 0.5004 | - | - |
| **BExtG_eta** | 3.8019 | 0.0195 | 2.7484 | 4.5935 | 0.5871 | eta = 0.4978 | - |
| **BExtG_sz** | 2.7936 | 0.0171 | 1.5133 | 4.5796 | 0.8999 | sz = 0.0093 | - |
| **BExtG_Dur** | 2.1145 | 0.0159 | 1.037 | 2.37 | 0.5268 | Dur = 1.4749 | - |
| **BExtG_sampT** | 1.041 | 0.0144 | 0.531 | 12.6837 | 0.7459 | sampT =-0.0118 | - |
| **BExtG_C** | 0.6235 | 0.0147 | 0.4147 | 1.8688 | 0.4494 | - | K = 0.1042 |
| **BExtG_szC** | 0.6313 | 0.0146 | 0.4185 | 1.8496 | 0.4559 | sz = 0.00014 | K = 0.0983 |
| **BExtG_szC (NL)** | 1.0876 | 0.0154 | 0.6616 | 2.2876 | 0.4427 | sz = 0.0042 | t05 = 2.4824; K = 0.3838 |

**Table S7** Neurally constrained Integration, Extremum-tracking and Extremum-flagging models with one extra feature at neural constraint weighting **w = 100**. ‘BIntg’ refers to the bounded Integration model, ‘BExtE’ refers to Extremum-tracking, and ‘BExtG’ refers to Extremum-flagging which has a guess default. ‘FlagWidth’ refers to the width of half-sine while t05 and K are semi-saturation constant and collapsing rate parameters of the collapsing bound function.

| **w = 100** | **G^2^+Penalty** | **B** | **D1** | **D2** | **FlagWidth** | **ExtraParam** | **t05, K** |
| --- | --- | --- | --- | --- | --- | --- | --- |
| **BIntg** | 33.9233 | 0.3385 | 0.0562 | 1.0881 | 0.7442 | - | - |
| **BIntg_eta** | 32.1992 | 0.5465 | 0.1313 | 1.8076 | 0.7777 | eta = 0.2423 | - |
| **BIntg_sz** | 20.89 | 0.2986 | 0.1092 | 1.3247 | 0.7319 | sz = 0.6 | - |
| **BIntg_Dur** | 19.7169 | 0.3156 | 0.0596 | 0.9471 | 0.6368 | Dur = 0.7337 | - |
| **BIntg_sampT** | 33.8987 | 0.3528 | 0.0562 | 1.106 | 0.7012 | sampT =-0.0042 | - |
| **BIntg_C** | 13.7526 | 0.3655 | 0.0564 | 0.9717 | 0.692 | - | K = 0.4606 |
| **BIntg_szC** | 13.0046 | 0.5084 | 0.0796 | 0.9872 | 0.229 | sz = 0.3883 | K = 0.4354 |
| **BIntg_szC (NL)** | 13.0225 | 0.4801 | 0.0765 | 0.9585 | 0.3287 | sz = 0.3506 | t05 = 3.9994; K = 2.2817 |
| **BIntg_Leak** | 24.0384 | 0.2833 | 0.0623 | 0.9387 | 0.657 | Leak = 0.0011 | - |
| **BExtE** | 152.579 | 0.035 | 0.3736 | 10.9167 | 0.9 | - | - |
| **BExtE_szC** | 132.6948 | 0.04 | 0.374 | 12.2806 | 0.8998 | sz = 0 | K=0.3293 |
| **BExtE_szC (NL)** | 133.3623 | 0.04 | 0.3768 | 12.1414 | 0.9 | - | t05 = 2.4993; K = 0.9923 |
| **BExtG** | 54.2312 | 0.0142 | 0.4184 | 12.92 | 0.717 | - | - |
| **BExtG_eta** | 54.9588 | 0.0142 | 0.414 | 15.559 | 0.7245 | eta = 0.0005 | - |
| **BExtG_sz** | 54.6168 | 0.0142 | 0.4162 | 15.6643 | 0.7175 | sz = 0.0003 | - |
| **BExtG_Dur** | 54.1953 | 0.0141 | 0.3999 | 14.2966 | 0.725 | Dur = 1.6 | - |
| **BExtG_sampT** | 50.5462 | 0.0141 | 0.38 | 12.1925 | 0.728 | sampT=-0.0064 | - |
| **BExtG_C** | 11.5902 | 0.015 | 0.4419 | 2.3786 | 0.4679 | - | K=0.1062 |
| **BExtG_szC** | 11.6206 | 0.015 | 0.4404 | 2.4433 | 0.4756 | sz = 0.00013 | K = 0.1079 |
| **BExtG_szC (NL)** | 12.3083 | 0.0152 | 0.4475 | 2.3865 | 0.4519 | sz = 0.0003 | t05 = 2.478; K= 0.4045 |

**Table S8** Neurally constrained Integration, Extremum-tracking and Extremum-flagging models with one extra feature at neural constraint weighting **w = 1000**. ‘BIntg’ refers to the bounded Integration model, ‘BExtE’ refers to Extremum-tracking, and ‘BExtG’ refers to Extremum-flagging which has a guess default. ‘FlagWidth’ refers to the width of half-sine while t05 and K are semi-saturation constant and collapsing rate parameters of the collapsing bound function.

| **w = 1000** | **G^2^+Penalty** | **B** | **D1** | **D2** | **FlagWidth** | **ExtraParam** | **t05, K** |
| --- | --- | --- | --- | --- | --- | --- | --- |
| **BIntg** | 290.8717 | 0.34 | 0.051 | 1.0718 | 0.7162 | - | - |
| **BIntg_eta** | 210.0866 | 0.6204 | 0.1483 | 1.9997 | 0.7719 | eta = 0.3123 | - |
| **BIntg_sz** | 203.2793 | 0.3004 | 0.1112 | 1.3314 | 0.744 | sz = 0.599 | - |
| **BIntg_Dur** | 109.6498 | 0.3125 | 0.0714 | 0.9528 | 0.656 | Dur =0.6722 | - |
| **BIntg_sampT** | 292.6721 | 0.343 | 0.0449 | 1.0551 | 0.6858 | sampT =-0.0116 | - |
| **BIntg_C** | 91.2485 | 0.3647 | 0.0605 | 0.9583 | 0.684 | - | K= 0.4545 |
| **BIntg_szC** | 82.3693 | 0.5031 | 0.0935 | 1.0116 | 0.2729 | sz =0.3937 | K =0.4356 |
| **BIntg_szC (NL)** | 82.1676 | 0.5654 | 0.0972 | 1.0175 | 0.2001 | sz = 0.4247 | t05 = 3.9479; K = 2.3504 |
| **BIntg_Leak** | 164.8869 | 0.2772 | 0.0813 | 0.9063 | 0.592 | Leak =0.0016 | - |
| **BExtE** | 1476.2579 | 0.035 | 0.2877 | 10.8605 | 0.9 | - | - |
| **BExtE_szC** | 1277.5765 | 0.04 | 0.3088 | 12.2639 | 0.9 | sz = 0.0001 | K=0.3379 |
| **BExtE_szC(NL)** | 1284.3111 | 0.04 | 0.3077 | 12.3065 | 0.9 | sz = 0.0002 | t05 =2.4994 ; K = 0.9263 |
| **BExtG** | 446.1987 | 0.014 | 0.1908 | 15.9076 | 0.714 | - | - |
| **BExtG_eta** | 450.3286 | 0.014 | 0.1858 | 13.8723 | 0.7184 | eta = 0.0057 | - |
| **BExtG_sz** | 446.7493 | 0.0141 | 0.1755 | 15.2241 | 0.7177 | sz = 0.0003 | - |
| **BExtG_Dur** | 448.168 | 0.0141 | 0.1951 | 14.3656 | 0.716 | Dur = 1.5763 | - |
| **BExtG_sampT** | 444.2898 | 0.014 | 0.2284 | 8.5694 | 0.7166 | sampT=-0.0014 | - |
| **BExtG_C** | 91.0628 | 0.0149 | 0.3951 | 2.4963 | 0.4953 | - | K=0.105 |
| **BExtG_szC** | 93.3027 | 0.015 | 0.4047 | 2.5057 | 0.4923 | sz = 0.00034 | K = 0.1042 |
| **BExtG_szC (NL)** | 100.3675 | 0.0152 | 0.464 | 2.4708 | 0.4674 | sz = 0.0002 | t05 = 2.4869; K= 0.3973; |
